## Supplementary material for "Capsular specificity in temperate phages of *Klebsiella pneumoniae* is driven by diverse receptor-binding enzymes": S1 Text

### Supplementary Text S1:
### GWAS-based depolymerase predictions

#### Methodology

The purpose of this study was to find capsule-specific phage depolymerases (i.e., those with a characteristic right-handed β-helix fold) by searching for associations between 35 most frequent K-loci in our dataset of *K. pneumoniae* genomes and protein clusters (PCs) of prophages found in those genomes, using a genome-wide association study (GWAS) approach. However, our results demonstrate (a) that such depolymerases are not always the strongest predictions for a given K-locus (cf., Figure 3B), and (b) that more predictors are revealed as putative depolymerases when we lowered the filtering thresholds (cf., Figure 3C). We therefore developed a semi-supervised curation framework combining multiple quantitative and biological evaluation steps.

We ran the GWAS analysis using six different combinations of clustering parameters: coverage thresholds of 50% and 80%, and sequence identity thresholds of 0%, 50%, and 80%; altogether six combinations of coverage and identity with the 50/50 threshold referred to as 'default' and others as 'non-default'. For each parameter set and K-locus, we identified initial predictors, defined as PCs with statistically significant positive associations ($\beta > 0$, $p < 3\times10^{-5}$ after Bonferroni correction).

To evaluate the classification performance of these predictors for each K-locis, we calculated precision and recall in two complementary ways: at the level of individual bacterial isolates and at the level of sequence clusters (SCs), which represent bacterial lineages defined using PopPunk.

Calculations at the **isolate level** were based on the following definitions:

- **True Positive (TP)**: an isolate contains both the target K-locus and the PC protein.
- **False Positive (FP)**: an isolate contains PC protein but lacks the target K locus.
- **False Negative (FN)**: an isolate contains the target K-locus but lacks the PC protein.
- **True Negative (TN)**: neither present.

Calculations at the **SC level** were designed to account for clonal structure. Here, we considered whether any isolate within an SC had the relevant feature:

- **TP (SC)**: the SC contains at least one isolate with both the target K locus and PC protein.
- **FP (SC)**: no isolate in the SC carries the target K locus, but at least one has the PC protein.
- **FN (SC)**: at least one isolate in the SC has the K locus, but none contain the PC protein.

- **FP and FN (SC)**: the SC contains both the K locus and the PC protein, but never in the same isolate.

$$\text{Precision} = \frac{\text{TP}}{\text{TP} + \text{FP}}$$

$$\text{Recall} = \frac{\text{TP}}{\text{TP} + \text{FN}}$$

$$\text{Precision (SC)} = \frac{\text{TP (SC)}}{\text{TP (SC)} + \text{FP (SC)}}$$

$$\text{Recall (SC)} = \frac{\text{TP (SC)}}{\text{TP (SC)} + \text{FN (SC)}}$$

This SC-level calculation helps down-weight associations that may be driven by clonal expansions of individual lineages, providing a more balanced view of lineage-level associations. However, it also relies on assumptions that may not always hold—for example, that the presence or absence of a K locus or protein within an SC is broadly representative of the entire cluster. In reality, some SCs may harbour substantial intra-lineage diversity or contain recent recombination events that break such associations. Since our dataset is ecologically diverse and representative, we used both isolate-level and SC-level precision and recall values to identify putative depolymerases using the GWAS approach.

We next applied a moderate filtering step by retaining only predictors with the precision value of 0.5 (isolate or SC) obtained at any level of clustering, hypothesising that some of these predictors could relatively well explain the K-locus distribution in the data but their other predictive metrics could be low in value due to biological nature of these proteins (e.g., high modularity or diversity). The remaining predictors were then manually curated and visually assessed, and finally classified into two classes of predictors, depending on the results, as described below.

We examined results for each K-locus by studying how precision and recall values (on both isolate and SC level) change between default clustering level and other clustering levels of all pectin lyase-like PCs (i.e., those for which we detected the presence of ECOD's pectin lyase-like domain, T-level:207.2.1). Importantly, we also tracked how protein clusters merged or split across clustering levels, ensuring that predictors reflect biologically meaningful sequence groups rather than technical artifacts of clustering. A non-default clustering threshold (other than 50/50) was chosen if it:

- (i) yielded high-precision SC-level PCs annotated as pectin lyase-like that were absent at default thresholds, or

- (ii) outperformed default clustering for pectin lyase-like PCs (i.e., higher precision with comparable recall, or comparable precision with higher recall).

Having selected the optimal clustering threshold for each K-locus, we next manually evaluated the shortlisted pectin lyase-like PCs by mapping their distribution onto the bacterial phylogeny alongside K-locus presence/absence data using Phandango. This phylogenetic mapping provided an additional layer of biological validation, helping to exclude associations driven by indirect linkage or recent HGT events. The bacterial tree was subsampled for visual clarity by downsampling a random subset of bacterial isolates without the K-locus of interest and without the examined PCs.

Candidate predictors were then classified into three categories:

- **'Strong'**: PC contained ≥10 protein sequences and was almost exclusively found in isolates with the target K-locus across the tree.
- **'Likely'**: PC contained <10 sequences, or occasional presence outside the target K-locus.
- **'Weak'**: no clear K-locus association; these were discarded.

For each candidate predictor, we selected a representative sequence by analyzing multiple sequence alignments and typically choosing the most frequent variant within the putative receptor-binding domain. Finally, all remaining predictors that passed the precision filtering but had no functional hit or had a functional hit to the alanine racemase-c domain (hits to N-terminal; ECOD T-level: 1.1.7) were examined if they were false-negatives. We found only one case (KL10, see Figure 3) where a PC had only a hit to alanine racemase-c fold where the presence of the right-handed β-helix fold was revealed via structural modelling with AlphaFold3.

Altogether, using this multi-step approach—combining statistical association, clustering optimization, multi-level precision/recall assessment, and phylogenetic mapping—we identified 12 strong predictors and 14 likely predictors, as shown in detail in the Figures below.

### Results

## KL3

In default clustering with 50% sequence identity and 50% coverage there are 3 PCs (PC0288, PC0449, PC0537) with pectin lyase-like fold, where 2 PCs have high precision on isolates and SCs and 1 PC has moderate precision calculated on isolates and SCs (see Figure 1A).

Increasing sequence identity to 80% leads to split of 2 PCs having high precision into 3 PCs with lower precision and lower recall calculated on isolates and SCs. Increase in sequence coverage to 80% leads to loss of 2 PCs, 1 PC with moderate precision and 1 PC with high precision calculated on isolates and SCs.

Therefore, we have chosen PC0288, PC0449, PC0537 from the clustering level with 50% sequence identity and 50% sequence coverage, because no other clustering level offers PCs with similar protein content to PCs with pectin lyase-like fold from default clustering with higher precision calculated on SCs or comparable precision and higher recall calculated on SCs, nor new PCs with pectin lyase-like fold and high precision calculated on SCs.

Further visualisation of these three clusters against the subsampled bacterial tree (see Figure 1B) shows that PC0449 and PC0537 are classified as 'strong' since both contain 10 or more protein sequences and were exclusively found in isolates with KL3 with confidence good or higher in multiple, distant bacterial lineages. PC0288 is classified as 'weak' since around 1/3 of protein sequences from this cluster were found in isolates having different K-locus or KL3 with confidence low or undetermined.

## KL7

In default clustering with 50% sequence identity and 50% sequence coverage there is 1 PC (PC0285) with pectin lyase-like fold, which has moderate precision and high recall calculated on SCs (see Figure 2A).

Increasing sequence identity and/or sequence coverage to 80% leads to increase in precision and decrease in recall calculated on SCs.

Both alternative clustering levels offer near-identical increase in precision and decrease in recall calculated on SCs, thus we have *a priori* chosen the PC0551 from clustering level with 50% sequence identity and 80% sequence coverage.

Further visualization of this cluster against the subsampled bacterial tree (see Figure 2B) shows that PC0551 is classified as 'likely' since it contains more than 10 protein sequences and is found nearly-exclusively in bacterial isolates having KL7 with confidence good or higher in multiple, distant bacterial lineages.

## KL10

In default clustering with 50% sequence identity and 50% sequence coverage there is 1 PC with alanine racemase-C fold (PC0844) (see Figure 3A). We confirmed the presence of pectin lyase-like fold in this PC using AlphaFold3 structural modelling and analyzed it further.

Increase in sequence identity and/or sequence coverage to 80% offers small changes in values of precision and recall calculated on isolates and SCs.

Therefore, we have chosen PC0844 from clustering with 50% sequence identity and 50% sequence coverage, because because no other clustering level offers PCs with similar protein content to PCs with alanine racemase-c fold from default clustering with higher precision calculated on SCs or comparable precision and higher recall calculated on SCs, nor new PCs with pectin lyase-like or alanine racemase-c folds and high precision calculated on isolates and SCs.

Further visualization of the associated cluster against the subsampled bacterial tree (see Figure 3B) shows PC0844 is classified as ‘strong’ since it contains more than 10 protein sequences and is found exclusively in bacterial isolates having KL10 with confidence good or better in multiple, distant bacterial lineages.

## KL14

In default clustering with 50% sequence identity and 50% sequence coverage there are 3 PCs (PC0367, PC1103, PC1304) with pectin lyase-like fold, where 1 PC has high precision and moderate recall calculated on SCs, 1 PC has moderate precision and high recall calculated on SCs, 1 PC has moderate precision and low recall calculated on SCs (see Figure 4A).

Increasing the sequence identity and/or coverage threshold to 80% results in comparable precision and recall values calculated on isolates and SCs. However, raising the identity threshold to 80% leads to the loss of 1 PC with moderate precision and low recall calculated on isolates and SCs.

Therefore, we have chosen PC0367, PC1103 and PC1304 from default clustering with 50% sequence identity and 50% sequence coverage, because no other clustering level offers PCs with similar protein content to PCs with pectin lyase-like fold from default clustering with higher precision calculated on SCs or comparable precision and higher recall calculated on SCs, nor new PCs with pectin lyase-like fold and high precision calculated on SCs.

Further visualization of these 3 PCs against the subsampled bacterial phylogenetic tree (see Figure 4B) shows that PC0367 is classified as ‘strong’ since it contains 10 or more protein sequences and was near-exclusively found in isolates with KL14 with confidence good or higher in multiple, distant bacterial lineages. PC1103 is classified as ‘likely’ since it contains less than 10 protein sequences and was near-exclusively found in isolates with KL14 with confidence good or higher in multiple, distant bacterial lineages. PC1304, even though it has a high value of precision calculated on SCs, was classified as ‘weak’ since it has less than 5 protein sequences and the association is driven by a single bacterial lineage.

## KL15

In default clustering with 50% sequence identity and 50% sequence coverage there is 1 PC (PC1771) with pectin lyase-like fold, high precision and low recall calculated on isolates and SCs (see Figure 5A).

Increasing sequence identity and/or sequence coverage to 80% leads to comparable values of precision and recall calculated on isolates and SCs. However, raising the coverage threshold to 80% leads to new associated pectin lyase-like PC with low precision and recall calculated on isolates and SCs.

Therefore, we have chosen PC1771 with clustering 50% sequence identity and 50% sequence coverage, because no other clustering level offers PCs with similar protein content to PCs with pectin lyase-like fold from default clustering with higher precision calculated on SCs or comparable precision and higher recall calculated on SCs, nor new PCs with pectin lyase-like fold and high precision calculated on SCs.

Further visualization of the associated PC against the subsampled bacterial tree (see Figure 5B) shows that PC1771 is classified as 'likely' since it contains less than 10 protein sequences and is found exclusively in bacterial isolates having KL15 with confidence good or higher in 3 different bacterial lineages.

## KL22

In default clustering with 50% sequence identity and 50% sequence coverage there are no associated PCs with pectin lyase-like fold (see Figure 6A). Alternative clustering levels do not offer any PCs with pectin lyase-like fold, except clustering with 80% sequence identity and 50% sequence coverage with one PC (PC1712) having high precision and low recall calculated on isolates and SCs.

Therefore, we have chosen PC1712 from clustering 80% sequence identity and 50% sequence coverage, because it's the only PC with pectin lyase-like fold and high precision calculated on isolates and SCs from all clustering levels.

Further visualization of this cluster against the subsampled bacterial tree (see Figure 6B) shows that PC1712 is classified as 'likely' since it has been found exclusively in isolates with KL22 with confidence good or higher, has less than 10 protein sequences, and was found in 2 bacterial lineages.

## KL24

In default clustering with 50% sequence identity and 50% sequence coverage there are 2 PCs with pectin lyase-like fold (PC1193, PC0491) and 1 PC with alanine racemase-c fold (PC1831) (see Figure 7A), where 1 PC with pectin lyase-like fold has moderate value of precision and recall calculated on isolates and SCs and, the 2 remaining PCs, one with pectin lyase-like fold and second with alanine racemase-C fold have moderate precision and low recall calculated on isolates and SCs.

Using AlphaFold3 structure modelling we found that PC1831 does not contain pectin lyase-like fold, thus we reject this PC from list of putative prophage depolymerases.

Increasing sequence identity to 80% leads to loss of 1 PC with pectin lyase-like fold with higher precision calculated on isolates and SCs. Increasing coverage to 80% leads to increase in precision and recall calculated on isolates and SCs for 2 PCs with pectin lyase-like fold which are similar in protein content to 2 PCs with pectin lyase-like fold from default clustering level.

Therefore, we have chosen the PC0406 and PC1397 from clustering level with identity 50% and coverage 80%, because it offers increased precision and recall calculated on isolates and SCs in comparison to PCs with pectin lyase-fold from default clustering level.

Further visualization of these clusters against the subsampled bacterial tree (see Figure 7B) shows that PC0406 is classified as ‘likely’ since it has more than 10 protein sequences and has been found nearly-exclusively in isolates having KL24 with confidence good or higher in multiple, distant bacterial lineages. PC1391 is classified as ‘likely’ since it has exactly 5 protein sequences and was found nearly-exclusively in isolates having KL24 with confidence good or higher in 2 distant bacterial lineages.

## KL25

In default clustering with 50% sequence identity and 50% sequence coverage there are 3 PCs with pectin lyase-like fold (PC0259, PC0500, PC0291) (see Figure 8A), 1 PC having moderate precision and high recall calculated on isolates and SCs, 1 PC having high precision and moderate recall calculated on isolates and SCs, and 1 PC having low precision and low recall calculated on isolates and SCs.

Increasing sequence identity or sequence coverage to 80% leads to comparable precision and recall calculated on SCs for high precision PC, decrease in recall calculated on SCs for moderate precision PC, and changes of recall calculated on SCs for low precision PC, decrease for increased identity and increase for increased coverage.

Increasing sequence identity and sequence coverage to 80% leads to a split of 1PC with moderate precision and high recall calculated on isolates and SCs into 2 PCs with higher precision and lower recall calculated on SCs and increase in precision calculated on isolates and SCs for all remaining PCs with pectin lyase-like fold.

Therefore, we have chosen PC1605, PC0538, PC0533, PC0848 from clustering with 80% sequence identity and 80% sequence coverage, because it offers increase in precision calculated on SCs for all PCs with pectin lyase-like fold in comparison to default clustering.

Further visualisation of these clusters against the subsampled bacterial tree (see Figure 8B) shows that PC1605 is classified as ‘strong’ since it was exclusively found in isolates with KL25 with good or higher confidence in multiple bacterial lineages. PC0538 is classified as ‘likely’ since it has 10 or more sequences and was found nearly-exclusively in isolates with KL25 with confidence good or higher in multiple bacterial lineages. PC0533 and PC0848 are classified as ‘weak’ since both were found multiple times in isolates with different KL or with KL25 with confidence low or undetermined.

## KL28

In default clustering with 50% sequence identity and 50% sequence coverage there is 1 PC with pectin lyase-like fold (PC0279) (see Figure 9A) with high precision and high recall calculated on isolates and SCs.

Increasing identity to 80% offers 1 PC with pectin lyase-like fold with comparable values of precision and recall calculated on isolates and SCs.

Increasing coverage to 80% leads to split of the PC into 5 PCs, where all PCs have lower values of recall calculated on isolates and SCs, 2 PCs have higher values of precision calculated on isolates and SCs, and 3 PCs have lower values of precision calculated on isolates and SCs.

Therefore, we have chosen PC0279 from clustering with 50% sequence identity and 50% sequence coverage, because no other clustering level offers PCs with similar protein content to PCs with pectin lyase-like fold from default clustering with clearly higher precision calculated on SCs or comparable precision and higher recall calculated on SCs, nor new PCs with pectin lyase-like fold and high precision calculated on SCs.

Further visualization of this cluster against the subsampled bacterial tree (see Figure 9B) shows that PC0279 is classified as ‘strong’ since it has more than 10 protein sequences and it was found nearly-exclusively in isolates with KL28 with confidence good or higher in multiple, distant bacterial lineages.

## KL30

In default clustering with 50% sequence identity and 50% coverage there is 1 PC (PC0827) with pectin lyase-like fold, high precision and low recall calculated on isolates and SCs (see Figure 10A).

Increasing sequence identity and/or sequence coverage to 80% leads to comparable values of precision and recall calculated on isolates and SCs.

Therefore, we have chosen PC0827 with clustering 50% sequence identity and 50% sequence coverage, because no other clustering level offers PCs with similar protein content to PCs with pectin lyase-like fold from default clustering with higher precision calculated on SCs or comparable precision and higher recall calculated on SCs, nor new PCs with pectin lyase-like fold and high precision calculated on SCs.

Further visualization of the cluster against the subsampled bacterial tree (see Figure 10B) shows that PC0827 is classified as 'likely' since it contains less than 10 protein sequences, and is found exclusively in bacterial isolates having KL30 with confidence good or higher in 2 distant bacterial lineages.

## KL38

In default clustering with 50% sequence identity and 50% sequence coverage there is 1 PC (PC0677) with pectin lyase-like topology, high precision and moderate recall calculated on isolates and SCs (see Figure 11A). Remaining PCs with pectin lyase-like topology have low precision and low recall calculated on isolates and SCs.

Increasing sequence identity and/or sequence coverage to 80% leads to a comparable values of precision and recall calculated on isolates and SCs for the high precision PC, and loss of low precision PCs with pectin lyase-like fold.

Therefore, we have chosen PC0854 with clustering 80% sequence identity and 50% sequence coverage, because it offers increase in precision and recall for high precision PC with pectin lyase-like fold.

Further visualization of the cluster against the subsampled bacterial tree (see Figure 11B) shows that PC0854 is classified as 'strong' since it contains more than 10 protein sequences, is found near-exclusively in bacterial isolates having KL38 with confidence good or higher in multiple, distant bacterial lineages.

## KL47

In default clustering with 50% sequence identity and 50% coverage there is 1 PC (PC1728) with pectin lyase-like topology, high precision and moderate recall calculated on isolates and SCs (see Figure 12A).

Increasing sequence identity and/or sequence coverage to 80% leads to comparable values of precision and recall calculated on isolates and SCs.

Therefore, we have chosen PC1728 with clustering 50% sequence identity and 50% sequence coverage, because no other clustering level offers PCs with similar protein content to PCs with pectin lyase-like fold from default clustering with higher precision calculated on SCs or comparable precision and higher recall calculated on SCs, nor new PCs with pectin lyase-like fold and high precision calculated on SCs.

Further visualization of the cluster against the subsampled bacterial tree (see Figure 12B) shows that PC1728 is classified as ‘likely’ since it contains less than 10 protein sequences, and is found exclusively in bacterial isolates having KL47 with confidence good or higher in 3 distant bacterial lineages.

## KL60

In default clustering with 50% sequence identity and 50% sequence coverage there are 3 PCs with pectin lyase-like fold (PC0848, PC0821, PC0464) (see Figure 13A), where 2 PCs have moderate precision and recall calculated on isolates and SCs, 1 PC have low precision and low recall calculated on isolates and SCs.

Increasing coverage to 80% offers comparable changes of precision and recall calculated on isolates and SCs, whereas increasing identity to 80% offers increased precision and increased recall calculated on SCs for all PCs with pectin lyase-like fold.

Therefore, we have chosen PC0871, PC1241 and PC1286 from clustering with 80% sequence identity and 50% sequence coverage, because all pectin lyase-like PCs have higher values of precision and recall calculated on isolates and SCs.

Further visualisation of these clusters against the subsampled bacterial tree (see Figure 13B) shows that PC0871 is classified as ‘strong’ since has more than 10 protein sequences, was nearly-exclusively found in isolates with KL60 with good or better confidence across multiple, distant bacterial lineages. PC1241 and PC1286 are classified as ‘weak’ because both have less than 10 protein sequences and are frequently found in isolates with different KL or KL60 with confidence low or undetermined.

## KL62

In default clustering with 50% sequence identity and 50% sequence coverage there is 1 PC (PC0400) (see Figure 14A) with high precision and high recall calculated on SCs.

Increasing sequence identity to 80% leads to decrease in recall calculated on SCs. Increasing sequence coverage to 80% leads to split of the PC with pectin lyase-like fold with high precision and high recall from default clustering level into 3 PCs, where 2 PCs have lower recall and higher precision calculated on SCs and 1 PC has comparable precision and lower recall calculated on SCs.

Therefore, we have chosen PC0769, PC0711, PC0756 with clustering 50% sequence identity and 80% sequence coverage, because it offers PCs with pectin lyase-like fold similar in protein content to the PC with pectin lyase-like fold from default clustering, but with higher precision calculated on SCs.

Further visualisation of these clusters against the subsampled bacterial tree (see Figure 14B) shows that PC0769, PC0711, PC0756 are classified as ‘strong’ since all have more than 10 proteins sequences and are exclusively or nearly-exclusively found in isolates with KL62 with confidence good or higher in multiple different bacterial lineages. However, due to high similarity sequence similarity (96.36%) across the whole length of the sequence between representative sequences of PC0756 and PC0769 we have excluded the PC0756 from further analysis.

## KL64

In default clustering with 50% sequence identity and 50% sequence coverage there are 3 PCs with pectin lyase-like fold (PC1304, PC0772, PC1375) (see Figure 15A), where 3 PCs have high precision and low recall calculated on SCs.

Increasing sequence identity and/or sequence coverage to 80% leads to comparable changes in precision and recall calculated on isolates and SCs.

Therefore, we have chosen PC1304, PC0772, PC1375 with clustering 50% sequence identity and 50% sequence coverage, because no other clustering level offers PCs with similar protein content to PCs with pectin lyase-like fold from default clustering with higher precision calculated on SCs or comparable precision and higher recall calculated on SCs, nor new PCs with pectin lyase-like fold and high precision calculated on SCs.

Further visualization of the cluster against the subsampled bacterial tree (see Figure 15B) shows that PC0772 is classified as ‘strong’ since it contains more than 10 protein sequences and is found exclusively in bacterial isolates having KL64 with good or higher confidence in multiple distant lineages. PC1375 is classified as ‘likely’ since it contains less than 10 protein sequences and is found exclusively in bacterial isolates having KL64 with good or higher confidence in 3 different bacterial lineages. PC1304 is classified as ‘weak’ since it has less than 10 protein sequences and 1/2 of them is found in isolates with different K-locus or in KL64 with confidence low or undetermined.

## KL111

In default clustering with 50% sequence identity and 50% sequence coverage there are 3 PCs with pectin lyase-like fold (PC0183, PC0259, PC0579) (see Figure 16A) all having moderate precision calculated on SCs, where 1 PC has high recall calculated on SCs, 1 PC has moderate recall calculated on SCs, 1 PC has low recall calculated on SCs.

Increasing sequence identity to 80% leads to comparable changes in precision and recall calculated on SCs for PCs with pectin lyase-like fold. Increasing coverage to 80% leads to increase in precision calculated on SCs for 2 PCs with low and moderate recall calculated on SCs.

We have chosen PC0692, PC0663, PC0258 with clustering 50% sequence identity and 80% sequence coverage, because this alternative clustering level offers increased precision calculated on SCs for 2 PCs with pectin lyase-like fold in comparison to default clustering level.

Further visualisation of these three clusters against the subsampled bacterial tree (see Figure 16B) shows that PC0692 is classified as ‘strong’ since it was near-exclusively found in isolates with KL111 with good or higher confidence in multiple, distant bacterial lineages and had more than 10 protein sequences, where PC0663, PC0258 were classified as ‘weak’ since they have more than 10 protein sequences and around half of these proteins is found in isolates with different K-locus or with KL111 with confidence low or undetermined.

## KL122

In default clustering with 50% sequence identity and 50% sequence coverage there is 1 PC (PC1357) with pectin lyase-like fold, high precision and low recall calculated on isolates and SCs (see Figure 17A).

Increasing sequence identity to 80% leads to comparable values of precision and recall calculated on isolates and SCs for the high precision PC. Increasing sequence coverage to 80% leads to loss of high precision PC.

Therefore, we have chosen PC1357 with clustering 50% sequence identity and 50% sequence coverage, because no other clustering level offers PCs with similar protein content to PCs with pectin lyase-like fold from default clustering with higher precision calculated on SCs or comparable precision and higher recall calculated on SCs, nor new PCs with pectin lyase-like fold and high precision calculated on SCs.

Further visualization of the cluster against the subsampled bacterial tree (see Figure 17B) shows that PC1357 is classified as ‘likely’ since it contains less than 10 protein sequences, and is found exclusively in bacterial isolates having KL122 with confidence good or higher in 2 bacterial lineages.

## KL127

In default clustering with 50% sequence identity and 50% sequence coverage there are 3 PCs (PC0183, PC1569, PC0502) (see Figure 18A) with pectin lyase-like fold, where 1 PC have high precision and low recall calculated on SCs, 1 PC has low precision and high recall calculated SCs and 1 PC has low precision and low recall calculated on SCs.

Increasing sequence coverage to 80% leads to comparable values of precision and recall calculated on SCs. Increasing sequence identity to 80% leads to split of PC0183 into 2 PCs (PC0692, PC0959) with higher precision and lower recall calculated on SCs.

Therefore, we have chosen PC0692, PC0959, PC1803 with clustering 80% sequence identity and 50% sequence coverage, because it offers PCs with similar protein content to the PCs with pectin lyase-like fold from default clustering level with higher precision calculated on SCs.

Further visualization of the clusters against the subsampled bacterial tree (see Figure 18B) shows PC0692 is classified as ‘likely’ since it has more than 10 protein sequences, but is sometimes found in isolates with different K-locus or KL127 with low or undetermined confidence in multiple, distant bacterial lineages. PC0959 is classified as ‘likely’ since it is found exclusively in isolates with KL127 in 3 different bacterial lineages, but has less than 10 protein sequences. PC1803 is classified as ‘likely’ since it was found near-exclusively in isolates with KL127 with confidence good or better in 4 bacterial lineages and has less than 10 protein sequences. PC0551 is classified as ‘weak’ since it has more than 10 protein sequences, but around half of them is found in isolates with different KL or KL127 with confidence lower or undetermined.

### Figures

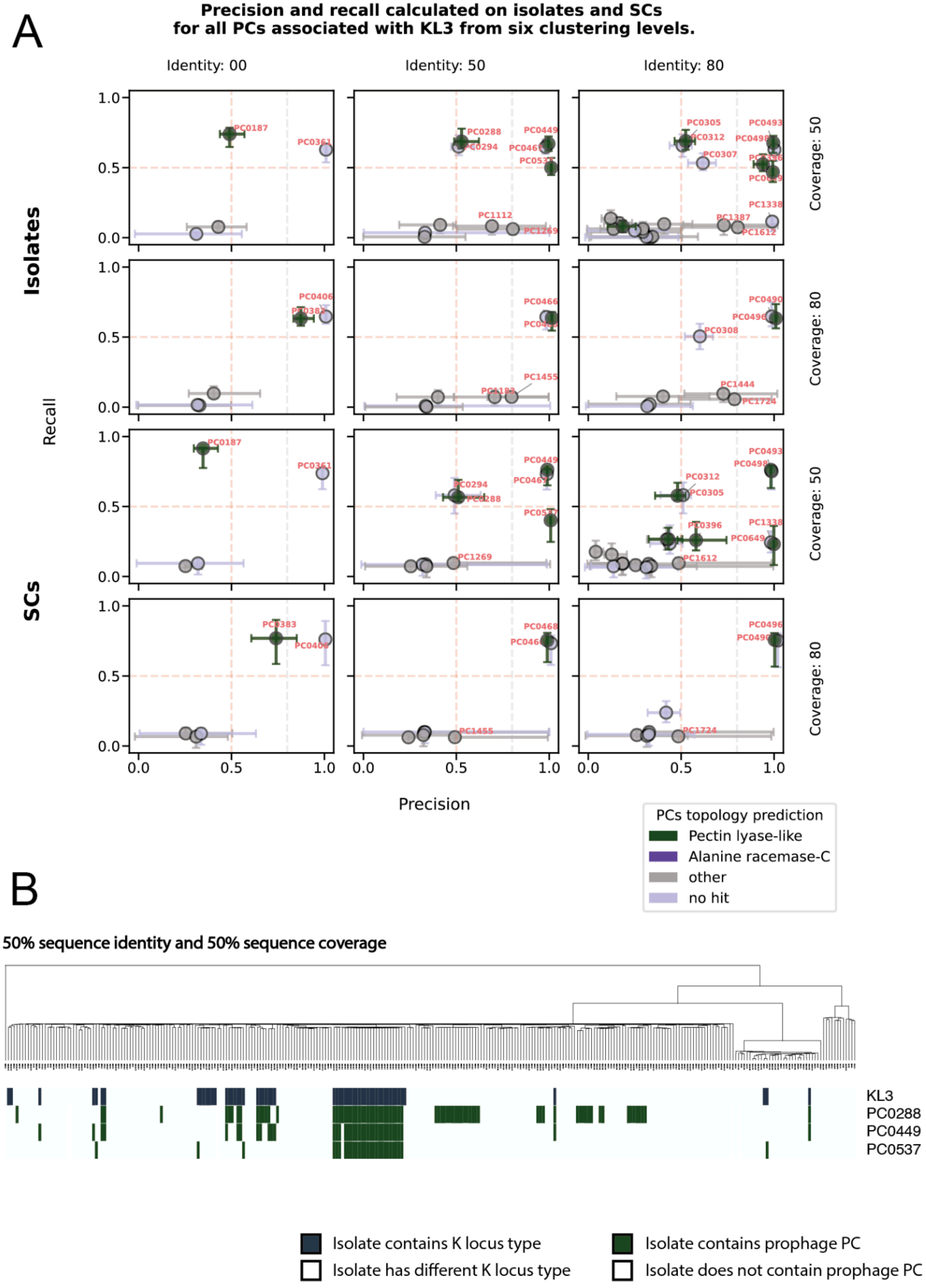
A
Precision and recall calculated on isolates and SCs
for all PCs associated with KL3 from six clustering levels.
Identity: 00
Identity: 50
Identity: 80
Coverage: 50
Coverage: 80
Isolates
SCs
Recall
Precision
PCs topology prediction
Pectin lyase-like
Alanine racemase-C
other
no hit
B
50% sequence identity and 50% sequence coverage
KL3
PC0288
PC0449
PC0537
Isolate contains K locus type
Isolate has different K locus type
Isolate contains prophage PC
Isolate does not contain prophage PC

**Figure 1.** Identification of KL3-specific prophage depolymerases using GWAS. (A) The figure shows precision and recall with CI 95% for GWAS-linked PCs across six sequence-clustering thresholds. Colours distinguish predicted ECOD folds: Pectin lyase-like, Alanine racemase-C, other ECOD folds or no similarity to ECOD database detected. (B) Distribution of KL3 and its associated PCs with pectin lyase-like fold at the 50 % identity and 50 % bidirectional coverage level. The bacterial phylogenetic tree has been sub-sampled, but only isolates lacking both KL3 and the PCs with pectin lyase-like fold were removed.

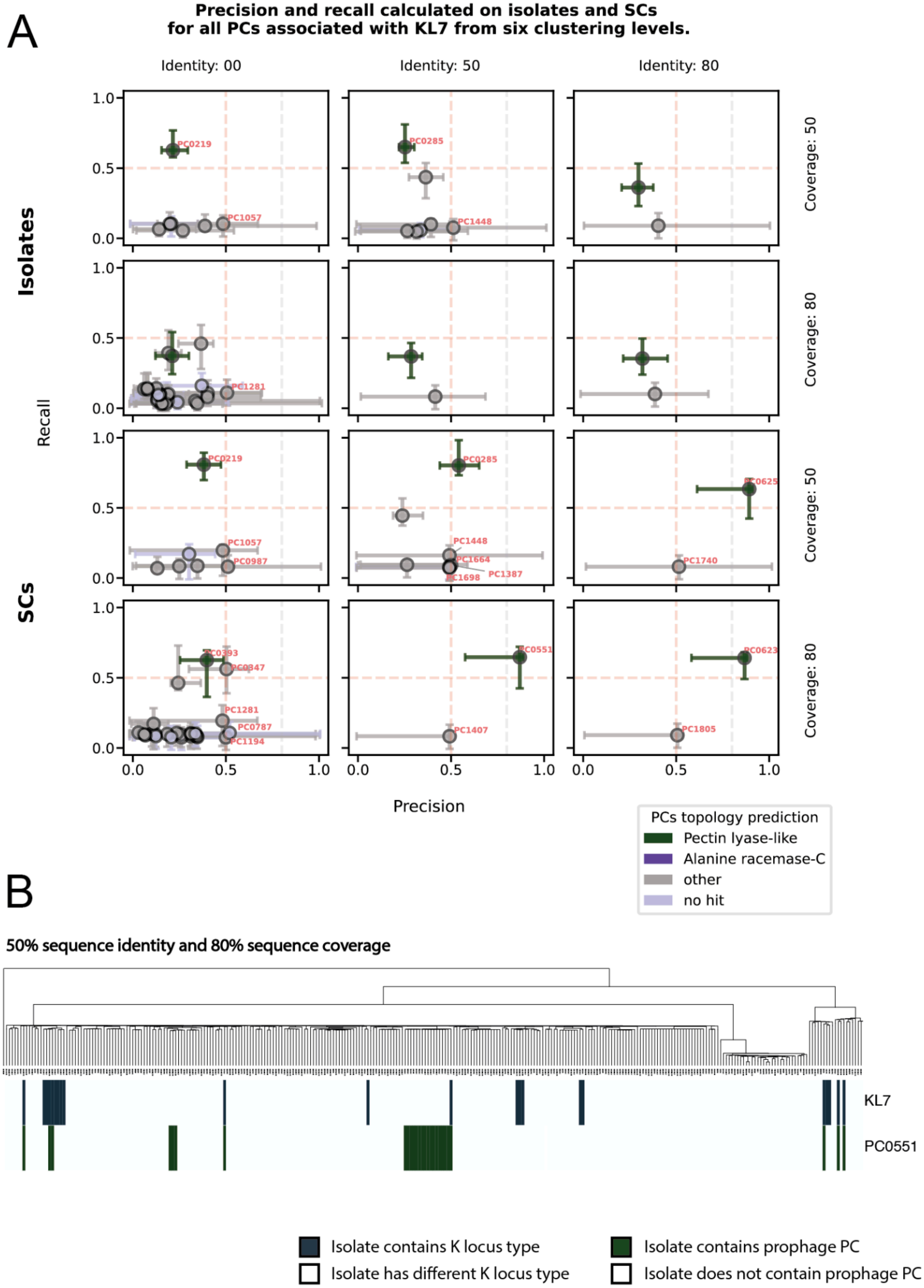

A
Precision and recall calculated on isolates and SCs
for all PCs associated with KL7 from six clustering levels.
Identity: 00
Identity: 50
Identity: 80
Coverage: 50
Coverage: 80
Coverage: 50
Coverage: 80
Isolates
SCs
Recall
Precision
0.0
0.5
1.0
PC0219
PC1057
PC0285
PC1448
PC1281
PC0987
PC1664
PC1387
PC1698
PC0625
PC1740
PC0393
PC0347
PC0787
PC1194
PC0551
PC1407
PC0623
PC1805
PCs topology prediction
Pectin lyase-like
Alanine racemase-C
other
no hit
B
50% sequence identity and 80% sequence coverage
KL7
PC0551
Isolate contains K locus type
Isolate has different K locus type
Isolate contains prophage PC
Isolate does not contain prophage PC

**Figure 2.** Identification of KL7-specific prophage depolymerases using GWAS. (A) The figure shows precision and recall with CI 95% for GWAS-linked PCs across six sequence-clustering thresholds. Colours distinguish predicted ECOD folds: Pectin lyase-like, Alanine racemase-C, other ECOD folds or no similarity to ECOD database detected. (B) Distribution of KL7 and associated PC with pectin lyase-like fold at the 50 % identity and 80 % bidirectional coverage level. The bacterial phylogenetic tree has been sub-sampled, but only isolates lacking both KL7 and the PCs with pectin lyase-like fold were removed.

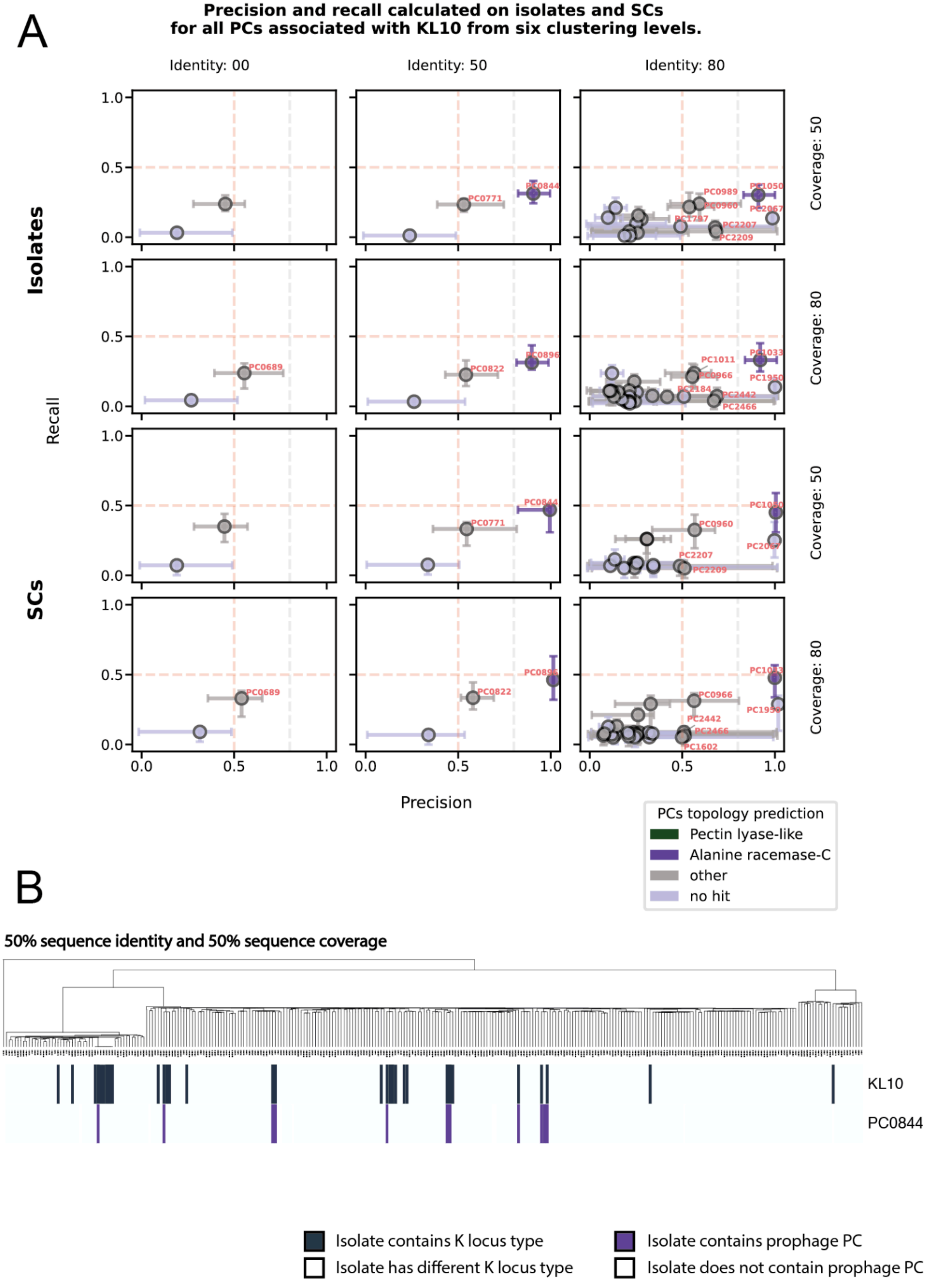
A
Precision and recall calculated on isolates and SCs
for all PCs associated with KL10 from six clustering levels.
Identity: 00
Identity: 50
Identity: 80
Coverage: 50
Coverage: 80
Coverage: 50
Coverage: 80
Isolates
SCs
Recall
Precision
0.0
0.5
1.0
PC0844
PC0771
PC0689
PC0822
PC0896
PCs topology prediction
Pectin lyase-like
Alanine racemase-C
other
no hit
B
50% sequence identity and 50% sequence coverage
KL10
PC0844
Isolate contains K locus type
Isolate has different K locus type
Isolate contains prophage PC
Isolate does not contain prophage PC

**Figure 3.** Identification of KL10-specific prophage depolymerases using GWAS. (A) The figure shows precision and recall with CI 95% for GWAS-linked PCs across six sequence-clustering thresholds. Colours distinguish predicted ECOD folds: Pectin-lyase-like, Alanine racemase-C, other ECOD folds or no similarity to ECOD database detected. (B) Distribution of KL10 and associated PC with Alanine racemase-C fold at the 50 % identity and 50 % bidirectional coverage level. The bacterial phylogenetic tree has been sub-sampled, but only isolates lacking both KL10 and the PCs with Alanine racemase-C fold were removed.

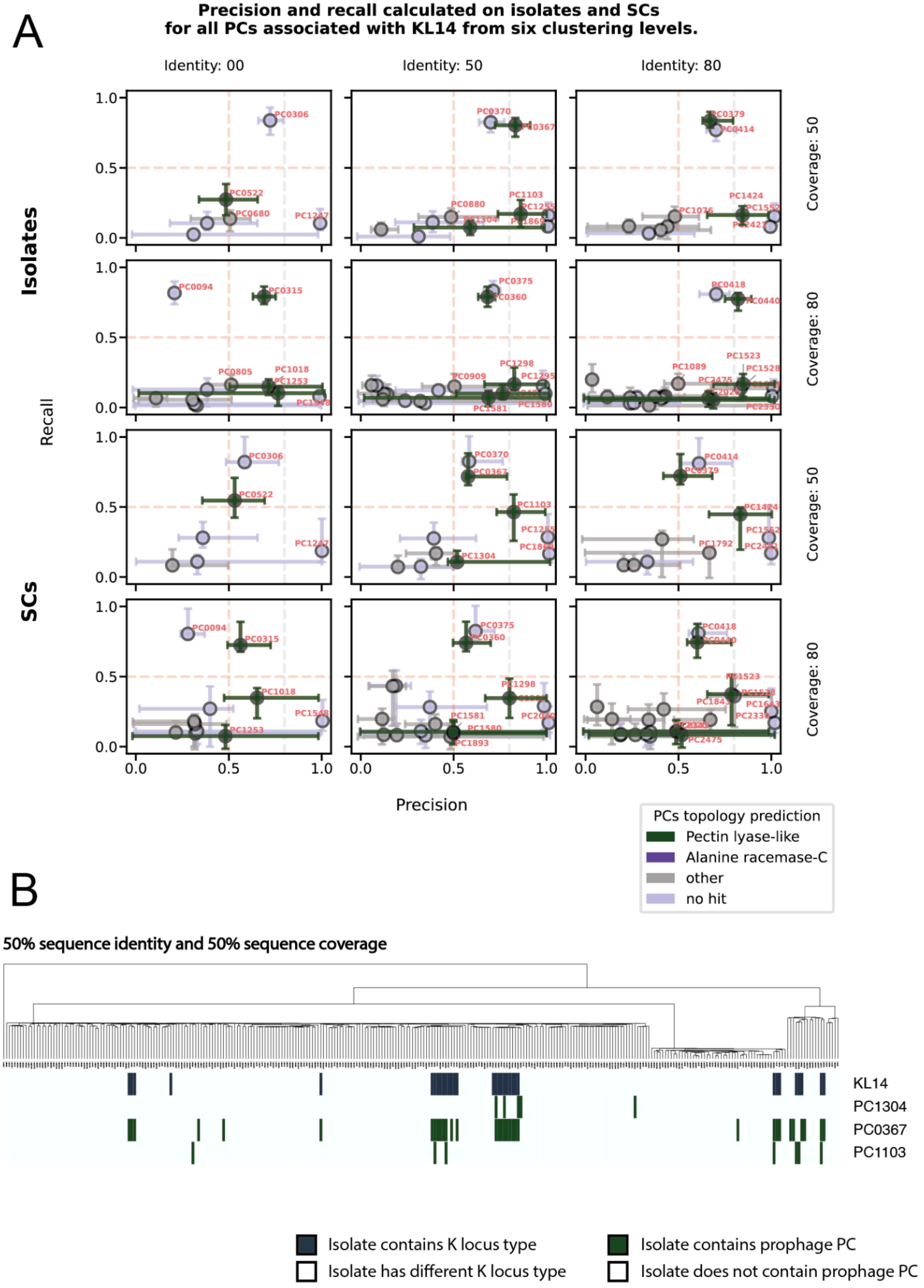
A
Precision and recall calculated on isolates and SCs
for all PCs associated with KL14 from six clustering levels.
Identity: 00
Identity: 50
Identity: 80
Coverage: 50
Coverage: 80
Isolates
SCs
Recall
Precision
PCs topology prediction
Pectin lyase-like
Alanine racemase-C
other
no hit
B
50% sequence identity and 50% sequence coverage
KL14
PC1304
PC0367
PC1103
Isolate contains K locus type
Isolate has different K locus type
Isolate contains prophage PC
Isolate does not contain prophage PC

**Figure 4.** Identification of KL14-specific prophage depolymerases using GWAS. (A) The figure shows precision and recall with CI 95% for GWAS-linked PCs across six sequence-clustering thresholds. Colours distinguish predicted ECOD folds: Pectin lyase-like, Alanine racemase-C, other ECOD folds or no similarity to ECOD database detected. (B) Distribution of KL14 and associated PC with pectin-lyase-like fold at the 50 % identity and 50 % bidirectional coverage level. The bacterial phylogenetic tree has been sub-sampled, but only isolates lacking both KL14 and the PCs with pectin lyase-like fold were removed.

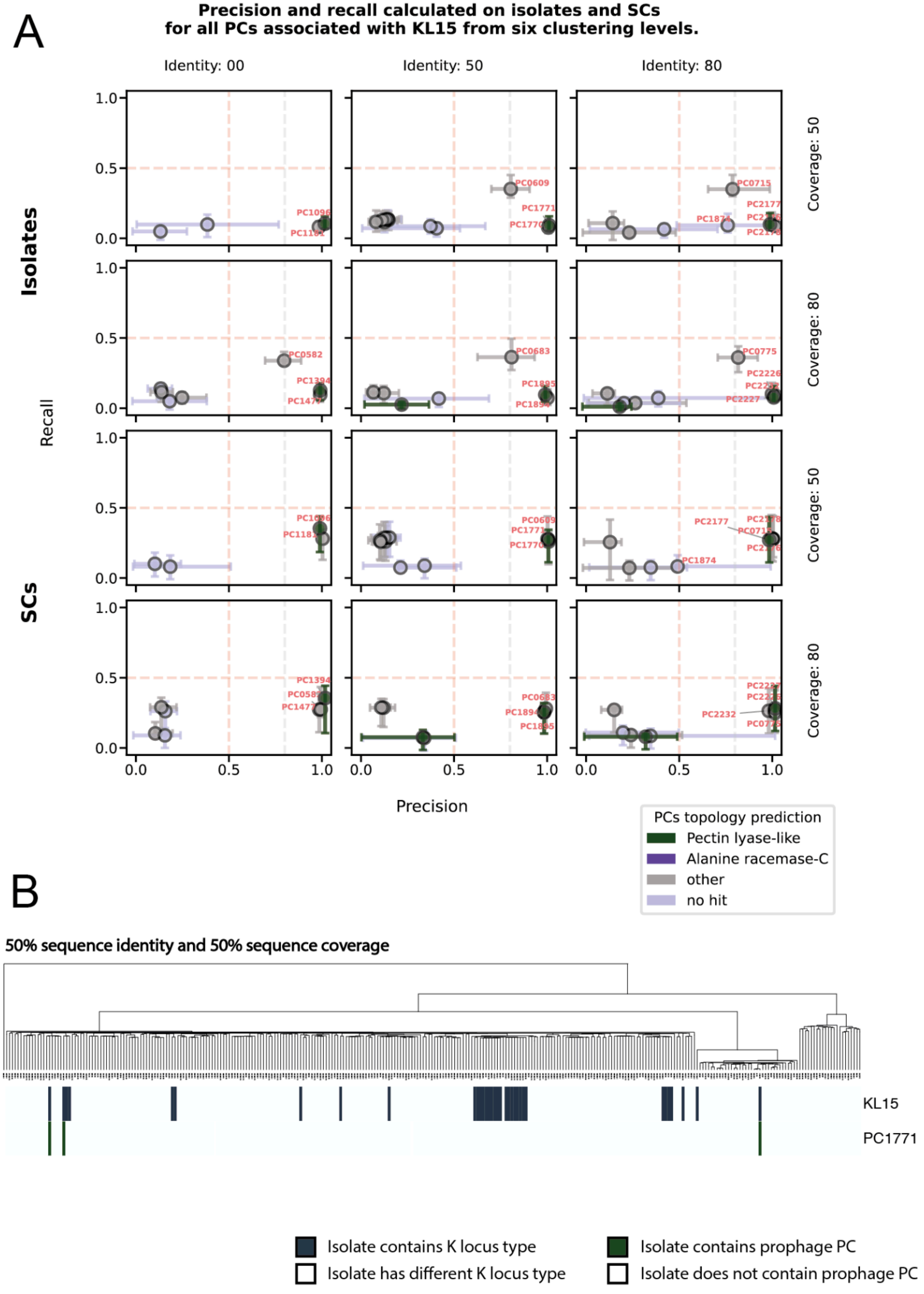
A
Precision and recall calculated on isolates and SCs
for all PCs associated with KL15 from six clustering levels.
Identity: 00
Identity: 50
Identity: 80
Coverage: 50
Coverage: 80
Coverage: 50
Coverage: 80
Isolates
SCs
Recall
Precision
PC0609
PC1771
PC0715
PC2177
PC0582
PC0683
PC0775
PC2226
PC2227
PC2177
PC1874
PC2232
PCs topology prediction
Pectin lyase-like
Alanine racemase-C
other
no hit
B
50% sequence identity and 50% sequence coverage
KL15
PC1771
Isolate contains K locus type
Isolate has different K locus type
Isolate contains prophage PC
Isolate does not contain prophage PC

**Figure 5.** Identification of KL15-specific prophage depolymerases using GWAS. (A) The figure shows precision and recall with CI 95% for GWAS-linked PCs across six sequence-clustering thresholds. Colours distinguish predicted ECOD folds: Pectin-lyase-like, Alanine racemase-C, other ECOD folds or no similarity to ECOD database detected. (B) Distribution of KL15 and associated PC with pectin lyase-like fold at the 50 % identity and 50 % bidirectional coverage level. The bacterial phylogenetic tree has been sub-sampled, but only isolates lacking both KL15 and the PCs with pectin lyase-like fold were removed.

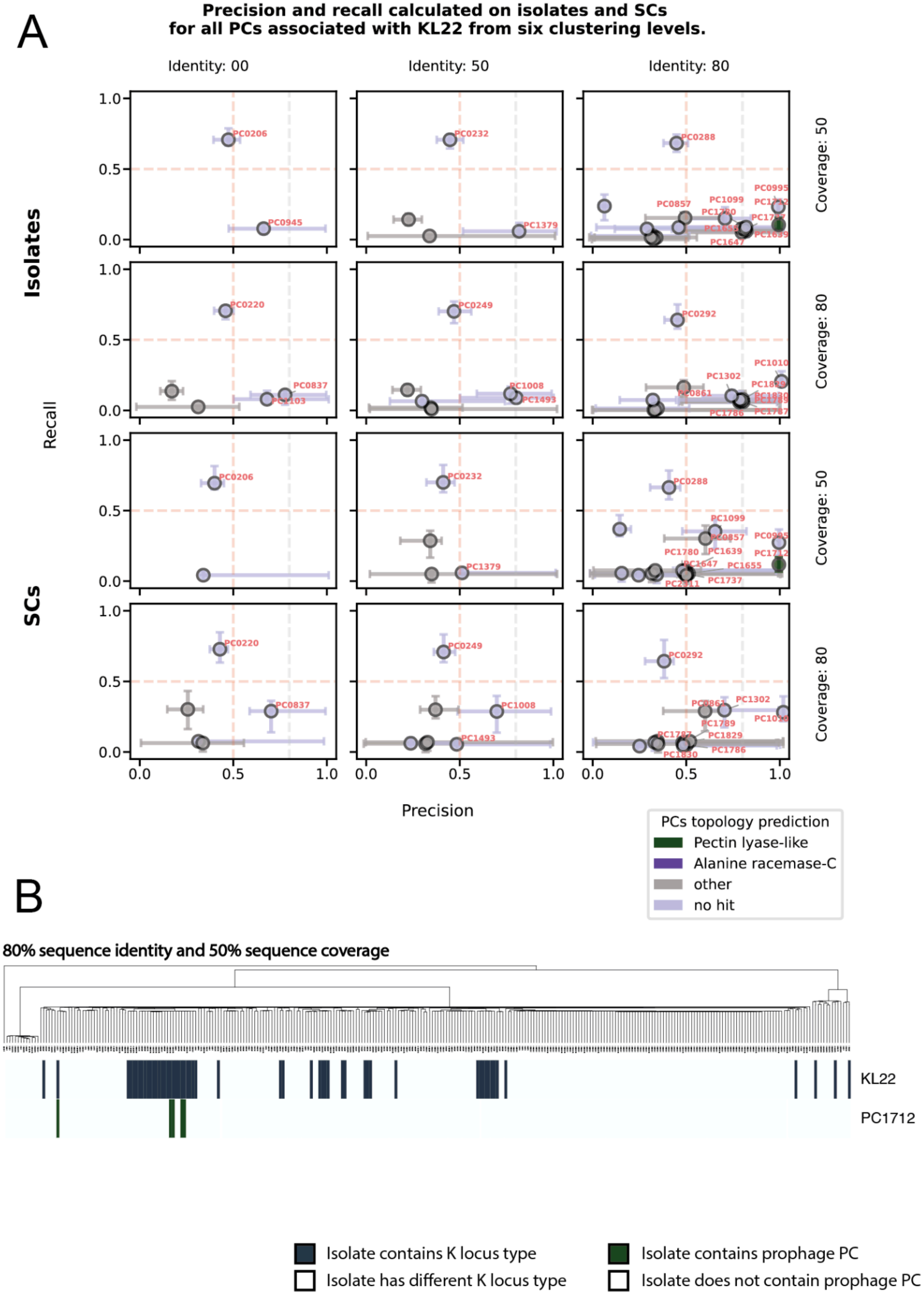
A
Precision and recall calculated on isolates and SCs
for all PCs associated with KL22 from six clustering levels.
Identity: 00
Identity: 50
Identity: 80
Coverage: 50
Coverage: 80
Isolates
SCs
Recall
Precision
PC0206
PC0945
PC0232
PC1379
PC0288
PC0220
PC0837
PC0249
PC1008
PC1493
PC0292
PC1099
PC0857
PC1302
PC1010
PC0861
PC1655
PC1737
PC1830
PC1829
PC1786
PC1789
PC1787
PC1780
PC1639
PC1647
PC1712
PC0995
PCs topology prediction
Pectin lyase-like
Alanine racemase-C
other
no hit
B
80% sequence identity and 50% sequence coverage
KL22
PC1712
Isolate contains K locus type
Isolate has different K locus type
Isolate contains prophage PC
Isolate does not contain prophage PC

**Figure 6.** Identification of KL22-specific prophage depolymerases using GWAS. (A) The figure shows precision and recall with CI 95% for GWAS-linked PCs across six sequence-clustering thresholds. Colours distinguish predicted ECOD folds: Pectin lyase-like, Alanine racemase-C, other ECOD folds or no similarity to ECOD database detected. (B) Distribution of KL22 and associated PC with pectin lyase-like fold at the 80 % identity and 50 % bidirectional coverage level. The bacterial phylogenetic tree has been sub-sampled, but only isolates lacking both KL22 and the PCs with pectin lyase-like fold were removed.

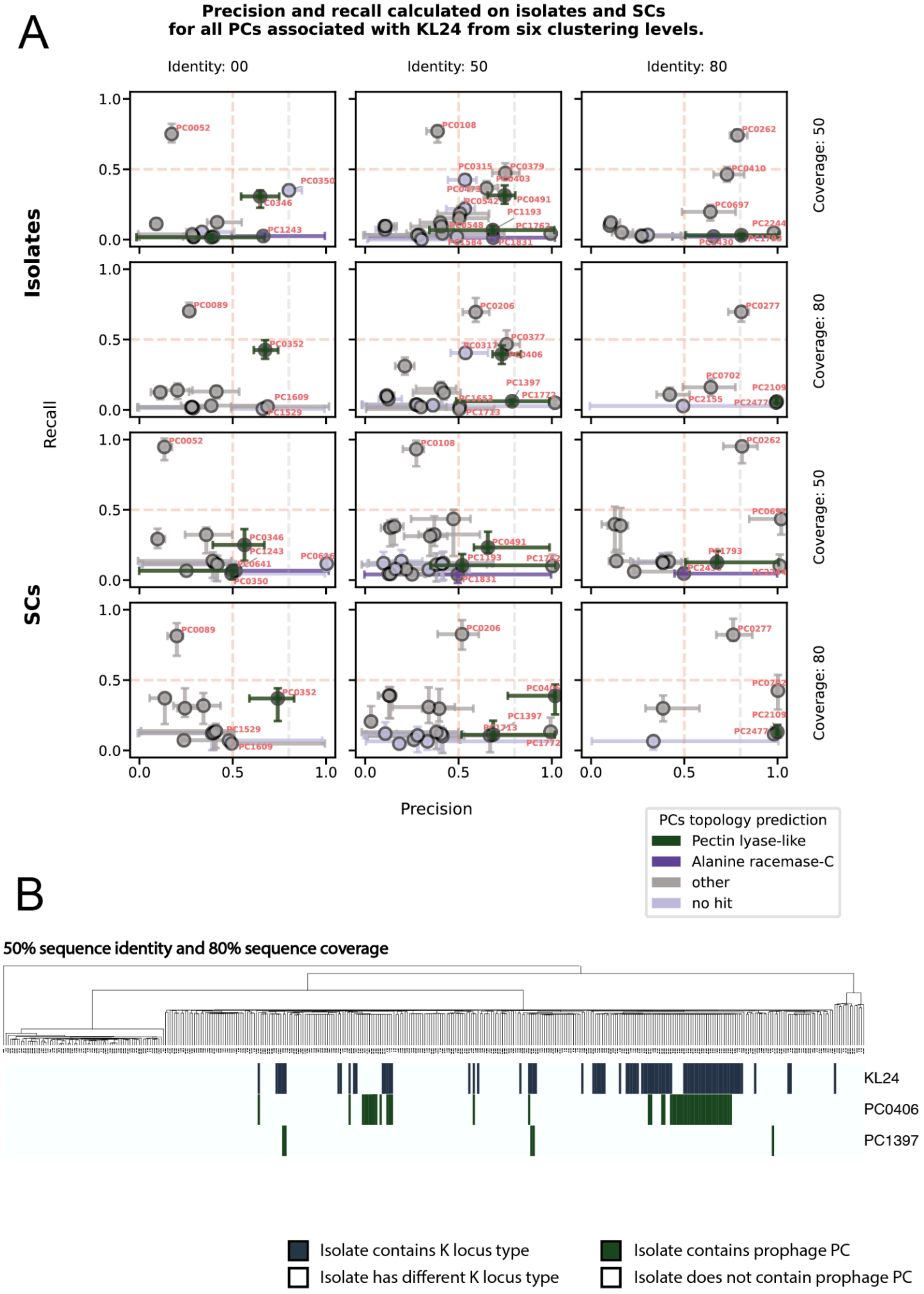
A
Precision and recall calculated on isolates and SCs
for all PCs associated with KL24 from six clustering levels.
Identity: 00
Identity: 50
Identity: 80
Coverage: 50
Coverage: 80
Coverage: 50
Coverage: 80
Isolates
SCs
Recall
Precision
0.0
0.5
1.0
PCs topology prediction
Pectin lyase-like
Alanine racemase-C
other
no hit
B
50% sequence identity and 80% sequence coverage
KL24
PC0406
PC1397
Isolate contains K locus type
Isolate has different K locus type
Isolate contains prophage PC
Isolate does not contain prophage PC

**Figure 7.** Identification of KL24-specific prophage depolymerases using GWAS. (A) The figure shows precision and recall with CI 95% for GWAS-linked PCs across six sequence-clustering thresholds. Colours distinguish predicted ECOD folds: Pectin lyase-like, Alanine racemase-C, other ECOD folds or no similarity to ECOD database detected. (B) Distribution of KL24 and associated PC with pectin lyase-like fold at the 50 % identity and 80 % bidirectional coverage level. The bacterial phylogenetic tree has been sub-sampled, but only isolates lacking both KL24 and the PCs with pectin lyase-like fold were removed.

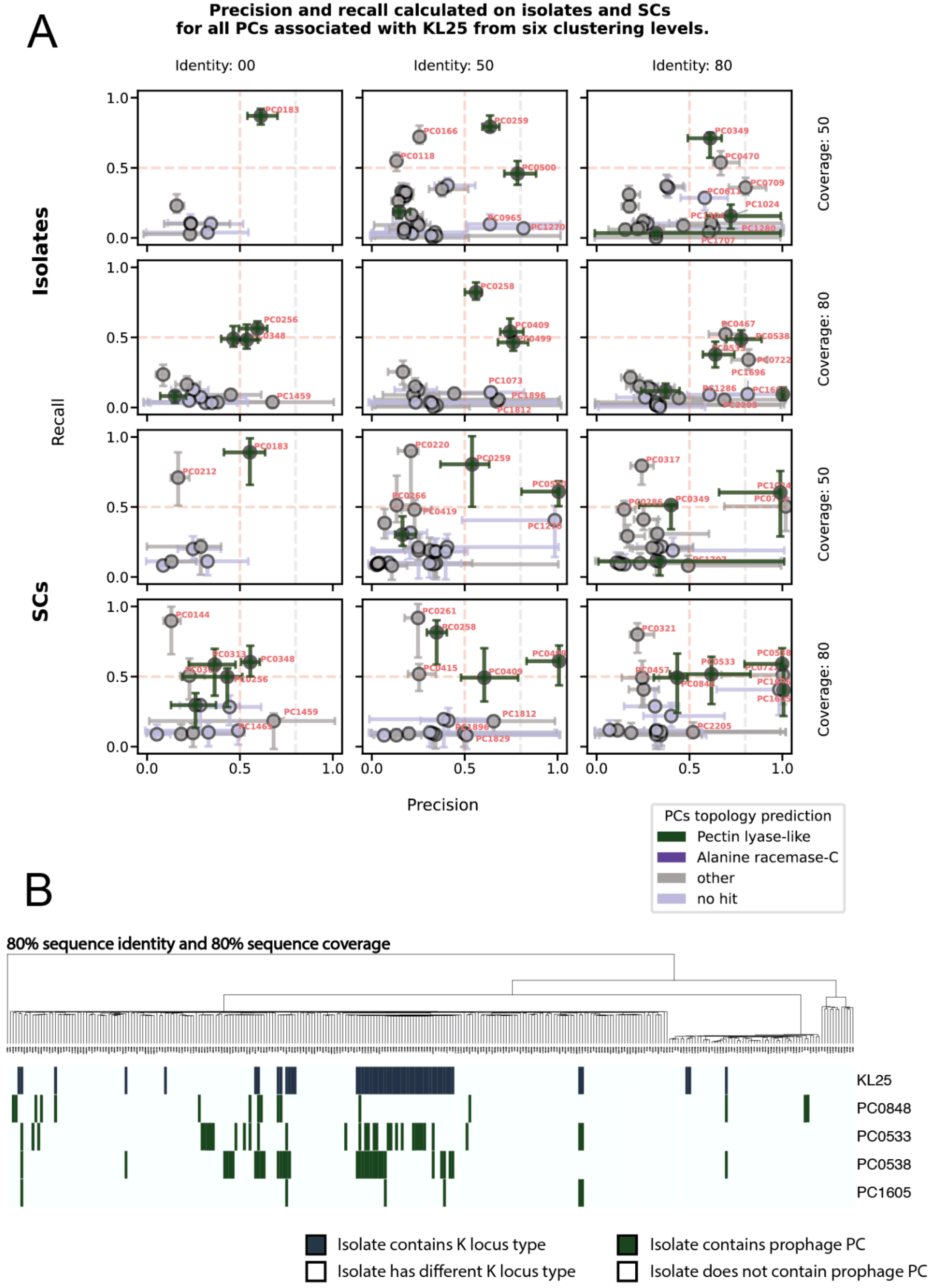
A
Precision and recall calculated on isolates and SCs
for all PCs associated with KL25 from six clustering levels.
Identity: 00
Identity: 50
Identity: 80
Coverage: 50
Coverage: 80
Isolates
SCs
Recall
Precision
PCs topology prediction
Pectin lyase-like
Alanine racemase-C
other
no hit
B
80% sequence identity and 80% sequence coverage
KL25
PC0848
PC0533
PC0538
PC1605
Isolate contains K locus type
Isolate has different K locus type
Isolate contains prophage PC
Isolate does not contain prophage PC

**Figure 8.** Identification of KL25-specific prophage depolymerases using GWAS. (A) The figure shows precision and recall with CI 95% for GWAS-linked PCs across six sequence-clustering thresholds. Colours distinguish predicted ECOD folds: Pectin lyase-like, Alanine racemase-C, other ECOD folds or no similarity to ECOD database detected. (B) Distribution of KL25 and associated PC with pectin lyase-like fold at the 80 % identity and 80 % bidirectional coverage level. The bacterial phylogenetic tree has been sub-sampled, but only isolates lacking both KL25 and the PCs with pectin lyase-like fold were removed.

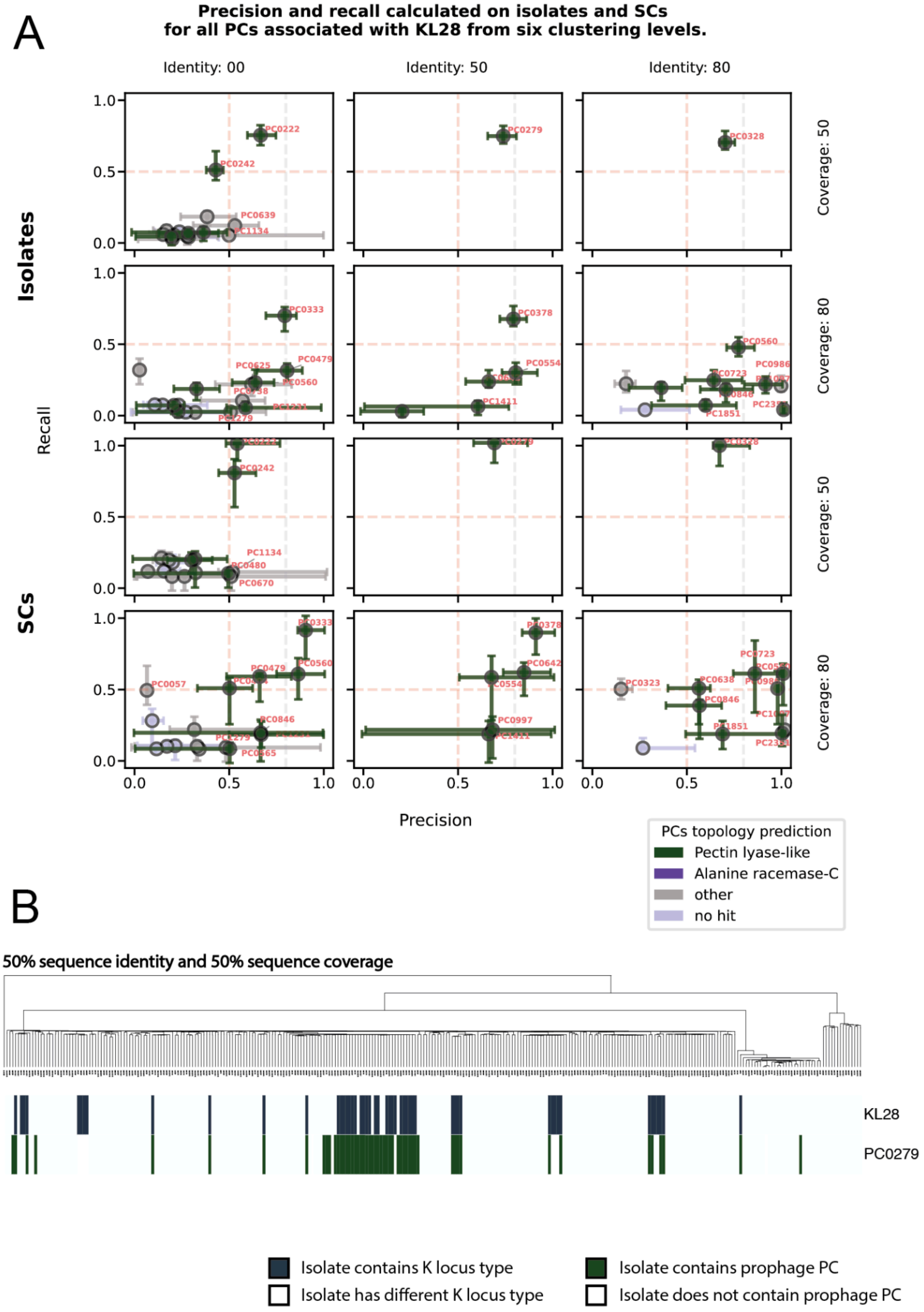
A
Precision and recall calculated on isolates and SCs
for all PCs associated with KL28 from six clustering levels.
Identity: 00
Identity: 50
Identity: 80
Coverage: 50
Coverage: 80
Coverage: 50
Coverage: 80
Isolates
SCs
Recall
Precision
PCs topology prediction
Pectin lyase-like
Alanine racemase-C
other
no hit
B
50% sequence identity and 50% sequence coverage
KL28
PC0279
Isolate contains K locus type
Isolate has different K locus type
Isolate contains prophage PC
Isolate does not contain prophage PC

**Figure 9.** Identification of KL28-specific prophage depolymerases using GWAS. (A) The figure shows precision and recall with CI 95% for GWAS-linked PCs across six sequence-clustering thresholds. Colours distinguish predicted ECOD folds: Pectin lyase-like, Alanine racemase-C, other ECOD folds or no similarity to ECOD database detected. (B) Distribution of KL28 and associated PC with pectin-lyase-like fold at the 50 % identity and 50 % bidirectional coverage level. The bacterial phylogenetic tree has been sub-sampled, but only isolates lacking both KL28 and the PCs with pectin lyase-like fold were removed.

A

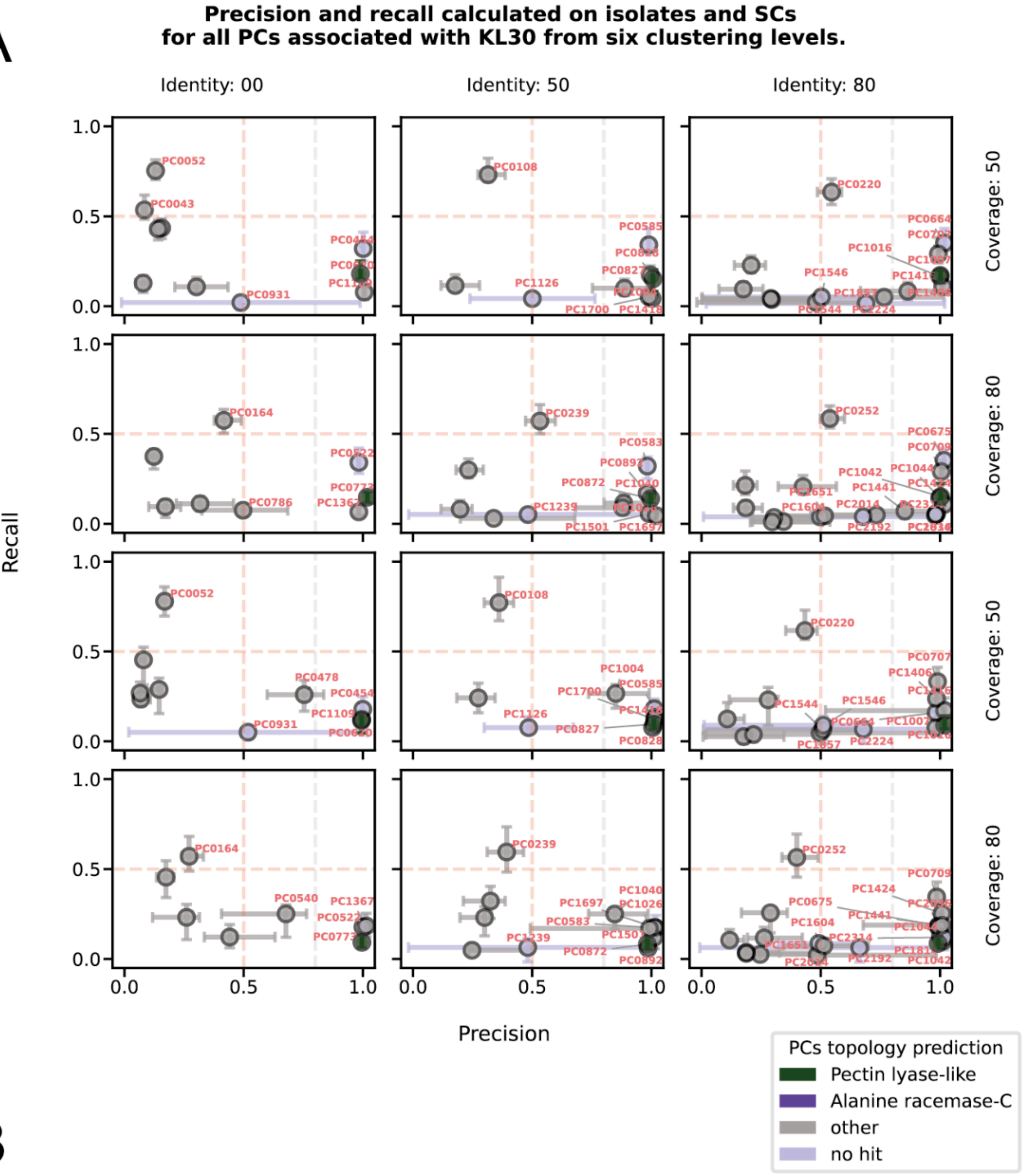

Precision and recall calculated on isolates and SCs
for all PCs associated with KL30 from six clustering levels.
Identity: 00
Identity: 50
Identity: 80
Coverage: 50
Coverage: 80
Isolates
SCs
Recall
Precision
PCs topology prediction
Pectin lyase-like
Alanine racemase-C
other
no hit


B

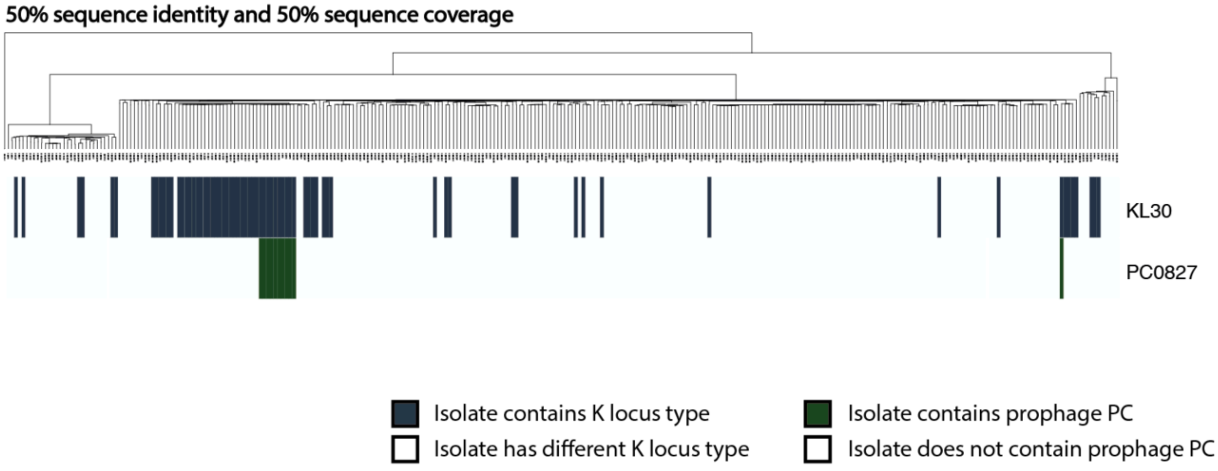

50% sequence identity and 50% sequence coverage
KL30
PC0827
Isolate contains K locus type
Isolate has different K locus type
Isolate contains prophage PC
Isolate does not contain prophage PC

**Figure 10.** Identification of KL30-specific prophage depolymerases using GWAS. (A) The figure shows precision and recall with CI 95% for GWAS-linked PCs across six sequence-clustering thresholds. Colours distinguish predicted ECOD folds: Pectin lyase-like, Alanine racemase-C, other ECOD folds or no similarity to ECOD database detected. (B) Distribution of KL30 and associated PC with pectin lyase-like fold at the 50 % identity and 50 % bidirectional coverage level. The bacterial phylogenetic tree has been sub-sampled, but only isolates lacking both KL30 and the PCs with pectin lyase-like fold were removed.

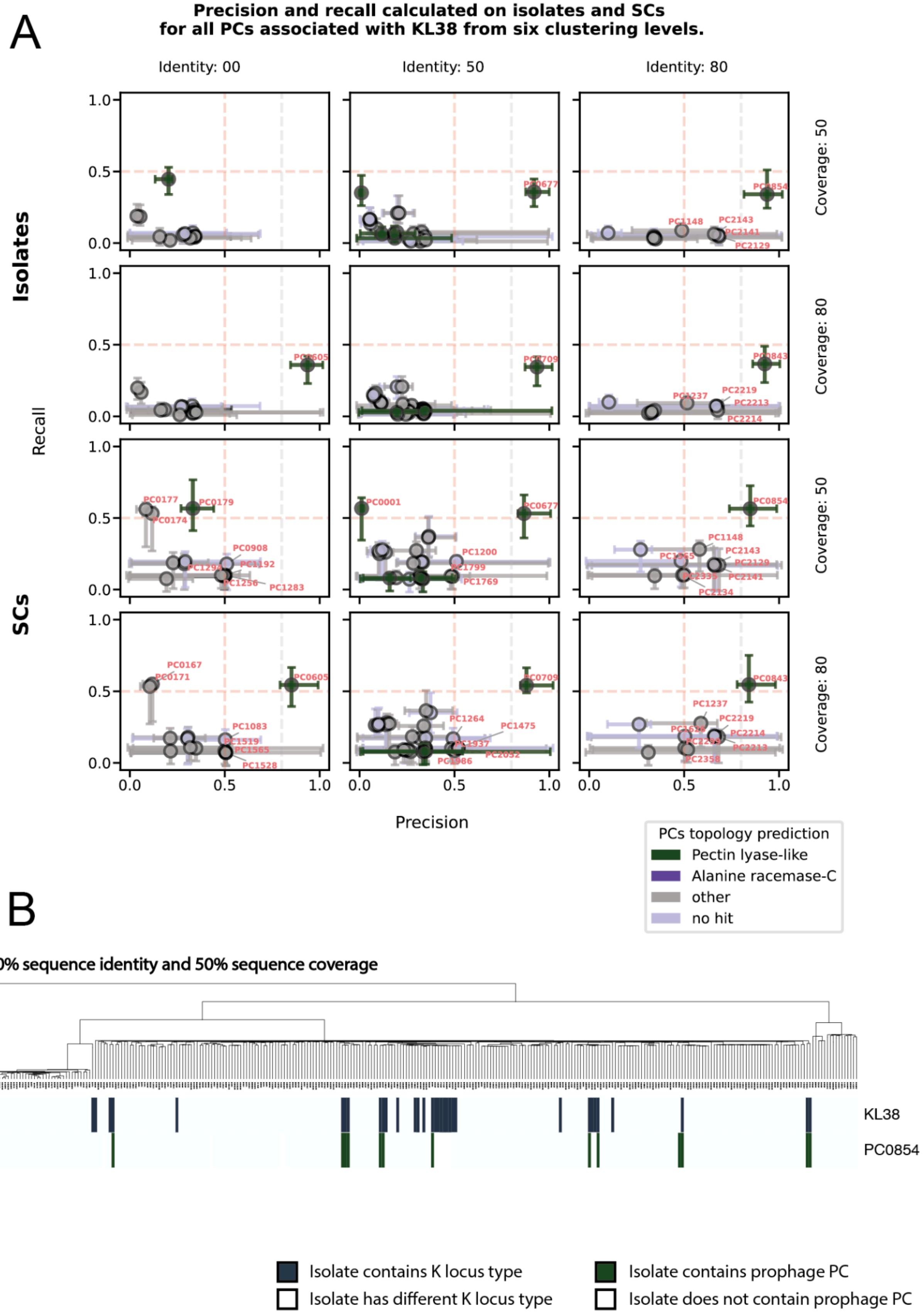

A
Precision and recall calculated on isolates and SCs
for all PCs associated with KL38 from six clustering levels.
Identity: 00
Identity: 50
Identity: 80
Coverage: 50
Coverage: 80
Coverage: 50
Coverage: 80
Isolates
SCs
Recall
Precision
PCs topology prediction
Pectin lyase-like
Alanine racemase-C
other
no hit
B
80% sequence identity and 50% sequence coverage
KL38
PC0854
Isolate contains K locus type
Isolate has different K locus type
Isolate contains prophage PC
Isolate does not contain prophage PC

**Figure 11.** Identification of KL38-specific prophage depolymerases using GWAS. (A) The figure shows precision and recall with CI 95% for GWAS-linked PCs across six sequence-clustering thresholds. Colours distinguish predicted ECOD folds: Pectin lyase-like, Alanine racemase-C, other ECOD folds or no similarity to ECOD database detected. (B) Distribution of KL38 and associated PC with pectin-lyase-like fold at the 80 % identity and 50 % bidirectional coverage level. The bacterial phylogenetic tree has been sub-sampled, but only isolates lacking both KL38 and the PCs with pectin lyase-like fold were removed.

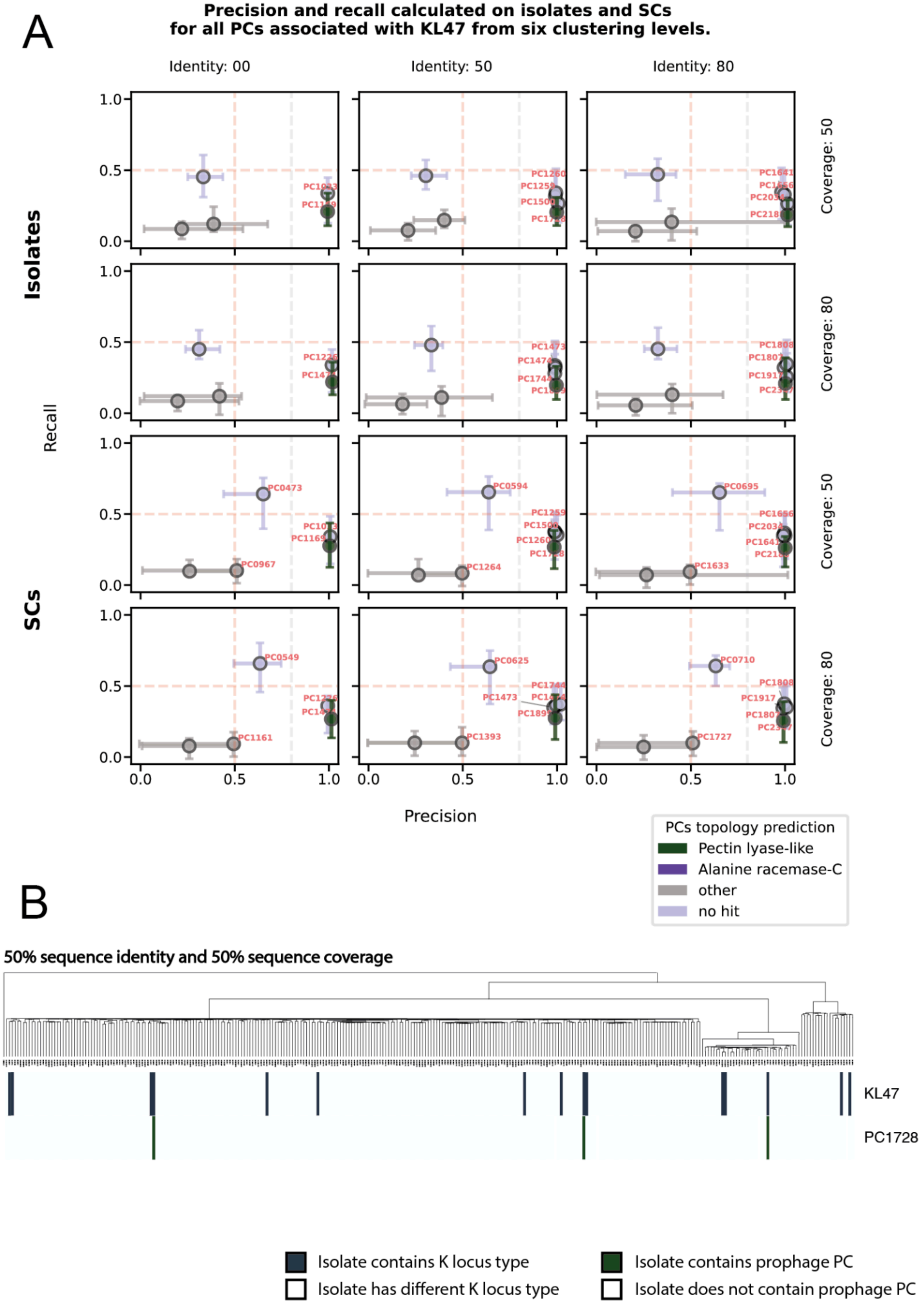
A
Precision and recall calculated on isolates and SCs
for all PCs associated with KL47 from six clustering levels.
Identity: 00
Identity: 50
Identity: 80
Coverage: 50
Coverage: 80
Isolates
SCs
Recall
Precision
0.0
0.5
1.0
PC0473
PC0967
PC0594
PC1264
PC0695
PC1633
PC0549
PC1161
PC0625
PC1473
PC1393
PC0710
PC1727
PCs topology prediction
Pectin lyase-like
Alanine racemase-C
other
no hit
B
50% sequence identity and 50% sequence coverage
KL47
PC1728
Isolate contains K locus type
Isolate has different K locus type
Isolate contains prophage PC
Isolate does not contain prophage PC

**Figure 12.** Identification of KL47-specific prophage depolymerases using GWAS. (A) The figure shows precision and recall with CI 95% for GWAS-linked PCs across six sequence-clustering thresholds. Colours distinguish predicted ECOD folds: Pectin lyase-like, Alanine racemase-C, other ECOD folds or no similarity to ECOD database detected. (B) Distribution of KL47 and associated PC with pectin lyase-like fold at the 50 % identity and 50 % bidirectional coverage level. The bacterial phylogenetic tree has been sub-sampled, but only isolates lacking both KL47 and the PCs with pectin lyase-like fold were removed.

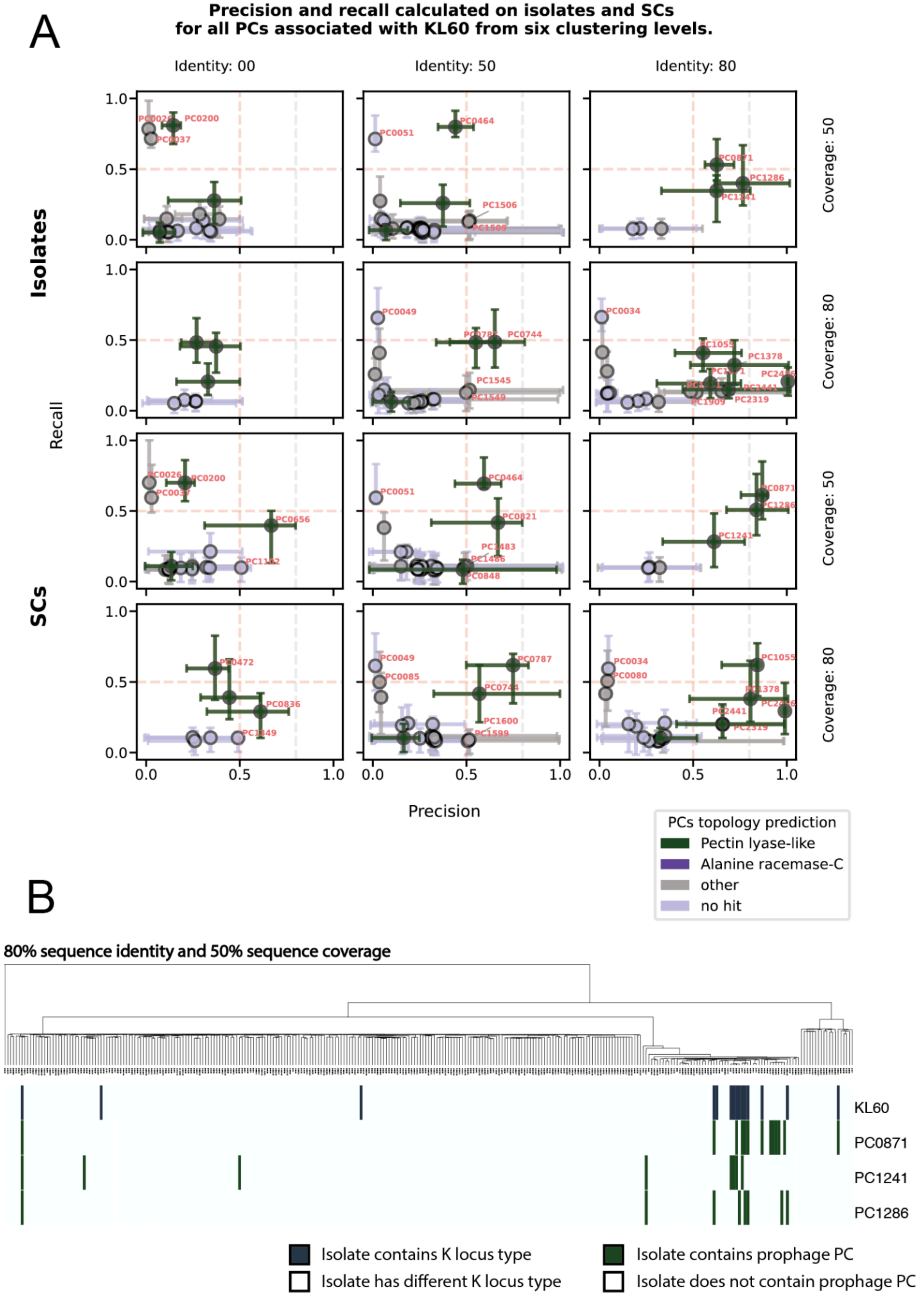
A
Precision and recall calculated on isolates and SCs
for all PCs associated with KL60 from six clustering levels.
Identity: 00
Identity: 50
Identity: 80
Coverage: 50
Coverage: 80
Coverage: 50
Coverage: 80
Isolates
SCs
Recall
Precision
0.0
0.5
1.0
PC0200
PC0037
PC0464
PC0051
PC1506
PC1509
PC0871
PC1286
PC1241
PC0049
PC0744
PC1545
PC1549
PC0034
PC1055
PC1378
PC1909
PC2319
PC0026
PC0656
PC0464
PC0051
PC0821
PC1486
PC0848
PC0472
PC0836
PC0049
PC0085
PC0787
PC1600
PC1599
PC0080
PC2441
PCs topology prediction
Pectin lyase-like
Alanine racemase-C
other
no hit
B
80% sequence identity and 50% sequence coverage
KL60
PC0871
PC1241
PC1286
Isolate contains K locus type
Isolate has different K locus type
Isolate contains prophage PC
Isolate does not contain prophage PC

**Figure 13.** Identification of KL60-specific prophage depolymerases using GWAS. (A) The figure shows precision and recall with CI 95% for GWAS-linked PCs across six sequence-clustering thresholds. Colours distinguish predicted ECOD folds: Pectin lyase-like, Alanine racemase-C, other ECOD folds or no similarity to ECOD database detected. (B) Distribution of KL60 and associated PC with pectin lyase-like fold at the 50 % identity and 50 % bidirectional coverage level. The bacterial phylogenetic tree has been sub-sampled, but only isolates lacking both KL60 and the PCs with pectin lyase-like fold were removed.

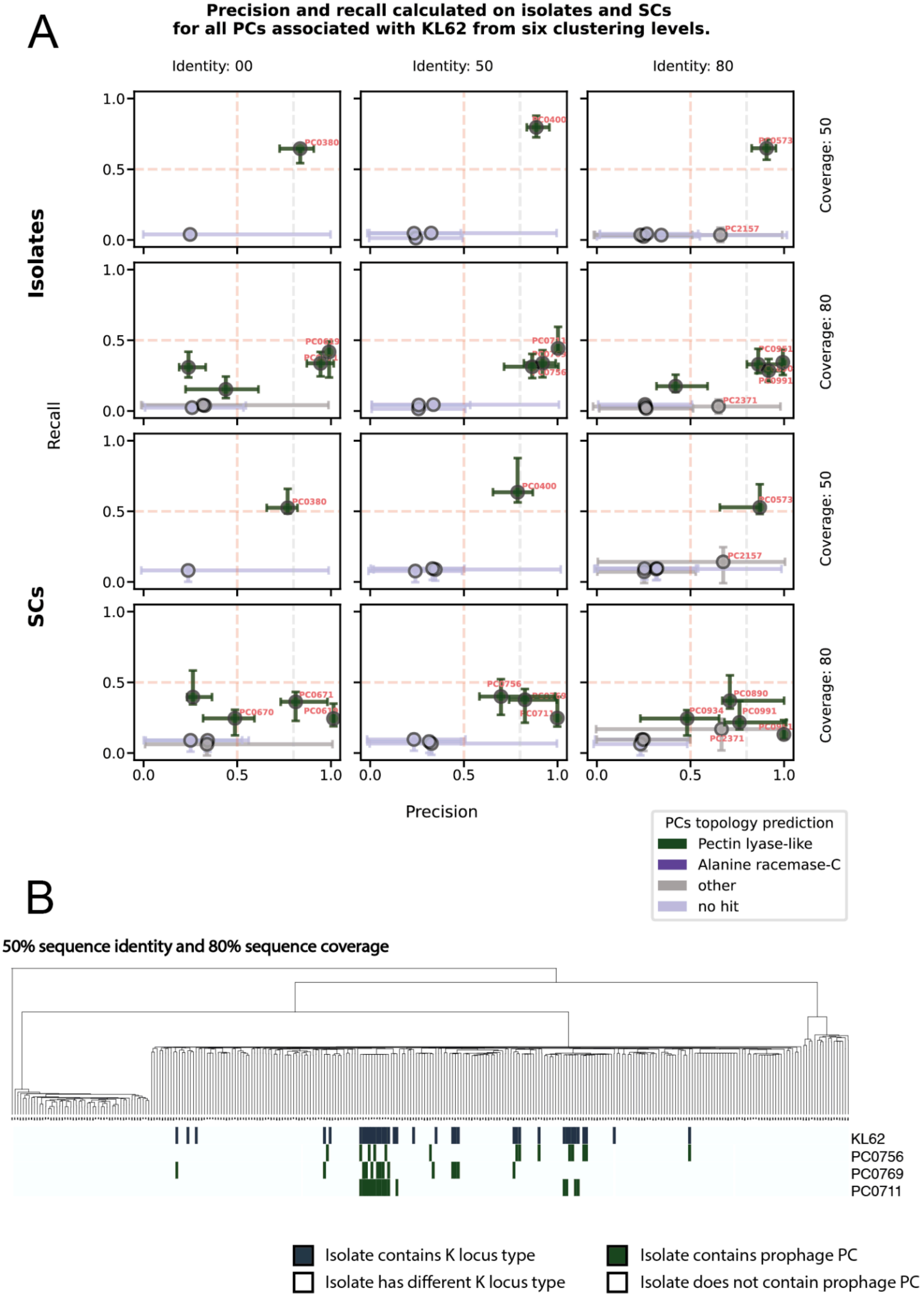
A
Precision and recall calculated on isolates and SCs
for all PCs associated with KL62 from six clustering levels.
Identity: 00
Identity: 50
Identity: 80
Coverage: 50
Coverage: 80
Coverage: 50
Coverage: 80
Isolates
SCs
Recall
Precision
PC0380
PC0400
PC0573
PC2157
PC0756
PC0991
PC2371
PC0671
PC0670
PC0711
PC0890
PC0934
PCs topology prediction
Pectin lyase-like
Alanine racemase-C
other
no hit
B
50% sequence identity and 80% sequence coverage
KL62
PC0756
PC0769
PC0711
Isolate contains K locus type
Isolate has different K locus type
Isolate contains prophage PC
Isolate does not contain prophage PC

**Figure 14.** Identification of KL62-specific prophage depolymerases using GWAS. (A) The figure shows precision and recall with CI 95% for GWAS-linked PCs across six sequence-clustering thresholds. Colours distinguish predicted ECOD folds: Pectin lyase-like, Alanine racemase-C, other ECOD folds or no similarity to ECOD database detected. (B) Distribution of KL62 and associated PC with pectin lyase-like fold at the 50 % identity and 50 % bidirectional coverage level. The bacterial phylogenetic tree has been sub-sampled, but only isolates lacking both KL62 and the PCs with pectin lyase-like fold were removed.

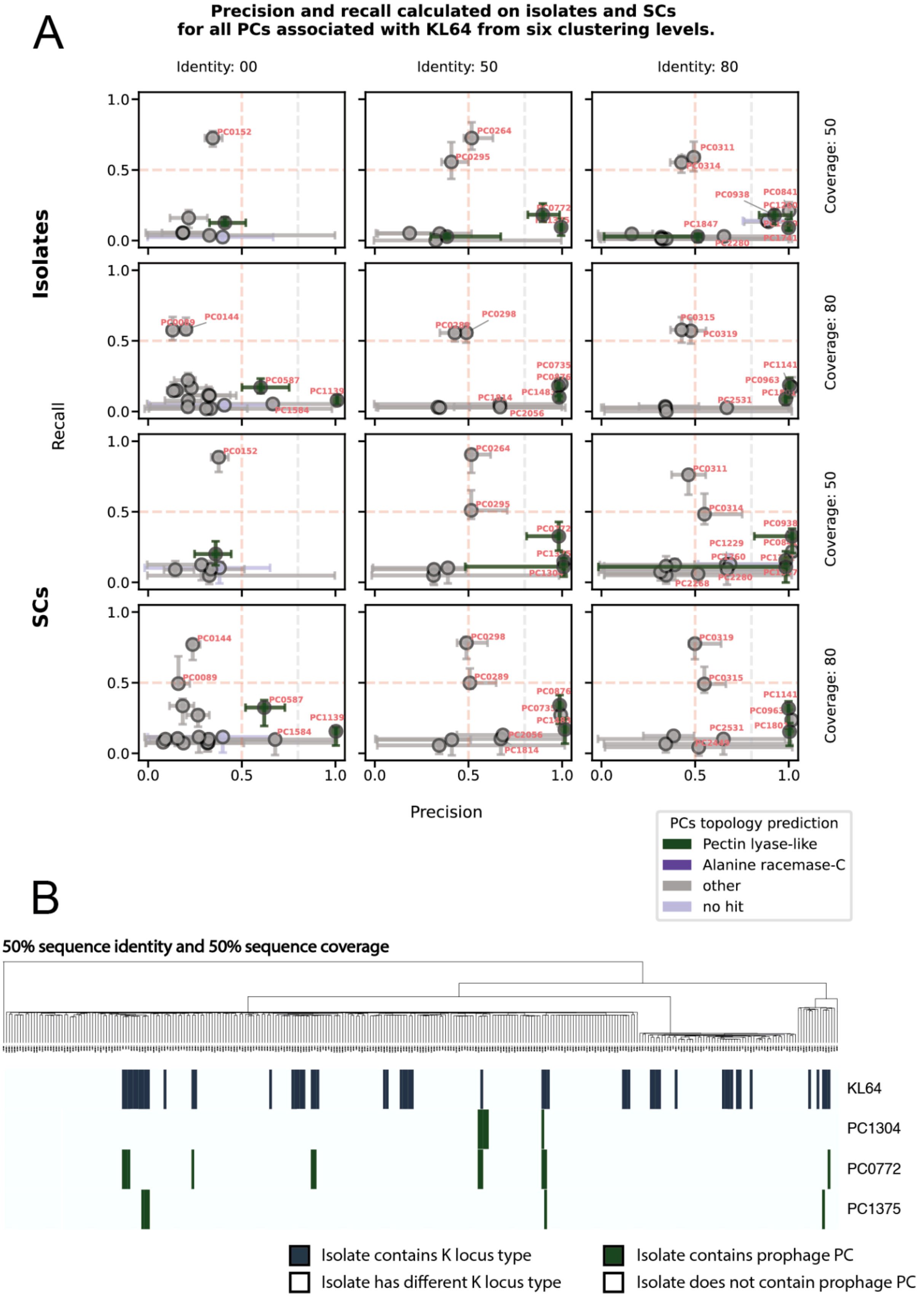
A
Precision and recall calculated on isolates and SCs
for all PCs associated with KL64 from six clustering levels.
Identity: 00
Identity: 50
Identity: 80
Coverage: 50
Coverage: 80
Coverage: 50
Coverage: 80
Isolates
SCs
Recall
Precision
PC0152
PC0264
PC0295
PC0311
PC0314
PC0144
PC0298
PC0315
PC0319
PC0587
PC1139
PC1584
PC0289
PC0772
PC1304
PC0876
PC0735
PC2056
PC1814
PC0938
PC0841
PC1847
PC2280
PC1141
PC0963
PC2531
PC1229
PC2268
PCs topology prediction
Pectin lyase-like
Alanine racemase-C
other
no hit
B
50% sequence identity and 50% sequence coverage
KL64
PC1304
PC0772
PC1375
Isolate contains K locus type
Isolate has different K locus type
Isolate contains prophage PC
Isolate does not contain prophage PC

**Figure 15.** Identification of KL64-specific prophage depolymerases using GWAS. (A) The figure shows precision and recall with CI 95% for GWAS-linked PCs across six sequence-clustering thresholds. Colours distinguish predicted ECOD folds: Pectin lyase-like, Alanine racemase-C, other ECOD folds or no similarity to ECOD database detected. (B) Distribution of KL64 and associated PC with pectin lyase-like fold at the 50 % identity and 50 % bidirectional coverage level. The bacterial phylogenetic tree has been sub-sampled, but only isolates lacking both KL64 and the PCs with pectin lyase-like fold were removed.

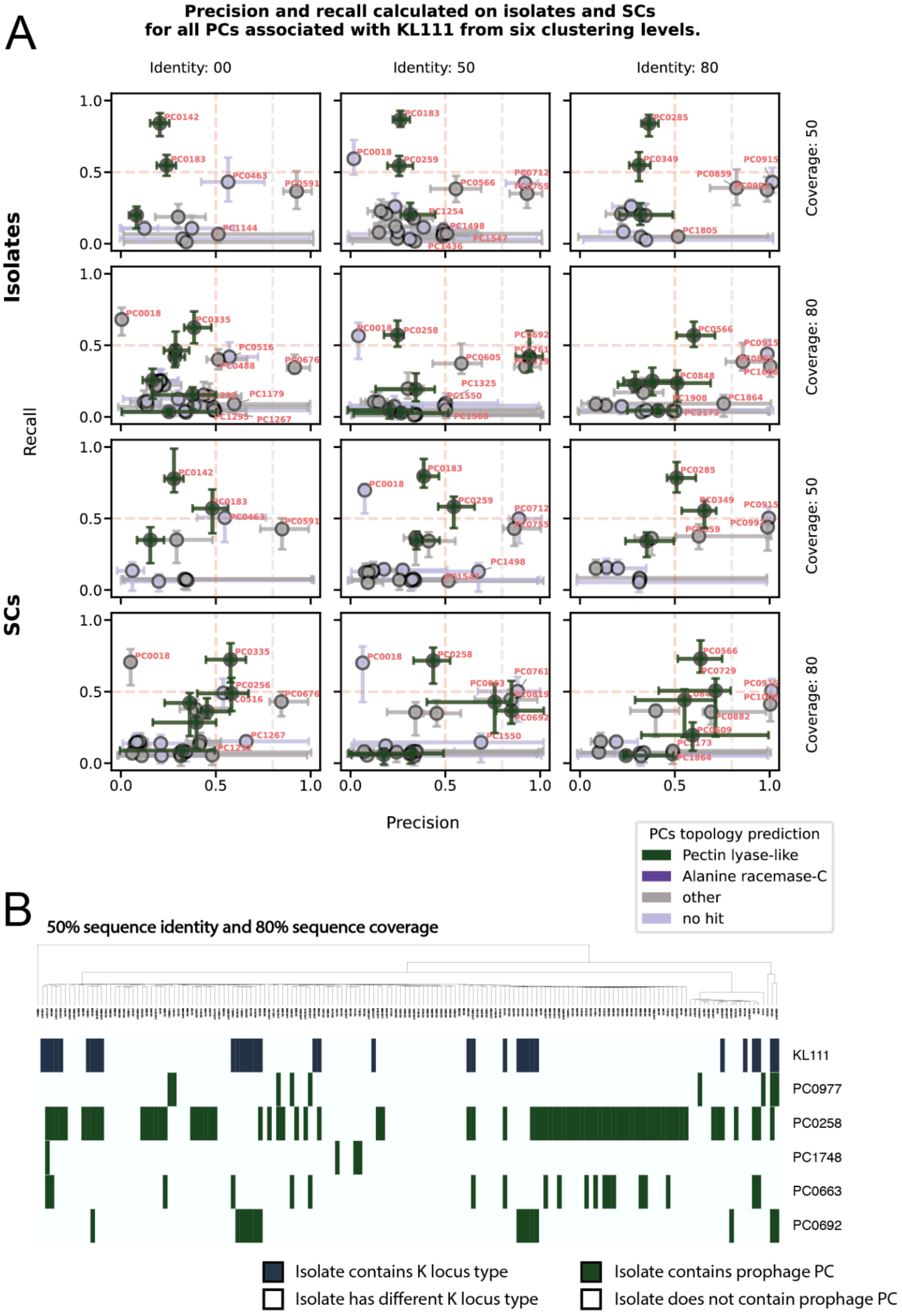

A
Precision and recall calculated on isolates and SCs
for all PCs associated with KL111 from six clustering levels.
Identity: 00
Identity: 50
Identity: 80
Coverage: 50
Coverage: 80
Coverage: 50
Coverage: 80
Isolates
SCs
Recall
Precision
PCs topology prediction
Pectin lyase-like
Alanine racemase-C
other
no hit
B
50% sequence identity and 80% sequence coverage
KL111
PC0977
PC0258
PC1748
PC0663
PC0692
Isolate contains K locus type
Isolate has different K locus type
Isolate contains prophage PC
Isolate does not contain prophage PC

**Figure 16.** Identification of KL111-specific prophage depolymerases using GWAS. (A) The figure shows precision and recall with CI 95% for GWAS-linked PCs across six sequence-clustering thresholds. Colours distinguish predicted ECOD folds: Pectin lyase-like, Alanine racemase-C, other ECOD folds or no similarity to ECOD database detected. (B) Distribution of KL111 and associated PC with pectin lyase-like fold at the 50 % identity and 80 % bidirectional coverage level. The bacterial phylogenetic tree has been sub-sampled, but only isolates lacking both KL111 and the PCs with pectin lyase-like fold were removed.

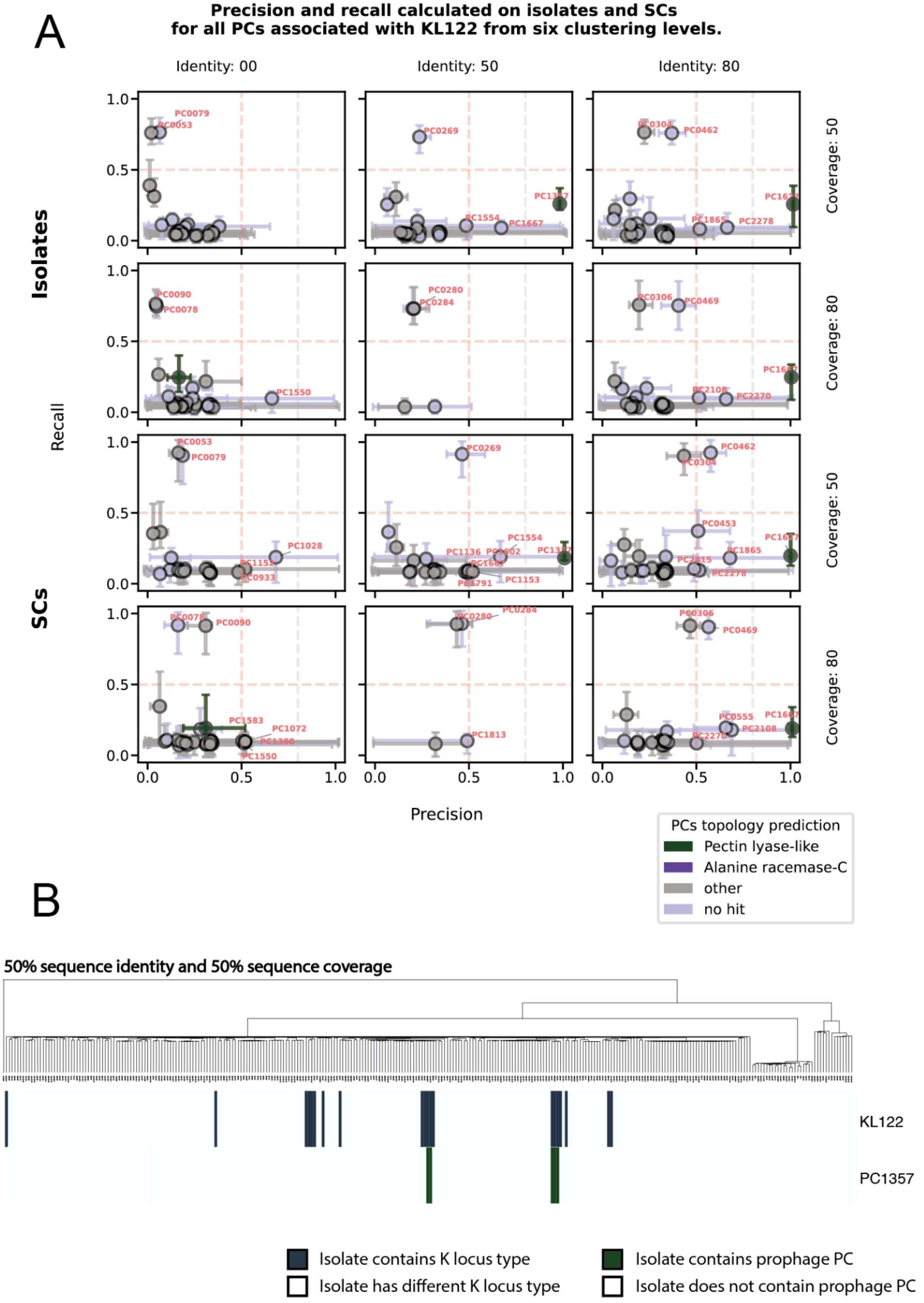
A
Precision and recall calculated on isolates and SCs
for all PCs associated with KL122 from six clustering levels.
Identity: 00
Identity: 50
Identity: 80
Coverage: 50
Coverage: 80
Coverage: 50
Coverage: 80
Isolates
SCs
Recall
Precision
PC0079
PC0053
PC0269
PC0462
PC1554
PC1667
PC1865
PC2278
PC0090
PC0078
PC0280
PC0284
PC0306
PC0469
PC1550
PC2108
PC2270
PC1028
PC0933
PC1136
PC1153
PC0453
PC1583
PC1072
PC1813
PC0555
PCs topology prediction
Pectin lyase-like
Alanine racemase-C
other
no hit
B
50% sequence identity and 50% sequence coverage
KL122
PC1357
Isolate contains K locus type
Isolate has different K locus type
Isolate contains prophage PC
Isolate does not contain prophage PC

**Figure 17.** Identification of KL122-specific prophage depolymerases using GWAS. (A) The figure shows precision and recall with CI 95% for GWAS-linked PCs across six sequence-clustering thresholds. Colours distinguish predicted ECOD folds: Pectin lyase-like, Alanine racemase-C, other ECOD folds or no similarity to ECOD database detected. (B) Distribution of KL122 and associated PC with pectin lyase-like fold at the 50 % identity and 50 % bidirectional coverage level. The bacterial phylogenetic tree has been sub-sampled, but only isolates lacking both KL122 and the PCs with pectin lyase-like fold were removed.

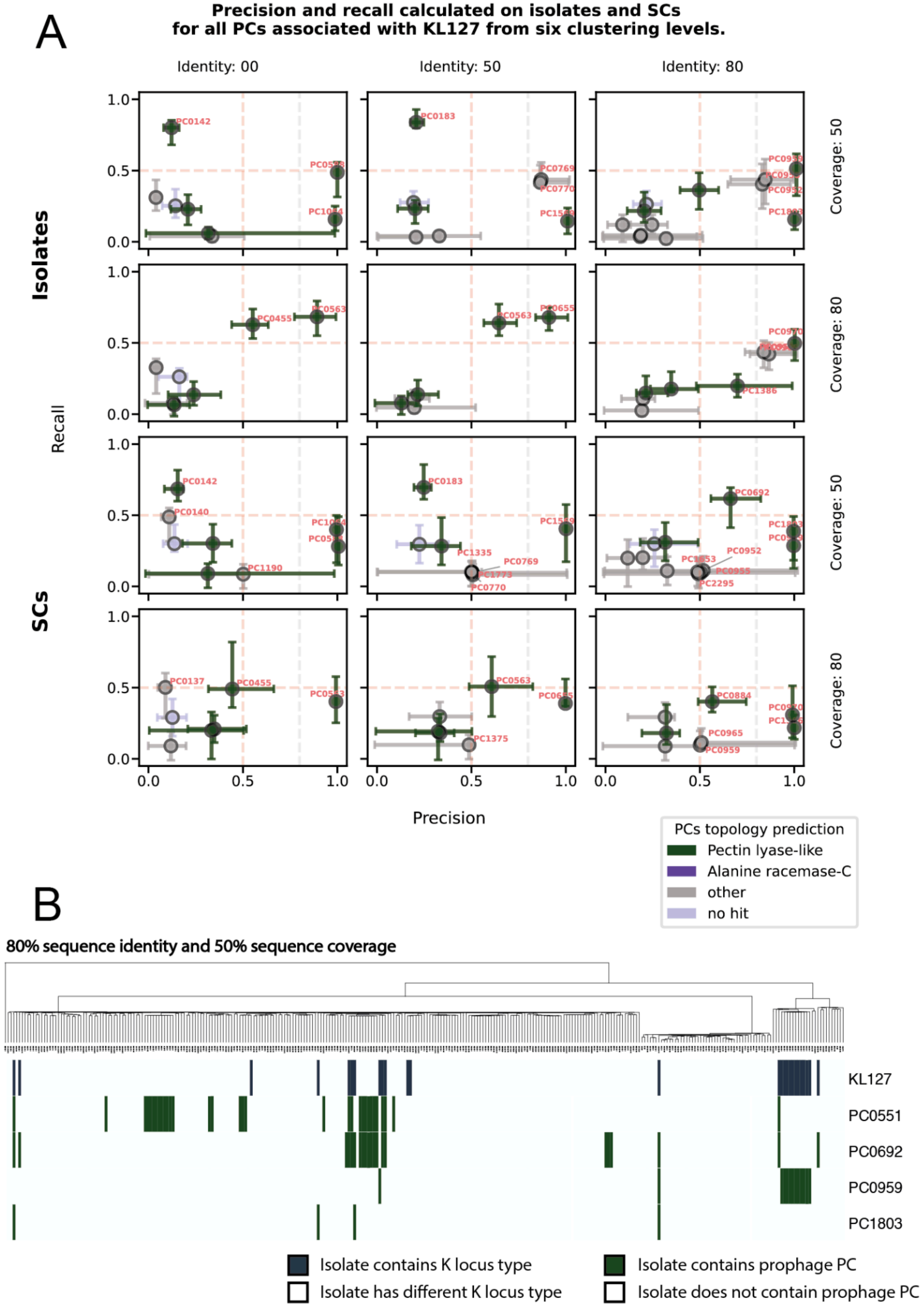
A
Precision and recall calculated on isolates and SCs
for all PCs associated with KL127 from six clustering levels.
Identity: 00
Identity: 50
Identity: 80
Coverage: 50
Coverage: 80
Isolates
SCs
Recall
Precision
PC0142
PC0183
PC0769
PC0770
PC0455
PC0563
PC0655
PC1386
PC0140
PC1190
PC1335
PC0692
PC0952
PC2295
PC0137
PC1375
PC0884
PC0965
PC0959
PCs topology prediction
Pectin lyase-like
Alanine racemase-C
other
no hit
B
80% sequence identity and 50% sequence coverage
KL127
PC0551
PC0692
PC0959
PC1803
Isolate contains K locus type
Isolate has different K locus type
Isolate contains prophage PC
Isolate does not contain prophage PC

**Figure 18.** Identification of KL127-specific prophage depolymerases using GWAS. (A) The figure shows precision and recall with CI 95% for GWAS-linked PCs across six sequence-clustering thresholds. Colours distinguish predicted ECOD folds: Pectin lyase-like, Alanine racemase-C, other ECOD folds or no similarity to ECOD database detected. (B) Distribution of KL127 and associated PC with pectin lyase-like fold at the 80 % identity and 50 % bidirectional coverage level. The bacterial phylogenetic tree has been sub-sampled, but only isolates lacking both KL127 and the PCs with pectin lyase-like fold were removed.
