## Supplementary material for "Capsular specificity in temperate phages of *Klebsiella pneumoniae* is driven by diverse receptor-binding enzymes": S2 Text

### Supplementary Text S2

#### Experimental validation of capsule depolymerase activity.

**Table 1 Experimental results for all active recombinant proteins identified through manual screening.** For each protein, the predicted host K-locus (based on prophage origin), experimentally confirmed K-locus specificity, expression level, and minimal halo-forming concentration (MHFC) are shown. Spot tests were performed using both purified proteins (tested at the initial concentration and in a 2-fold dilution series) and unpurified lysates as well as insoluble fractions, to comprehensively assess activity. The lowest concentration at which visible capsule degradation (halo formation) occurred was recorded as MHFC.

| **Protein name** | **319_37** | **738_68** | **1248_57** |
|---|---|---|---|
| **Host K-locus** | KL127 | KL143 | KL28 |
| **K-locus specificity** | KL127 | KL143 | KL23 |
| **Expression level** | Low | Medium | Medium |
| **Initial concentration** | 320 µg/ml | 795 µg/ml | 144 µg/ml |
| **MHFC** | 320 µg/ml | 0,087 µg/ml | 6,125 µg/ml |
| **Depolymerase activity (spot test)** | 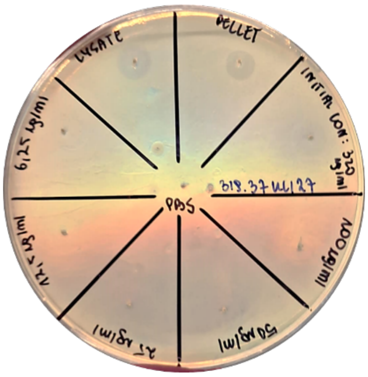  | 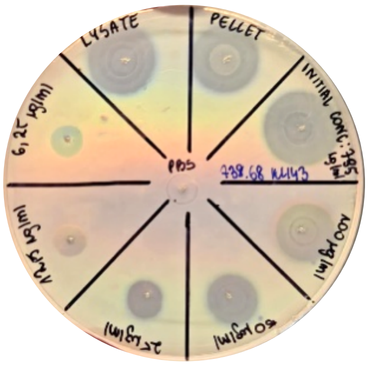  | 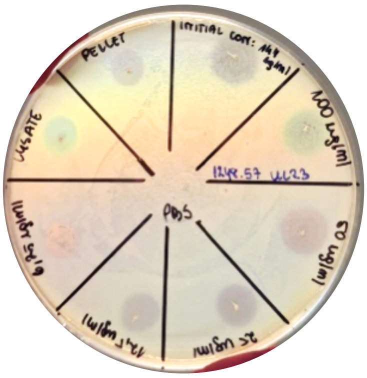  |

**Table 1. Experimental results for all active recombinant proteins identified through manual screening** (continued).

| Protein name | 1251_37 | 1409_59 | 1441_47 |
|---|---|---|---|
| Host K-locus | KL28 | KL28 | KL28 |
| K-locus specificity | KL28 | KL23 | KL23 |
| Expression level | Low | Low | Low |
| Initial concentration | 377 µg/ml | 130 µg/ml | 160 µg/ml |
| MHFC | 377 µg/ml | 25 µg/ml | 1,56 µg/ml |
| Depolymerase activity (spot test) | 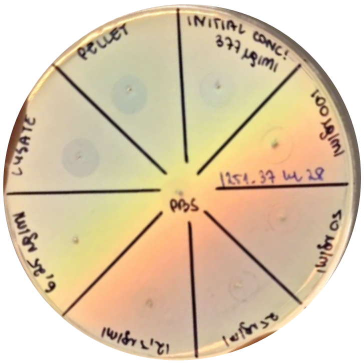 | 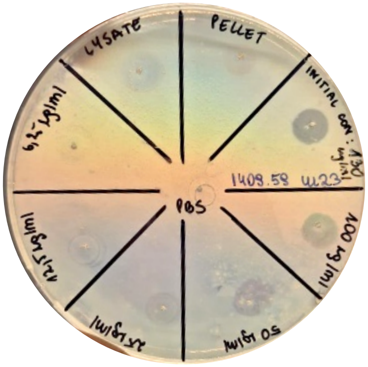 | 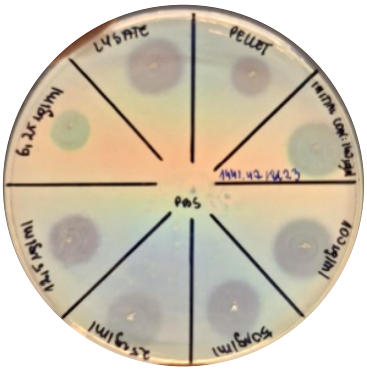 |

**Table 1. Experimental results for all active recombinant proteins identified through manual screening** (continued).

| Protein name | 617_77 | 434_33 | 248_38 |
|---|---|---|---|
| Host K-locus | KL38 | KL52 | KL55 |
| K-locus specificity | KL38 | KL52 | KL46 |
| Expression level | Medium | Medium | Low |
| Initial concentration | 434 µg/ml | 570 µg/ml | 260 µg/ml |
| MHFC | 12,5 µg/ml | 12,5 µg/ml | 6,25 µg/ml |
| Depolymerase activity (spot test) | 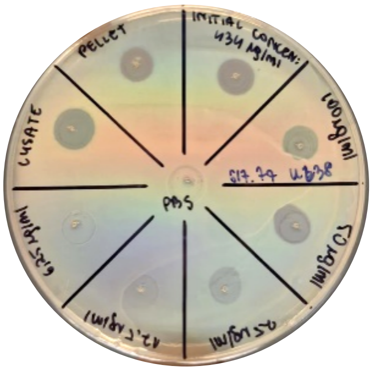 | 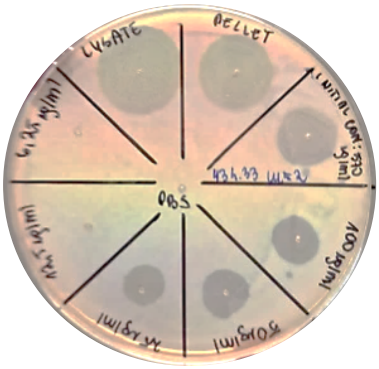 | 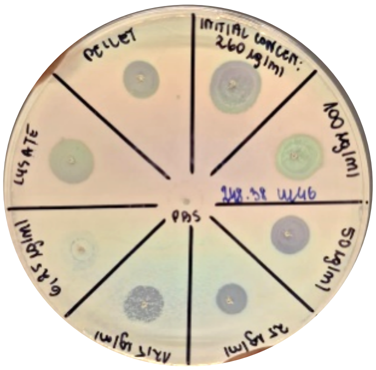 |

**Table 1. Experimental results for all active recombinant proteins identified through manual screening** (continued).

| **Protein name** | **1723_59** | **1724_71** | **914_74** |
|---|---|---|---|
| **Host K-locus** | KL60 | KL60 | KL62 |
| **K-locus specificity** | KL60 | KL60 | KL62 |
| **Expression level** | Low | Low | Low |
| **Initial concentration** | 261 µg/ml | 213 µg/ml | 120 µg/ml |
| **MHFC** | 261 µg/ml | 12,5 µg/ml | 3,125 µg/ml |
| **Depolymerase activity (spot test)** | 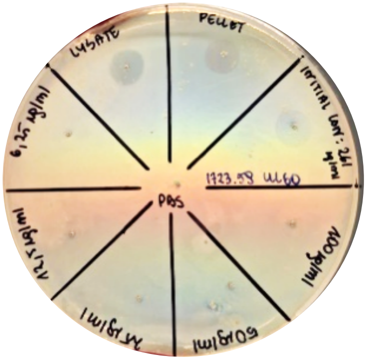 | 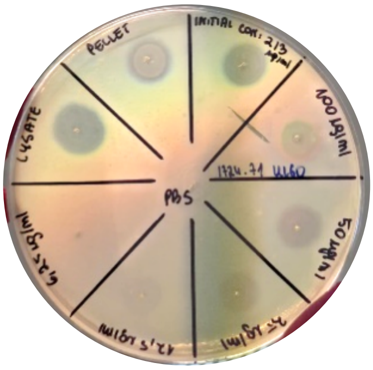 | 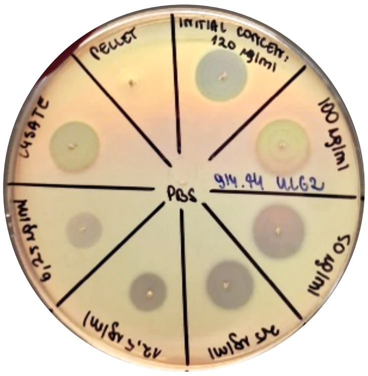 |

**Table 1. Experimental results for all active recombinant proteins identified through manual screening** (continued).

| **Protein name** | **914_77** | **1091_44** |
|---|---|---|
| **Host K-locus** | KL62 | KL64 |
| **K-locus specificity** | KL32 | KL64 |
| **Expression level** | Low | Low |
| **Initial concentration** | 143 µg/ml | 527 µg/ml |
| **MHFC** | --- | 25 µg/ml |
| **Depolymerase activity (spot test)** | 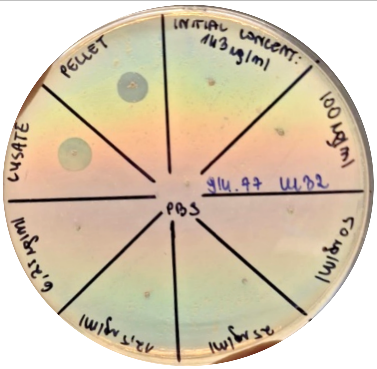 | 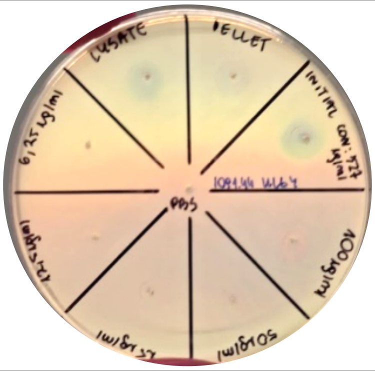 |

**Table 2. Experimental results for all active recombinant proteins identified through GWAS approach.** For each protein, the predicted host K-locus (based on prophage origin), experimentally confirmed K-locus specificity, expression level, and minimal halo-forming concentration (MHFC) are shown. Spot tests were performed using purified proteins (tested at the initial concentration and in a 2-fold dilution series). The lowest concentration at which visible capsule degradation (halo formation) occurred was recorded as MHFC.

| **Protein name** | **0367_12** | **0574_17** | **0391_11** |
|---|---|---|---|
| **Host K-locus** | KL62 | KL14 | KL62 |
| **K-locus specificity** | KL62 | KL14 | KL62 |
| **Expression level** | Medium | Medium | High |
| **Initial concentration** | 360 μg/ml | 4,5 mg/ml | 1,06 mg/ml |
| **MHFC** | 3,125 μg/ml | 6.25 μg/ml | 25 μg/ml |
| **Depolymerase activity (spot test)** | 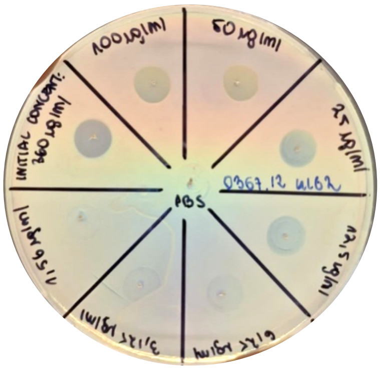  | 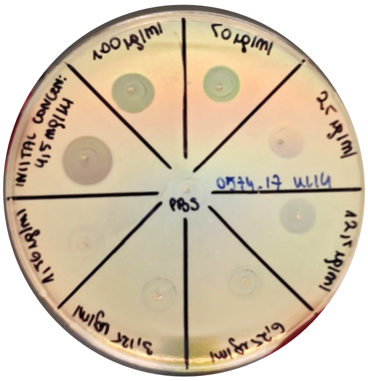  | 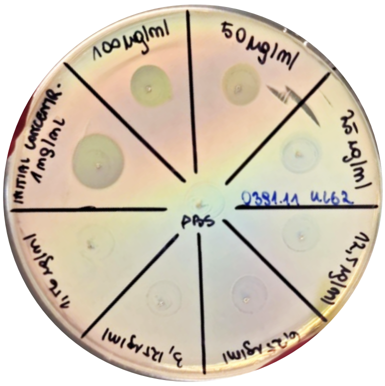 |

**Table 2. Experimental results for all active recombinant proteins identified through GWAS approach** (continued).

| Protein name | 0496_72 | 391_03 | |
|---|---|---|---|
| Host K-locus | KL122 | KL3 | |
| K-locus specificity | KL122 | KL3 | KL146 |
| Expression level | Medium | High | |
| Initial concentration | 710 µg/ml | 3,15 mg/ml | |
| MHFC | 1,56 µg/ml | 1,56 µg/ml | 3,125 µg/ml |
| Depolymerase activity (spot test) | 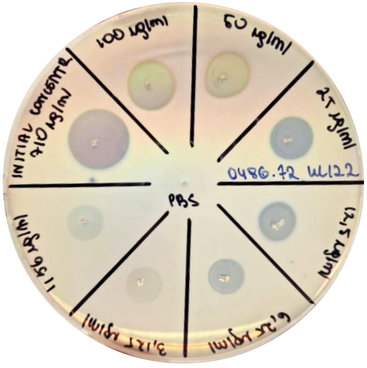 | 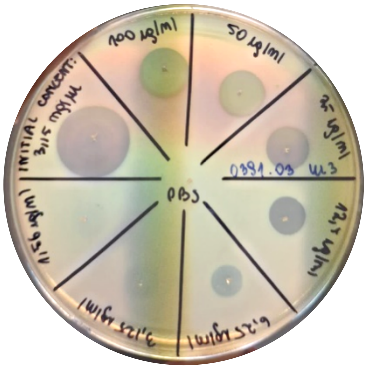 | 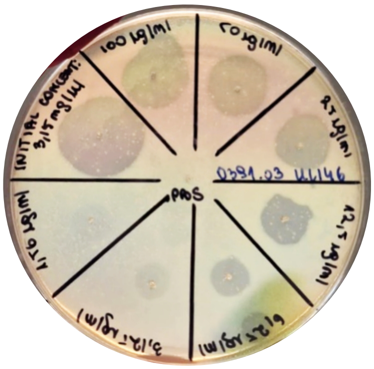 |

**Table 2. Experimental results for all active recombinant proteins identified through GWAS approach** (continued).

| Protein name | 184_43 |
|---|---|
| Host K-locus | KL111 |
| K-locus specificity | KL111 |
| Expression level | High |
| Initial concentration | 5 mg/ml |
| MHFC | 12,5 µg/ml |
| Depolymerase activity (spot test) | 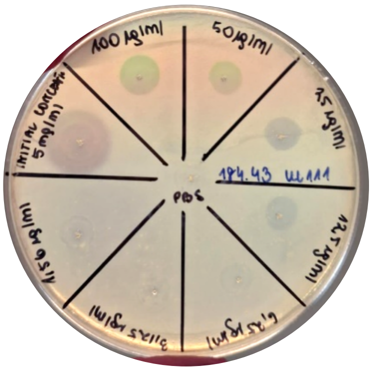 |

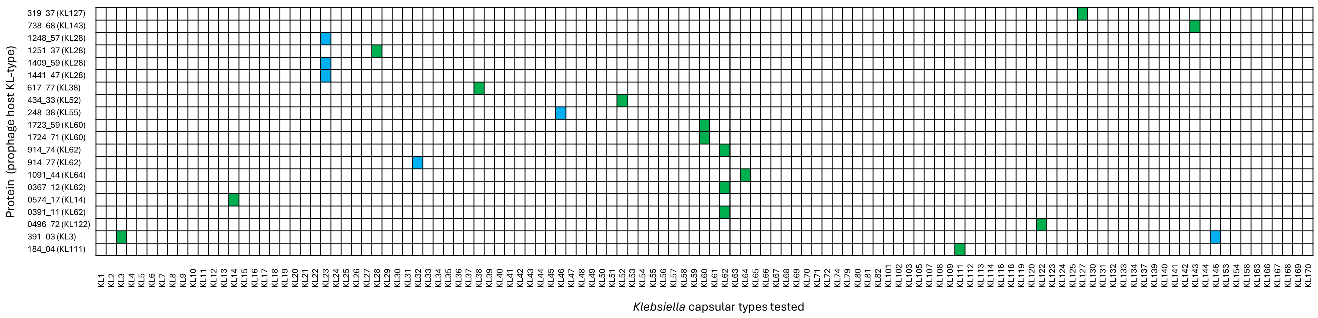


**Figure 1. Heatmap of depolymerase activity against Klebsiella pneumoniae capsular types.** Each recombinant depolymerase is shown along the X-axis, accompanied by the KL-type of its prophage host. The Y-axis lists all tested Klebsiella KL-types. Coloured squares represent positive activity signals, with green indicating activity against the prophage host KL-type and blue indicating activity against a distinct KL-type.
