## Supplementary material for "Capsular specificity in temperate phages of *Klebsiella pneumoniae* is driven by diverse receptor-binding enzymes": S3 Text

### Supplementary Text S3

#### Sequence and structure comparison of predicted and verified depolymerases

Here we provide a detailed comparison of protein sequences from the sequence similarity network shown in Figure 4B. Each figure in this supplementary text corresponds to one connected component of the network (panel A). A representative protein is then selected as a query, and its BLASTP alignments to other proteins in the component are visualised (panel B), excluding those which could not be successfully produced (grey nodes in the network). Alignment figures indicate the percentage identity for the aligned regions. Panel C shows AlphaFold3-predicted structural homotrimeric models for each of the aligned proteins.

To compare the structures of receptor-binding domains (RBDs) of the modelled proteins, we looked for the separating α-helix at the junction (as shown in Figure 7b of Ouyeng R. et al. Nat Comm 2022, PMID 36433970) to trim off the N-terminal from the RBD. Pairwise TM scores for the RBD were then computed using US-align (Zhang et al. 36038728 Nat Methods 2022, PMID 36038728) in oligomer alignment mode and additionally verified visually by aligning the two RBD structures with pymol.

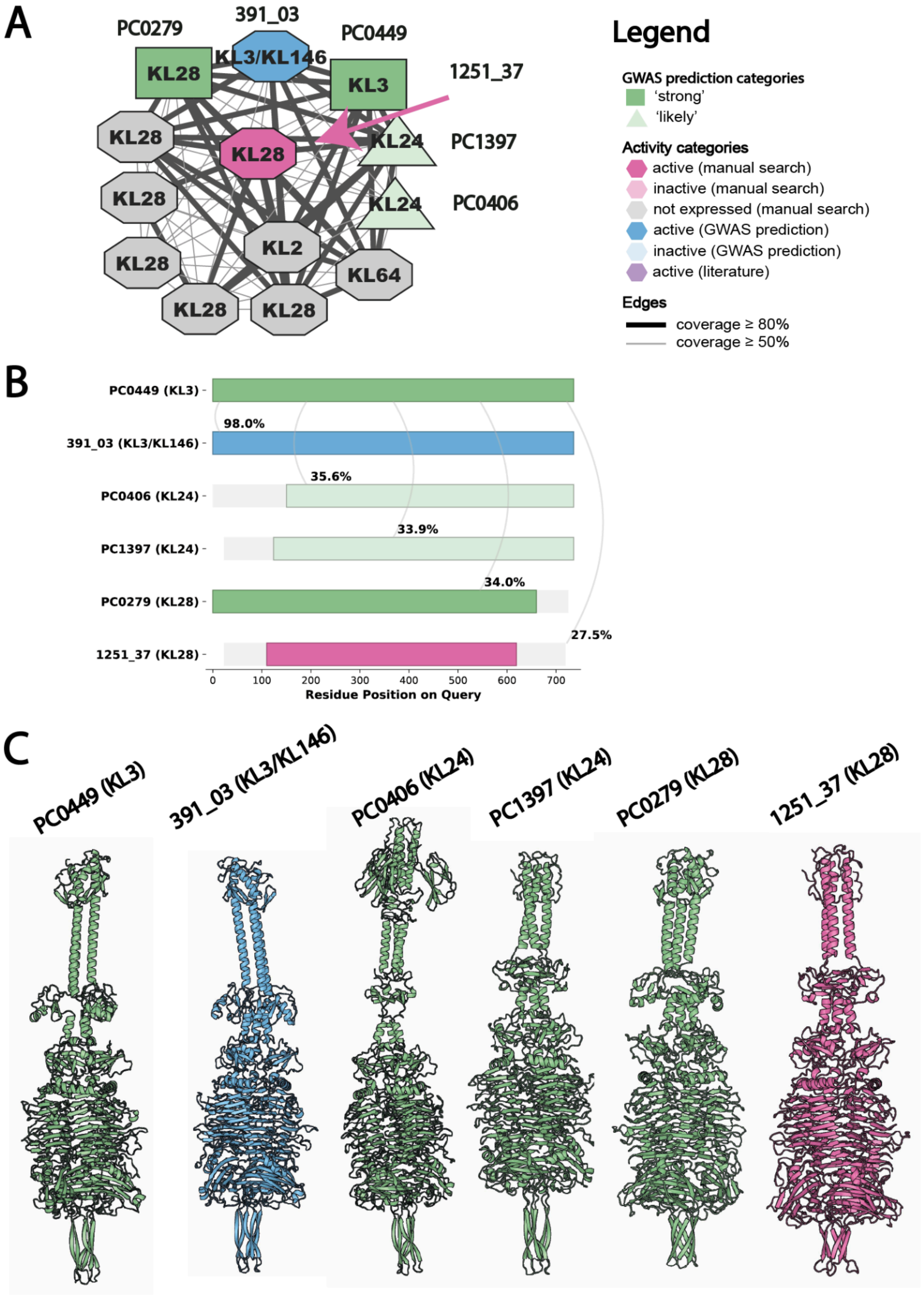
A
PC0279
391_03
PC0449
1251_37
KL28
KL3/KL146
KL3
KL28
KL28
KL24
PC1397
KL28
KL24
PC0406
KL2
KL28
KL64
KL28
KL28
Legend
GWAS prediction categories
'strong'
'likely'
Activity categories
active (manual search)
inactive (manual search)
not expressed (manual search)
active (GWAS prediction)
inactive (GWAS prediction)
active (literature)
Edges
coverage ≥ 80%
coverage ≥ 50%
B
PC0449 (KL3)
98.0%
391_03 (KL3/KL146)
35.6%
PC0406 (KL24)
33.9%
PC1397 (KL24)
34.0%
PC0279 (KL28)
27.5%
1251_37 (KL28)
0
100
200
300
400
500
600
700
Residue Position on Query
C
PC0449 (KL3)
391_03 (KL3/KL146)
PC0406 (KL24)
PC1397 (KL24)
PC0279 (KL28)
1251_37 (KL28)

**Figure 1. Analysis of the connected component with the KL3/KL28 predictions.** (A) Connected component of the sequence similarity containing 4 GWAS predictions: KL3 (PC0449; strong, dark green), KL28 (PC0279; strong, dark green), KL24 (PC0406; likely, light green), and KL24 (PC1397; likely, light green). A comparison of the representative sequences from these clusters revealed sequence similarity against 8 recombinant proteins, of which one was active against KL28 (1251_37; magenta) and the remaining 7 were unsuccessfully produced. We also commercially prepared a representative of the KL3 prediction (391_03; blue), which was active and is shown in the network. (B) Pairwise BLASTP alignments visualization of the query sequence (PC0449; KL3) against other GWAS predictions and the two active proteins. (C) Structural homotrimeric models generated by AlphaFold3 of the six protein sequences from panel B. The TM-score of alignment between the RBDs of PC0279 and 1251_37 is 0.99.

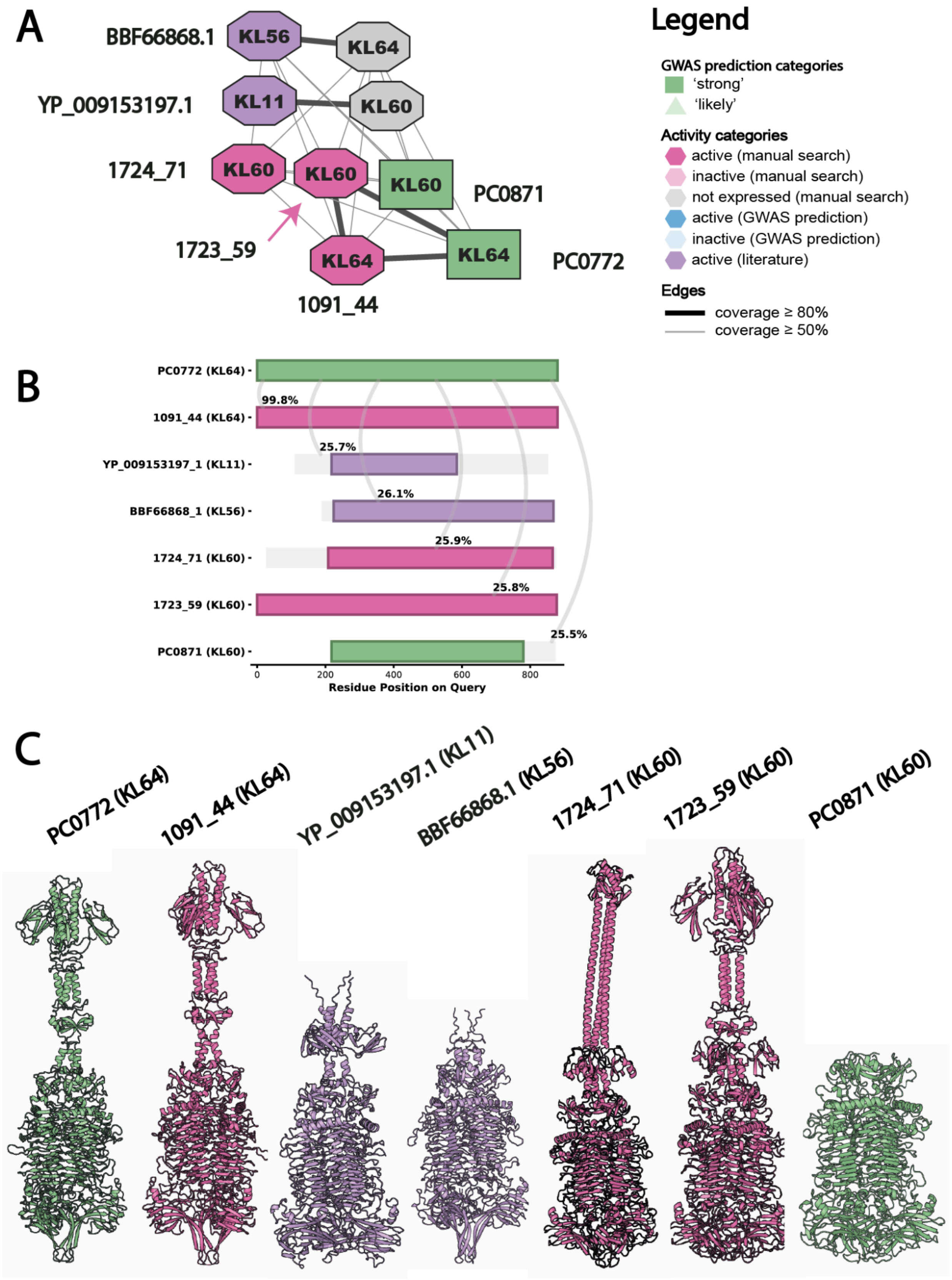
A
BBF66868.1
KL56
KL64
YP_009153197.1
KL11
KL60
1724_71
KL60
KL60
KL60
PC0871
1723_59
KL64
KL64
PC0772
1091_44
Legend
GWAS prediction categories
'strong'
'likely'
Activity categories
active (manual search)
inactive (manual search)
not expressed (manual search)
active (GWAS prediction)
inactive (GWAS prediction)
active (literature)
Edges
coverage ≥ 80%
coverage ≥ 50%
B
PC0772 (KL64)
1091_44 (KL64)
YP_009153197_1 (KL11)
BBF66868_1 (KL56)
1724_71 (KL60)
1723_59 (KL60)
PC0871 (KL60)
99.8%
25.7%
26.1%
25.9%
25.8%
25.5%
0
200
400
600
800
Residue Position on Query
C
PC0772 (KL64)
1091_44 (KL64)
YP_009153197.1 (KL11)
BBF66868.1 (KL56)
1724_71 (KL60)
1723_59 (KL60)
PC0871 (KL60)

**Figure 2. Analysis of the connected component with the KL60/KL64 predictions.** (A) Connected component of the sequence similarity containing 2 GWAS predictions: KL64 (PC0772; strong, dark green) and KL60 (PC0871; strong, dark green). A comparison of the representative sequences from these clusters revealed similarity substantial against 6 recombinant proteins, of which one was active against KL64 (1091_44; magenta), one was active against KL56 (BBF66868.1; purple), one was active against KL11 (YP_009153197.1; purple), two were active against KL60 (1724_71; magenta) (1723_59; magenta), and 2 were unsuccessfully produced. (B) Pairwise BLASTP alignments visualization of the query sequence (PC0772; KL64) against other GWAS predictions and the five active proteins. (C) Structural homotrimeric models generated by AlphaFold3 of the seven protein sequences from panel B. The TM-score of alignment between PC0871 and the RBDs of 1724_71 is 0.98 and between PC0871 and the RBDs of 1723_59 is 0.97.

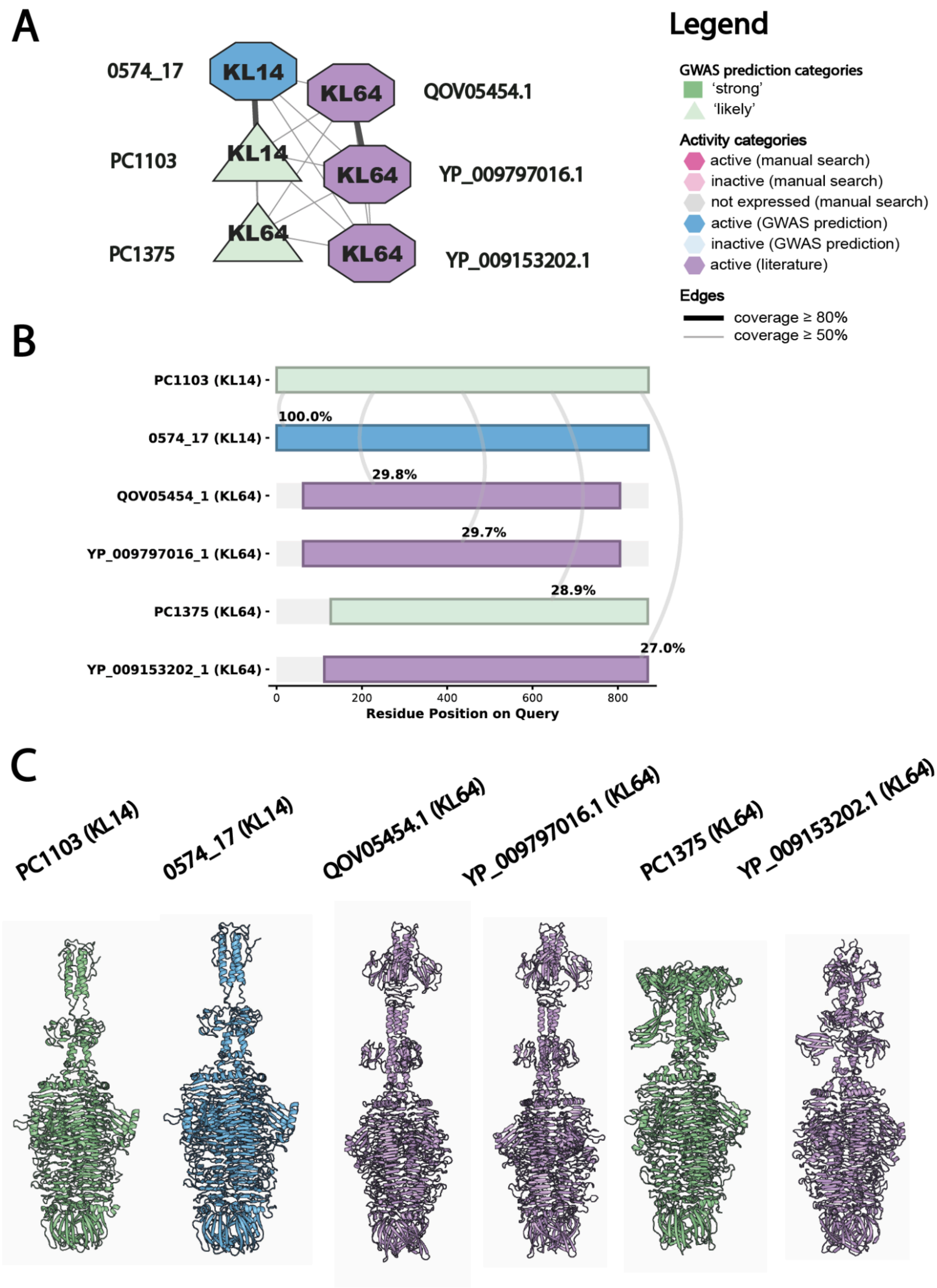
A
0574_17
KL14
KL64
QOV05454.1
PC1103
KL14
KL64
YP_009797016.1
PC1375
KL64
KL64
YP_009153202.1
Legend
GWAS prediction categories
'strong'
'likely'
Activity categories
active (manual search)
inactive (manual search)
not expressed (manual search)
active (GWAS prediction)
inactive (GWAS prediction)
active (literature)
Edges
coverage ≥ 80%
coverage ≥ 50%
B
PC1103 (KL14)
100.0%
0574_17 (KL14)
29.8%
QOV05454_1 (KL64)
29.7%
YP_009797016_1 (KL64)
28.9%
PC1375 (KL64)
27.0%
YP_009153202_1 (KL64)
0
200
400
600
800
Residue Position on Query
C
PC1103 (KL14)
0574_17 (KL14)
QOV05454.1 (KL64)
YP_009797016.1 (KL64)
PC1375 (KL64)
YP_009153202.1 (KL64)

**Figure 3. Analysis of the connected component with the KL14/KL64 predictions.** (A) Connected component of the sequence similarity containing 2 GWAS predictions: KL14 (PC1103; likely, light green), KL64 (PC1375; likely, light green). A comparison of the representative sequences from these clusters revealed similarity substantial against 3 recombinant proteins, of which all three were active against KL64 (QOV05454.1; purple) (YP_009797016.1; purple) (YP_009153202.1; purple). We also commercially produced a representative of the KL14 prediction (0574_17; blue), which was active and is shown in the network. (B) Pairwise BLASTP alignments visualization of the query sequence (PC1103; KL14) against other GWAS predictions and the four active proteins. (C) Structural homotrimeric models generated by AlphaFold3 of the six protein sequences from panel B. TM-score of alignment between the RBD of PC1375 and the RBDs of: (i) QOV05454_1 is 0.98, (ii) YP_009797016_1 is 0.98, (iii) YP_009153202_1 is 0.99.

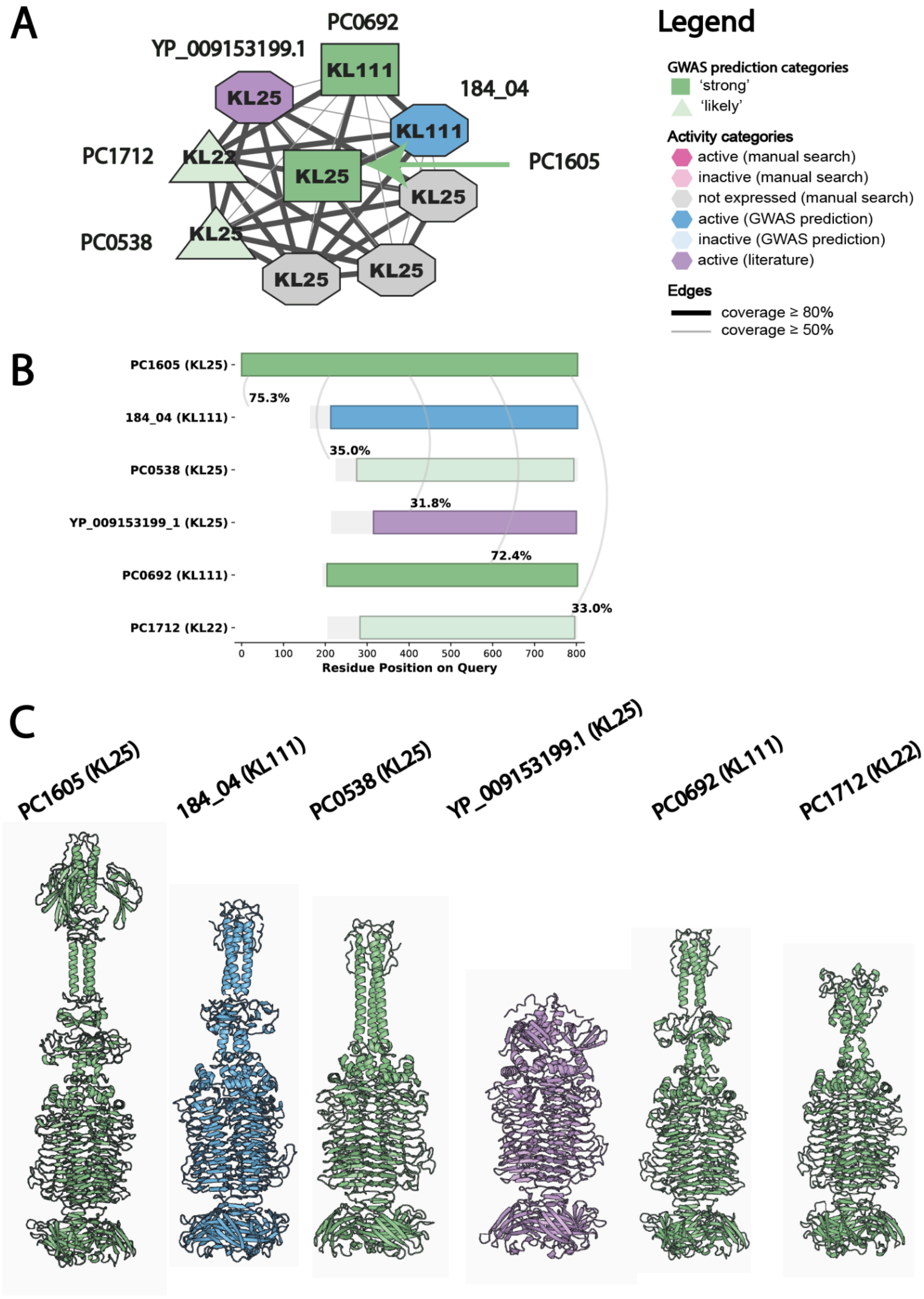
A
PC0692
YP_009153199.1
KL111
KL25
184_04
KL111
PC1712
KL22
KL25
PC1605
KL25
PC0538
KL25
KL25
KL25
Legend
GWAS prediction categories
'strong'
'likely'
Activity categories
active (manual search)
inactive (manual search)
not expressed (manual search)
active (GWAS prediction)
inactive (GWAS prediction)
active (literature)
Edges
coverage ≥ 80%
coverage ≥ 50%
B
PC1605 (KL25)
75.3%
184_04 (KL111)
35.0%
PC0538 (KL25)
31.8%
YP_009153199_1 (KL25)
72.4%
PC0692 (KL111)
33.0%
PC1712 (KL22)
0
100
200
300
400
500
600
700
800
Residue Position on Query
C
PC1605 (KL25)
184_04 (KL111)
PC0538 (KL25)
YP_009153199.1 (KL25)
PC0692 (KL111)
PC1712 (KL22)

**Figure 4. Analysis of the connected component with the KL22/KL25/KL111 predictions.** (A) Connected component of the sequence similarity containing 4 GWAS predictions: KL22 (PC1712; likely, light green), KL25 (PC0538; likely, light green), KL25 (PC1605; strong, dark green) and KL111 (PC0692, strong, dark green). A comparison of the representative sequences from these clusters revealed similarity substantial against 4 recombinant proteins, of which one was active against KL25 (YP_009153199.1; purple) and the remaining 3 were unsuccessfully produced. We also commercially produced a representative of the KL111 prediction (184_04; blue), which was active and is shown in the network. (B) Pairwise BLASTP alignments visualization of the query sequence (PC1605; KL25) against other GWAS predictions and the two active proteins. (C) Structural homotrimeric models generated by AlphaFold3 of the six protein sequences from panel B. The TM-score of alignment between the RBD of PC1605 and the RBD of YP_009153199_1 is 0.94. The TM-score of alignment between the RBD of PC0538 and the RBD of YP_009153199_1 is 0.97. The TM-score of alignment between the RBD of PC0692 and the RBD of YP_009153199_1 is 0.61. The TM-score of alignment between the RBD of PC1712 and the RBD of YP_009153199_1 is 0.96.

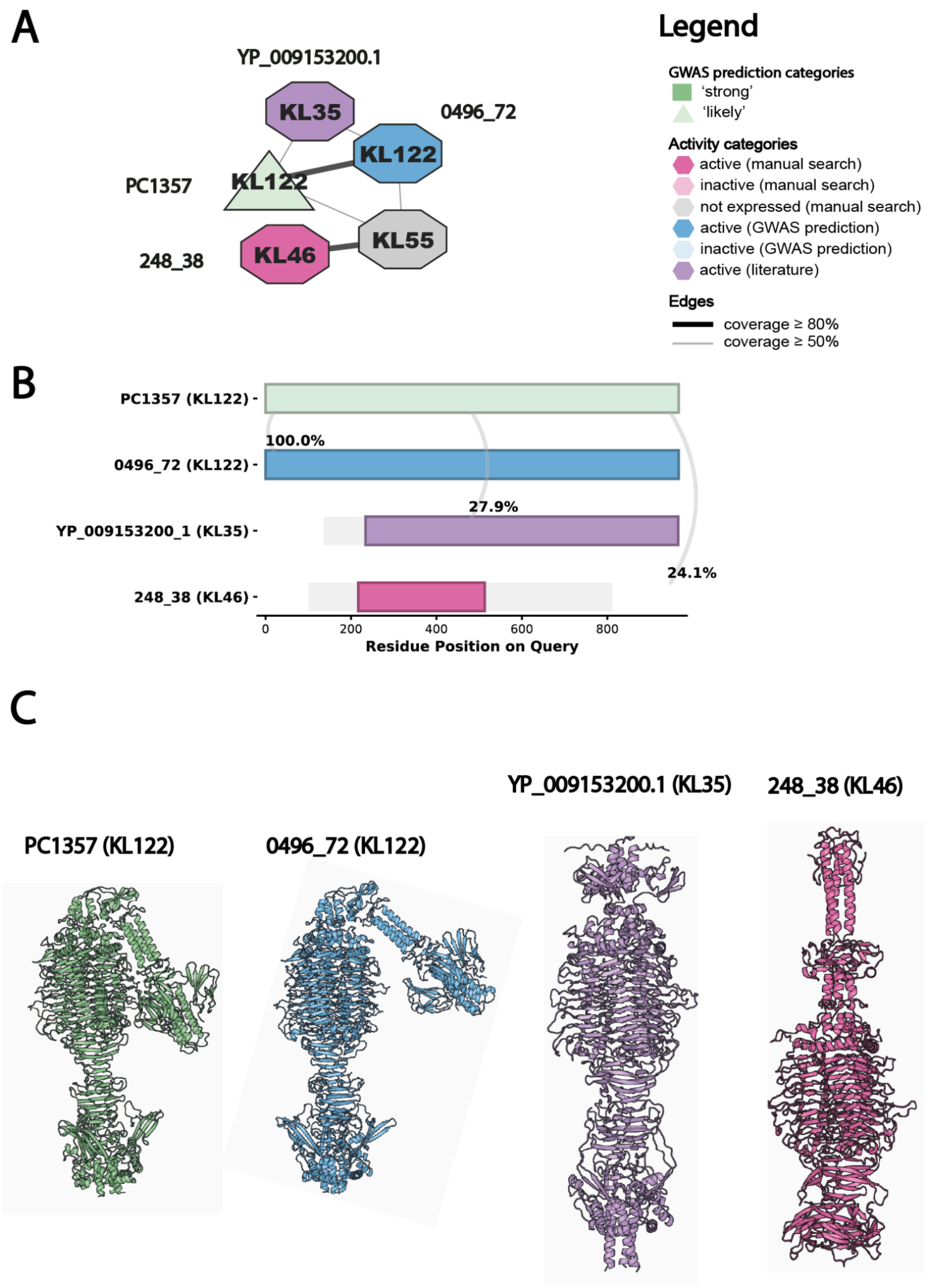
A
Legend
YP_009153200.1
KL35
0496_72
KL122
PC1357
KL122
KL55
248_38
KL46
GWAS prediction categories
'strong'
'likely'
Activity categories
active (manual search)
inactive (manual search)
not expressed (manual search)
active (GWAS prediction)
inactive (GWAS prediction)
active (literature)
Edges
coverage ≥ 80%
coverage ≥ 50%
B
PC1357 (KL122)
100.0%
0496_72 (KL122)
27.9%
YP_009153200_1 (KL35)
24.1%
248_38 (KL46)
0
200
400
600
800
Residue Position on Query
C
YP_009153200.1 (KL35)
248_38 (KL46)
PC1357 (KL122)
0496_72 (KL122)

**Figure 5. Analysis of the connected component with the KL122 prediction.** (A) Connected component of the sequence similarity containing 1 GWAS prediction: KL122 (PC1357; likely, light green). A comparison of the representative sequence from this cluster revealed similarity substantial against 3 recombinant proteins of which one was active against KL35 (YP_009153200.1; purple), one was active against KL46 (248_38; magenta) and one was unsuccessfully produced. We also commercially produced a representative of the KL122 prediction (0496_72; blue), which was active and is shown in the network. (B) Pairwise BLASTP alignments visualization of the query sequence (PC1357; KL122) against the three active proteins. (C) Structural homotrimeric models generated by AlphaFold3 of the three protein sequences from panel B. The TM-score of alignment between the RBD of PC1357 and the RBD of (i) YP_009153200.1 is 0.68 and (ii) 248_38 is 0.75.

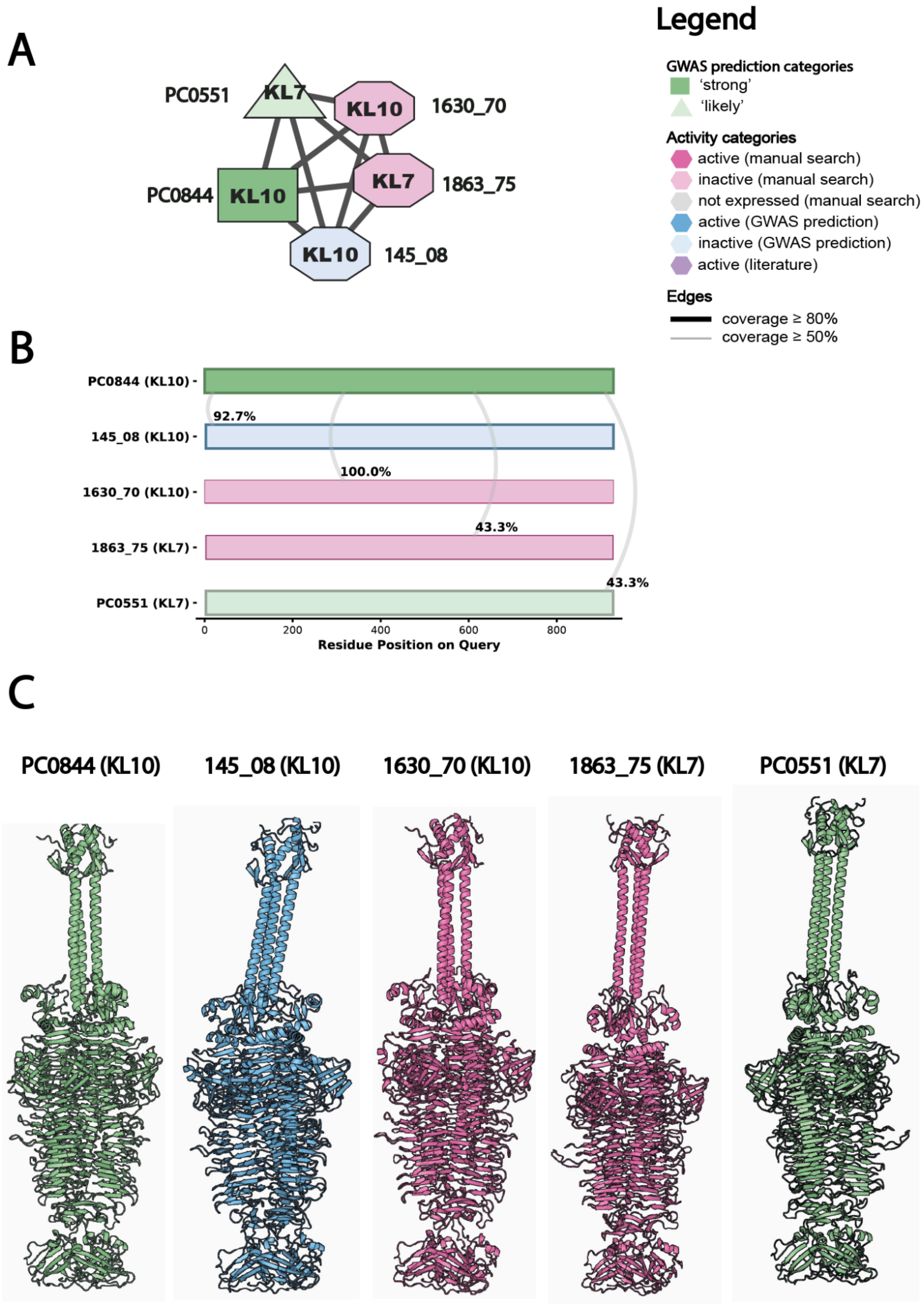
A
PC0551
KL7
KL10
1630_70
PC0844
KL10
KL7
1863_75
KL10
145_08
Legend
GWAS prediction categories
'strong'
'likely'
Activity categories
active (manual search)
inactive (manual search)
not expressed (manual search)
active (GWAS prediction)
inactive (GWAS prediction)
active (literature)
Edges
coverage ≥ 80%
coverage ≥ 50%
B
PC0844 (KL10)
145_08 (KL10)
1630_70 (KL10)
1863_75 (KL7)
PC0551 (KL7)
92.7%
100.0%
43.3%
43.3%
0
200
400
600
800
Residue Position on Query
C
PC0844 (KL10)
145_08 (KL10)
1630_70 (KL10)
1863_75 (KL7)
PC0551 (KL7)

**Figure 6. Analysis of the connected component with the KL10/KL7 predictions.** (A) Connected component of the sequence similarity containing 2 GWAS predictions: KL10 (PC0844; strong, dark green), KL7 (PC0551; likely, light green). A comparison of the representative sequences from these clusters revealed similarity substantial against 2 recombinant proteins, of which both KL10 (1630_70; light magenta) and KL7 (1863_75; light magenta) were produced, but inactive. We also commercially produced a representative of the KL10 prediction (145_08; blue), which was inactive and is shown in the network. (B) Pairwise BLASTP alignments visualization of the query sequence (PC0844; KL10) against other GWAS predictions and the three inactive proteins. (C) Structural homotrimeric models generated by AlphaFold3 of the five protein sequences from panel B.

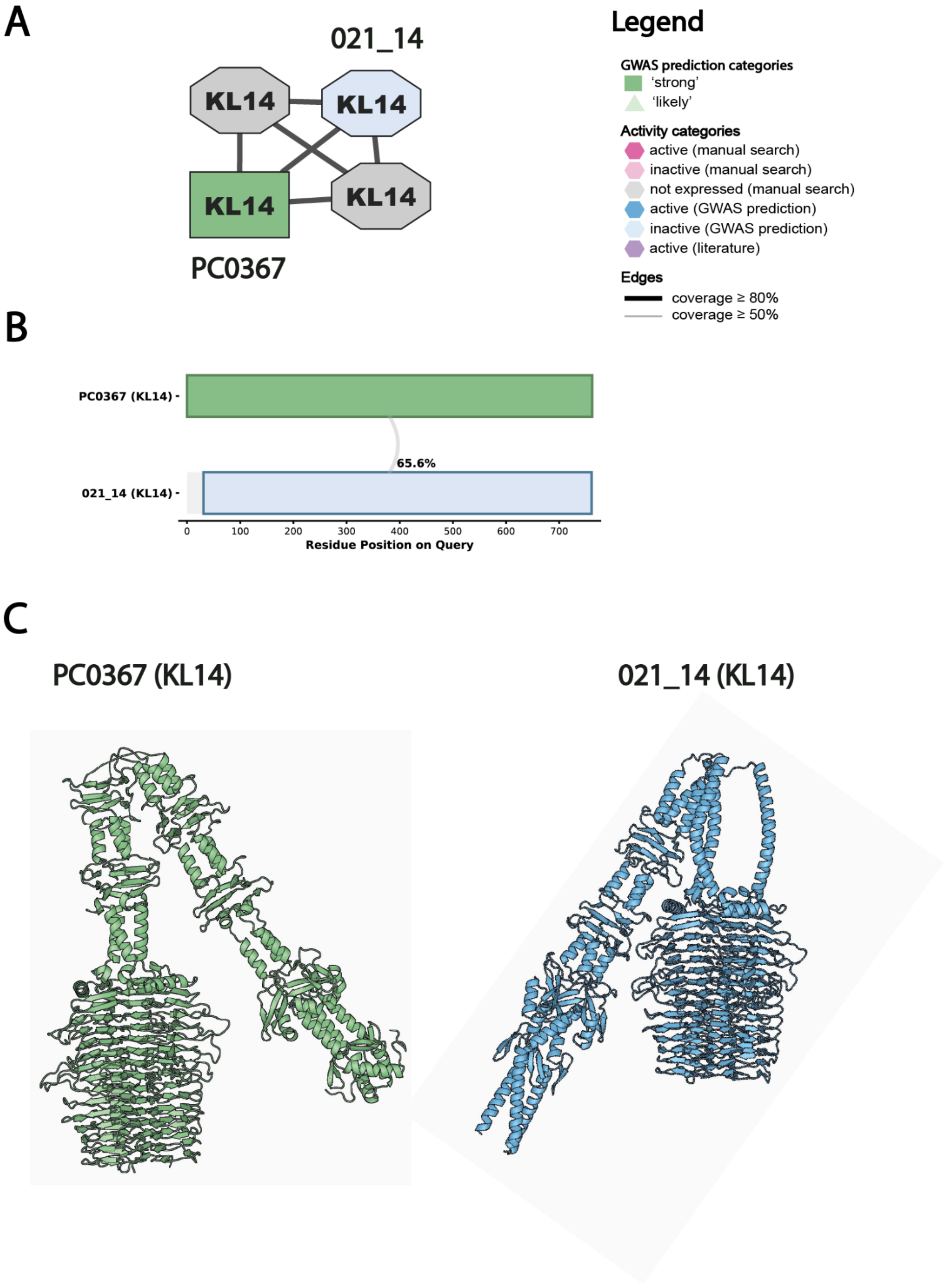
A
021_14
KL14
KL14
KL14
KL14
PC0367
Legend
GWAS prediction categories
'strong'
'likely'
Activity categories
active (manual search)
inactive (manual search)
not expressed (manual search)
active (GWAS prediction)
inactive (GWAS prediction)
active (literature)
Edges
coverage ≥ 80%
coverage ≥ 50%
B
PC0367 (KL14)
021_14 (KL14)
65.6%
0
100
200
300
400
500
600
700
Residue Position on Query
C
PC0367 (KL14)
021_14 (KL14)

**Figure 7. Analysis of the connected component with the KL14 prediction.** (A) Connected component of the sequence similarity containing 1 GWAS prediction: KL14 (PC0367; strong, dark green). A comparison of the representative sequences from this cluster revealed similarity substantial against 2 recombinant proteins, both unsuccessfully produced. We commercially produced a representative of the KL14 prediction (021_14; blue), which was inactive and is shown in the network. (B) Pairwise BLASTP alignments visualization of the query sequence (PC0367; KL14) against one commercially produced inactive protein. (C) Structural homotrimeric models generated by AlphaFold3 of the two protein sequences from panel B.

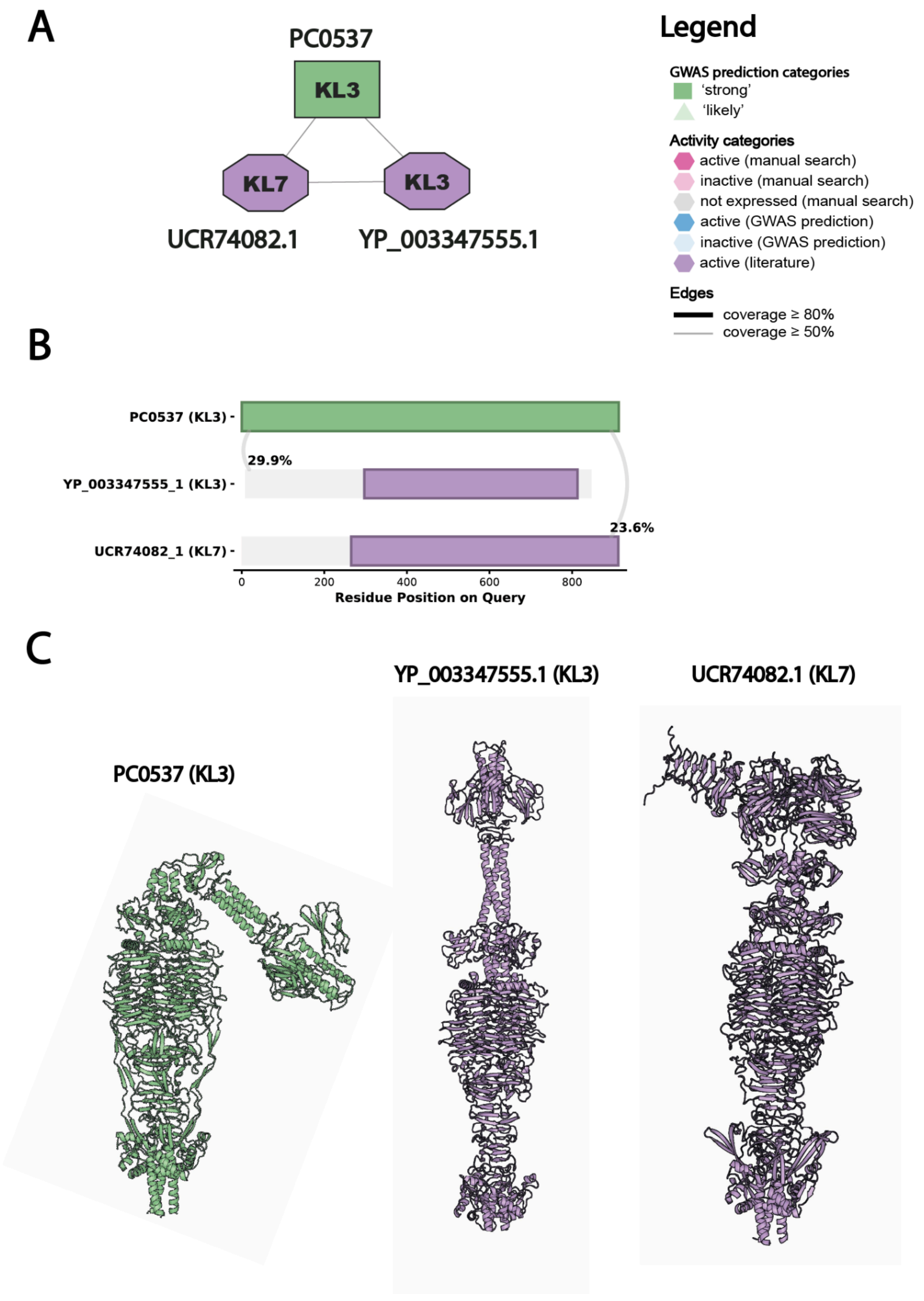
A
PC0537
KL3
KL7
KL3
UCR74082.1
YP_003347555.1
Legend
GWAS prediction categories
'strong'
'likely'
Activity categories
active (manual search)
inactive (manual search)
not expressed (manual search)
active (GWAS prediction)
inactive (GWAS prediction)
active (literature)
Edges
coverage ≥ 80%
coverage ≥ 50%
B
PC0537 (KL3)
YP_003347555_1 (KL3)
UCR74082_1 (KL7)
29.9%
23.6%
0
200
400
600
800
Residue Position on Query
C
YP_003347555.1 (KL3)
UCR74082.1 (KL7)
PC0537 (KL3)

**Figure 8. Analysis of the connected component with the KL3 prediction.** (A) Connected component of the sequence similarity containing 1 GWAS prediction: KL3 (PC0537; strong, dark green). A comparison of the representative sequence from this cluster revealed similarity substantial against 2 recombinant proteins, of which one was active against KL3 (YP_003347555.1; purple) and one was active against KL7 (URC74082.1; purple). (B) Pairwise BLASTP alignments visualization of the query sequence (PC0537; KL3) and two active proteins. (C) Structural homotrimeric models generated by AlphaFold3 of the three protein sequences from panel B. The TM-score of alignment between the RBD of PC0537 and  the RBD of (i) YP_003347555_1 is 0.75, and (ii) UCR74082.1 is 0.57.

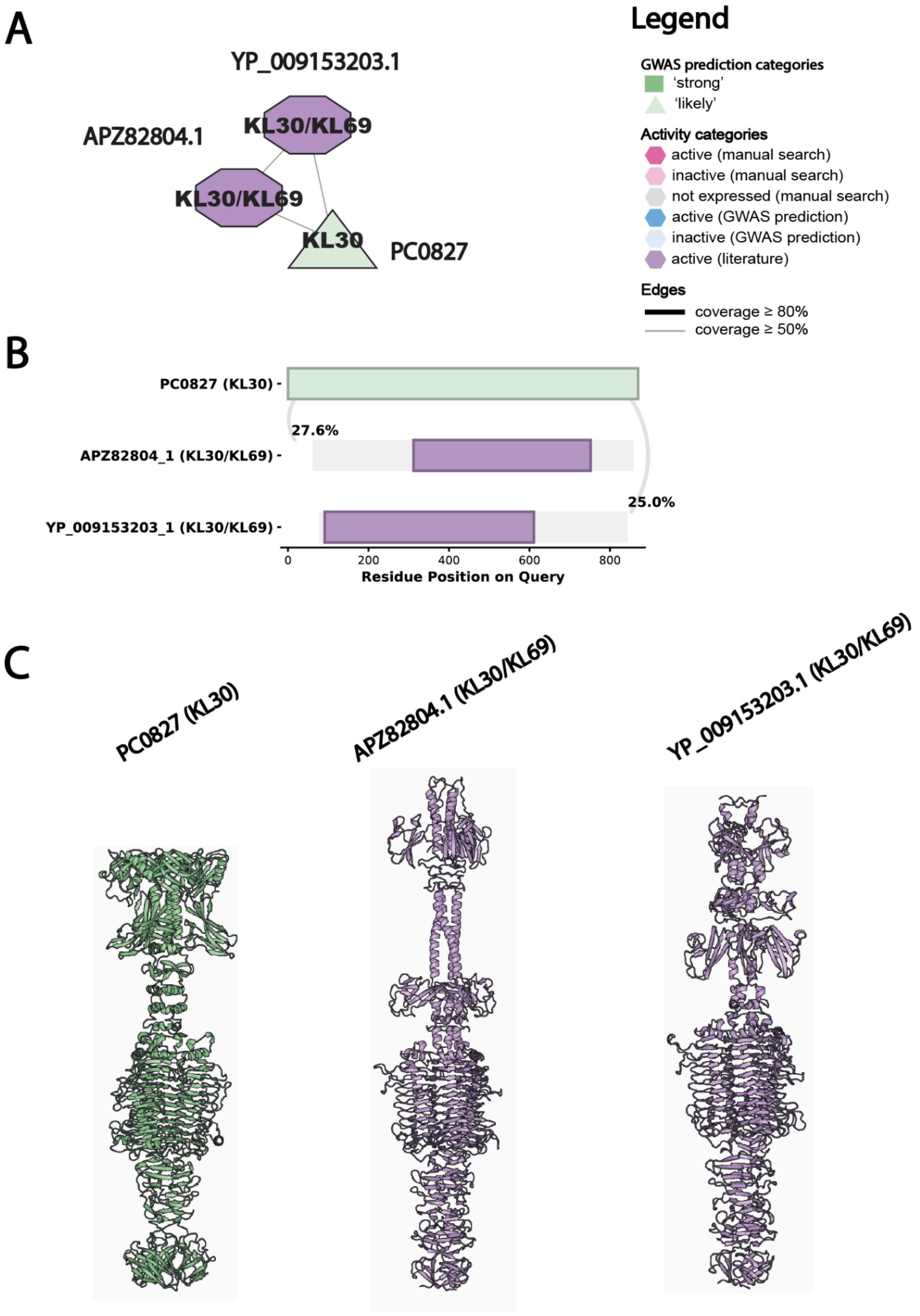
A
YP_009153203.1
APZ82804.1
KL30/KL69
KL30/KL69
KL30
PC0827
Legend
GWAS prediction categories
'strong'
'likely'
Activity categories
active (manual search)
inactive (manual search)
not expressed (manual search)
active (GWAS prediction)
inactive (GWAS prediction)
active (literature)
Edges
coverage ≥ 80%
coverage ≥ 50%
B
PC0827 (KL30)
27.6%
APZ82804_1 (KL30/KL69)
25.0%
YP_009153203_1 (KL30/KL69)
0
200
400
600
800
Residue Position on Query
C
PC0827 (KL30)
APZ82804.1 (KL30/KL69)
YP_009153203.1 (KL30/KL69)

**Figure 9. Analysis of the connected component with the KL30 prediction.** (A) Connected component of the sequence similarity containing 1 GWAS prediction: KL30 (PC0827; likely, light green). A comparison of the representative sequence from this cluster revealed similarity substantial against 2 recombinant proteins, both active and cross-specific against KL30 and KL69 (APZ82804.1; purple) (YP_009153203.1; purple). (B) Pairwise BLASTP alignments visualization of the query sequence (PC0827; KL30) against two active proteins. (C) Structural homotrimeric models generated by AlphaFold3 of the three protein sequences from panel B. The TM-score of alignment between the RBD of PC0827 and the RBD of: (i) APZ82804_1 is 0.65, and (ii) YP_009153203_1 is 0.65.

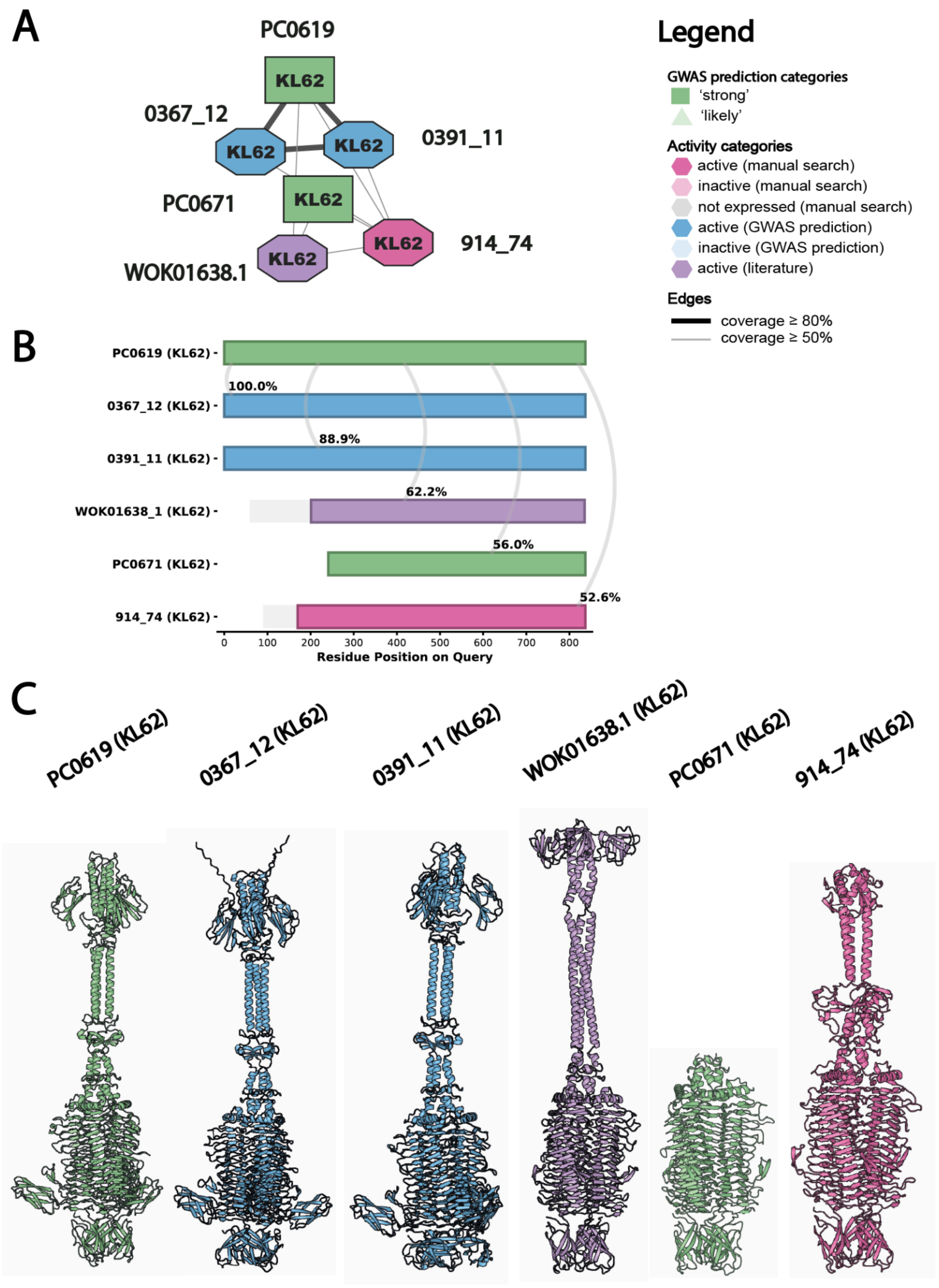
A
PC0619
KL62
0367_12
KL62
KL62
0391_11
PC0671
KL62
KL62
914_74
KL62
WOK01638.1
Legend
GWAS prediction categories
'strong'
'likely'
Activity categories
active (manual search)
inactive (manual search)
not expressed (manual search)
active (GWAS prediction)
inactive (GWAS prediction)
active (literature)
Edges
coverage ≥ 80%
coverage ≥ 50%
B
PC0619 (KL62)
0367_12 (KL62)
0391_11 (KL62)
WOK01638_1 (KL62)
PC0671 (KL62)
914_74 (KL62)
100.0%
88.9%
62.2%
56.0%
52.6%
0
100
200
300
400
500
600
700
800
Residue Position on Query
C
PC0619 (KL62)
0367_12 (KL62)
0391_11 (KL62)
WOK01638.1 (KL62)
PC0671 (KL62)
914_74 (KL62)

**Figure 10. Analysis of the connected component with the KL62 predictions.** (A) Connected component of the sequence similarity containing 2 GWAS predictions against: KL62 (PC0619; strong, dark green) and KL62 (PC0671; strong; dark green). A comparison of the representative sequence from these clusters revealed similarity substantial against 2 recombinant proteins, both active against KL62 (WOK01638.1; purple) (914_74; magenta). We also commercially produced two representatives of the KL62 (PC0619) prediction (0367_12 and 0391_11; blue), which were active and are shown in the network. (B) Pairwise BLASTP alignments visualization of the query sequence (PC0619; KL62) against other GWAS predictions and the four active proteins. (C) Structural homotrimeric models generated by AlphaFold3 of the six protein sequences from panel B. The TM-score of alignment between the RBD of PC0671 and the RBD of: (i) WOK01638_1 is 0.98, and (ii) 914_74 is 0.99.

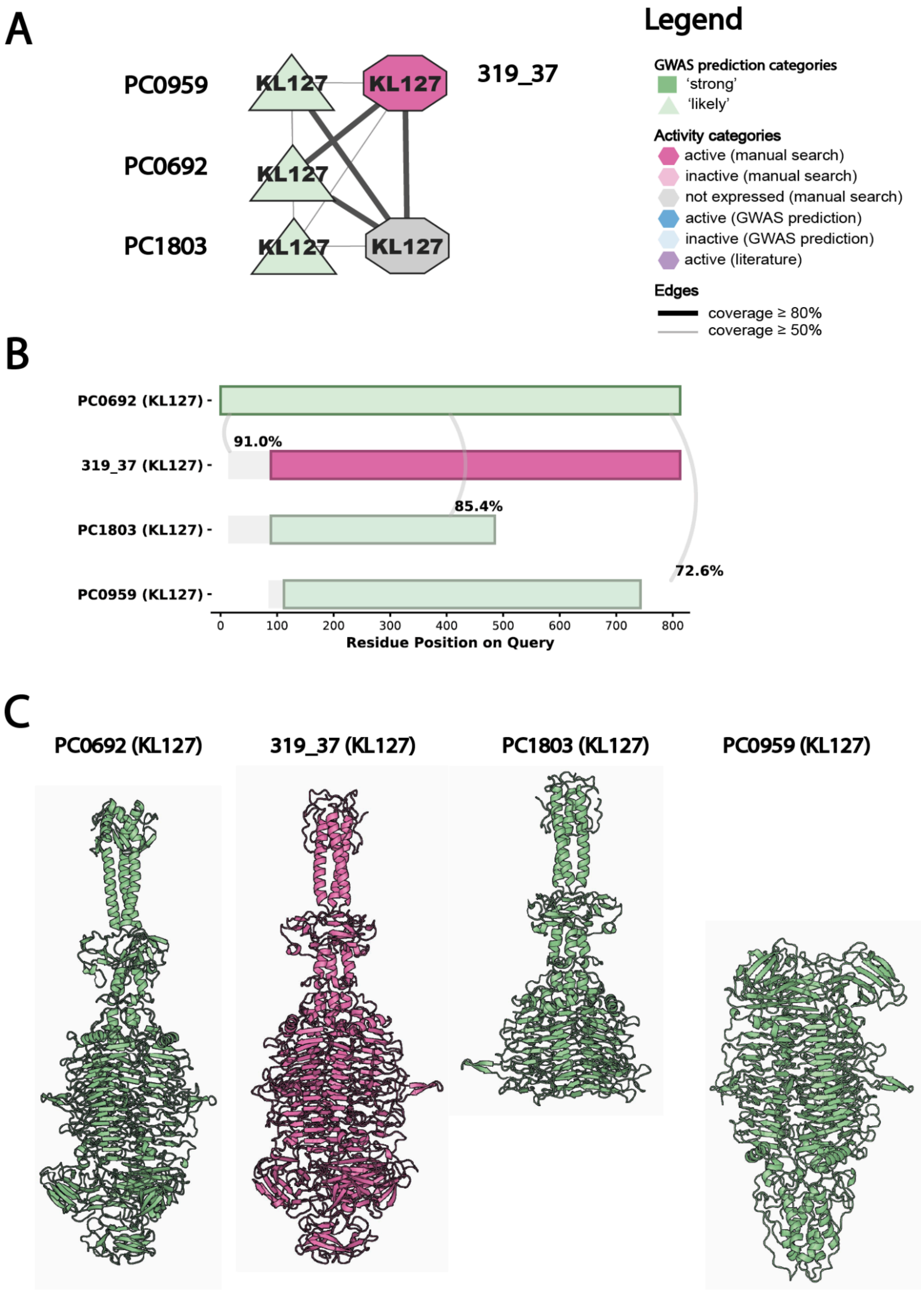

A
PC0959
PC0692
PC1803
KL127
319_37
Legend
GWAS prediction categories
'strong'
'likely'
Activity categories
active (manual search)
inactive (manual search)
not expressed (manual search)
active (GWAS prediction)
inactive (GWAS prediction)
active (literature)
Edges
coverage ≥ 80%
coverage ≥ 50%
B
PC0692 (KL127)
319_37 (KL127)
PC1803 (KL127)
PC0959 (KL127)
91.0%
85.4%
72.6%
0
100
200
300
400
500
600
700
800
Residue Position on Query
C
PC0692 (KL127)
319_37 (KL127)
PC1803 (KL127)
PC0959 (KL127)

**Figure 11. Analysis of the connected component with the KL127 predictions.** (A) Connected component of the sequence similarity containing 3 GWAS predictions all against KL127 (PC0959; likely, light green) (PC0692; likely; light green) (PC1803; likely; light green). A comparison of the representative sequences from these clusters revealed similarity substantial against 2 recombinant proteins, one active against KL127 (319_37; magenta) and second unsuccessfully produced. (B) Pairwise BLASTP alignments visualization of the query sequence (PC0692; KL127) against other GWAS predictions and one active protein. (C) Structural homotrimeric models generated by AlphaFold3 of the four protein sequences from panel B. The TM-score of alignment between the RBD of PC0692 and the RBD of 319_37 is 0.67 but these RBDs are nearly identical when aligned in pymol.

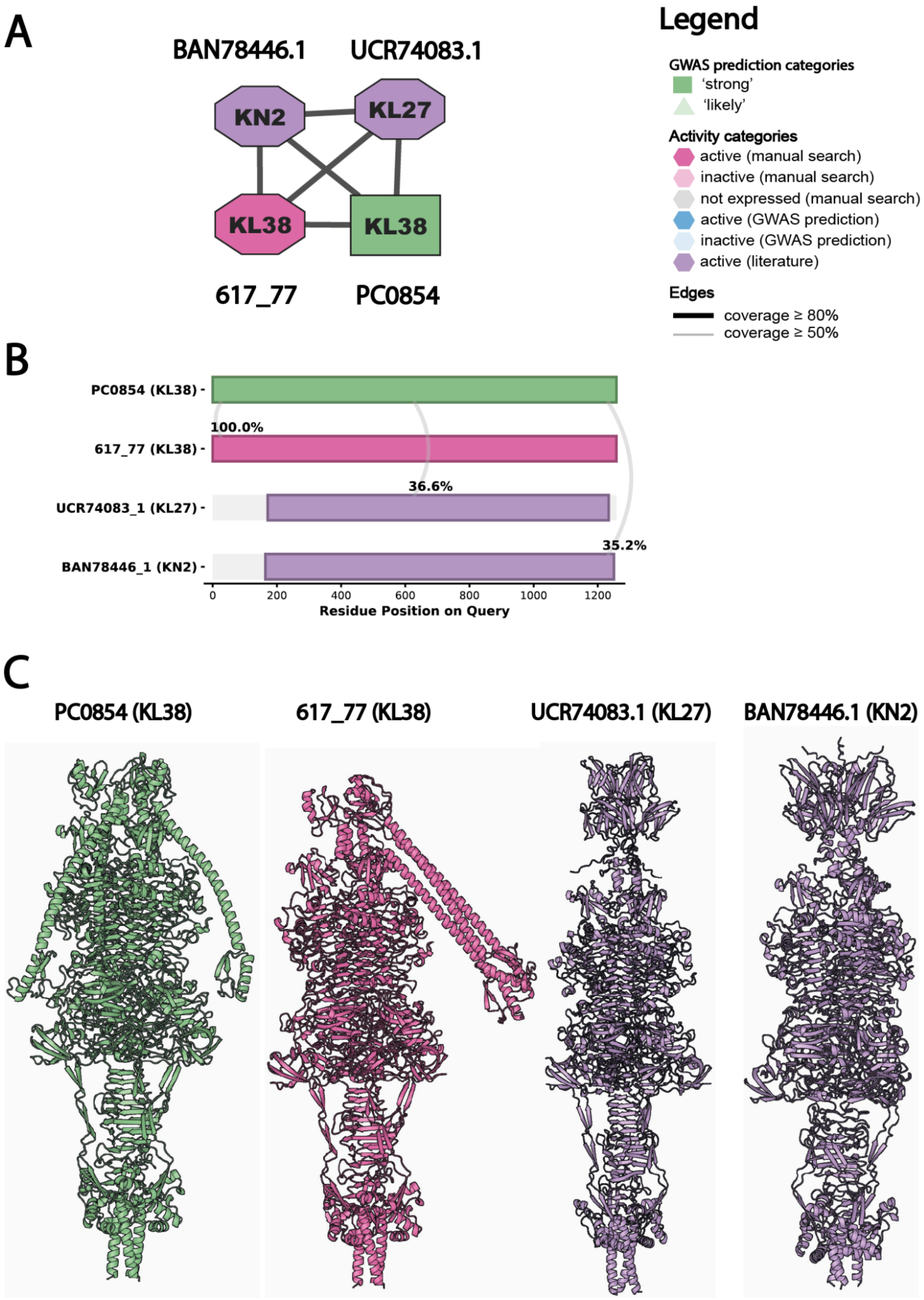
A
BAN78446.1
UCR74083.1
KN2
KL27
KL38
KL38
617_77
PC0854
Legend
GWAS prediction categories
'strong'
'likely'
Activity categories
active (manual search)
inactive (manual search)
not expressed (manual search)
active (GWAS prediction)
inactive (GWAS prediction)
active (literature)
Edges
coverage ≥ 80%
coverage ≥ 50%
B
PC0854 (KL38)
617_77 (KL38)
UCR74083_1 (KL27)
BAN78446_1 (KN2)
100.0%
36.6%
35.2%
0
200
400
600
800
1000
1200
Residue Position on Query
C
PC0854 (KL38)
617_77 (KL38)
UCR74083.1 (KL27)
BAN78446.1 (KN2)

**Figure 12. Analysis of the connected component with the KL38 prediction.** (A) Connected component of the sequence similarity containing 1 GWAS prediction against KL38 (PC0854; strong, dark green). A comparison of the representative sequence from this cluster revealed similarity substantial against 3 recombinant proteins of which, one is active against KL38 (617_77; magenta), one active against KL27 (UCR74083.1, purple) and one active against KN2 (BAN78446.1, purple). (B) Pairwise BLASTP alignments visualization of the query sequence (PC0854; KL38) against the three active proteins. (C) Structural homotrimeric models generated by AlphaFold3 of the four protein sequences from panel B. The TM-score of alignment between the RBD of PC0854 and the RBD of 617_77 is 0.99.
