## Supplementary Figures for "Capsular specificity in temperate phages of *Klebsiella pneumoniae* is driven by diverse receptor-binding enzymes"

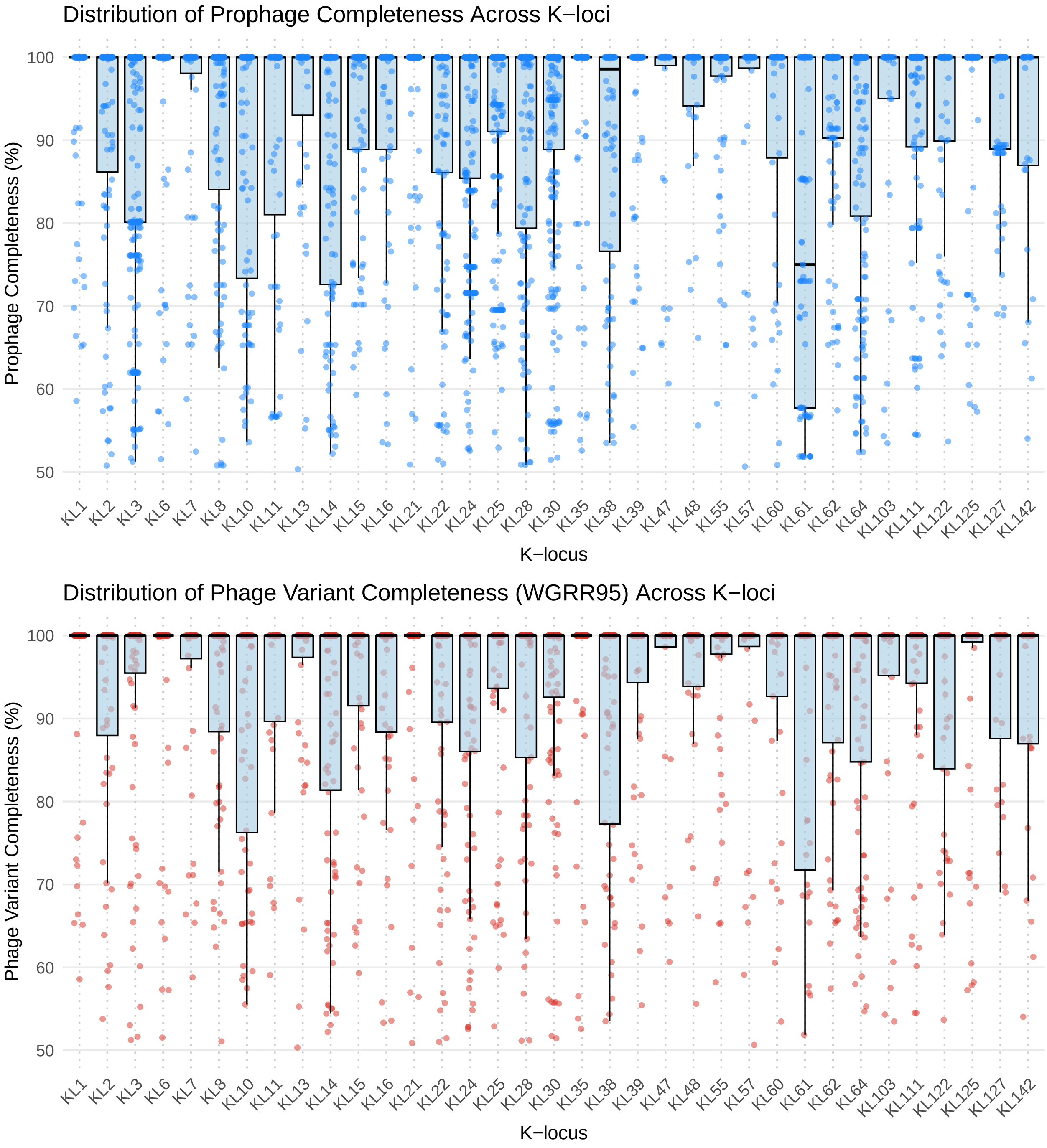


**Figure S1.** (Top) Distribution of prophage completeness as estimated by CheckV per K-type for the 35 K-types shown in Figure 1. Each point shows the completeness of each prophage detected in an isolate with the corresponding K-type. Box-plots are overlaid to show the median values and quantiles. (Bottom) The same but shown for a single representative per phage variant with the highest value of completeness.

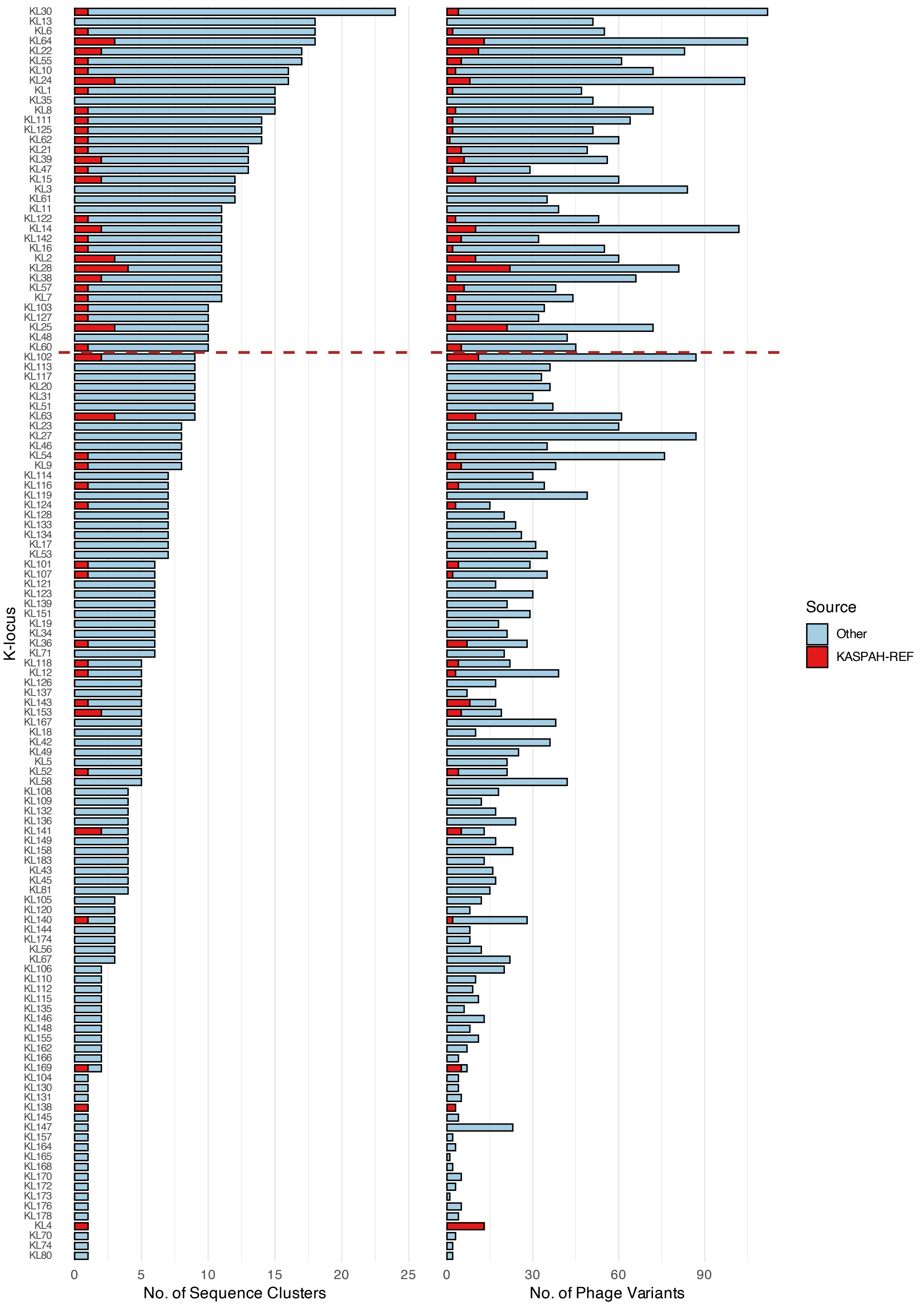


**Figure S2.** (Left) Full distribution of the number of Sequence Clusters (SCs) per K-type for all K-types in the dataset from Figure 1. (Right) Full distribution of the number of phage variants, obtained at wGRR=0.95, per K-type for all K-types in the dataset.

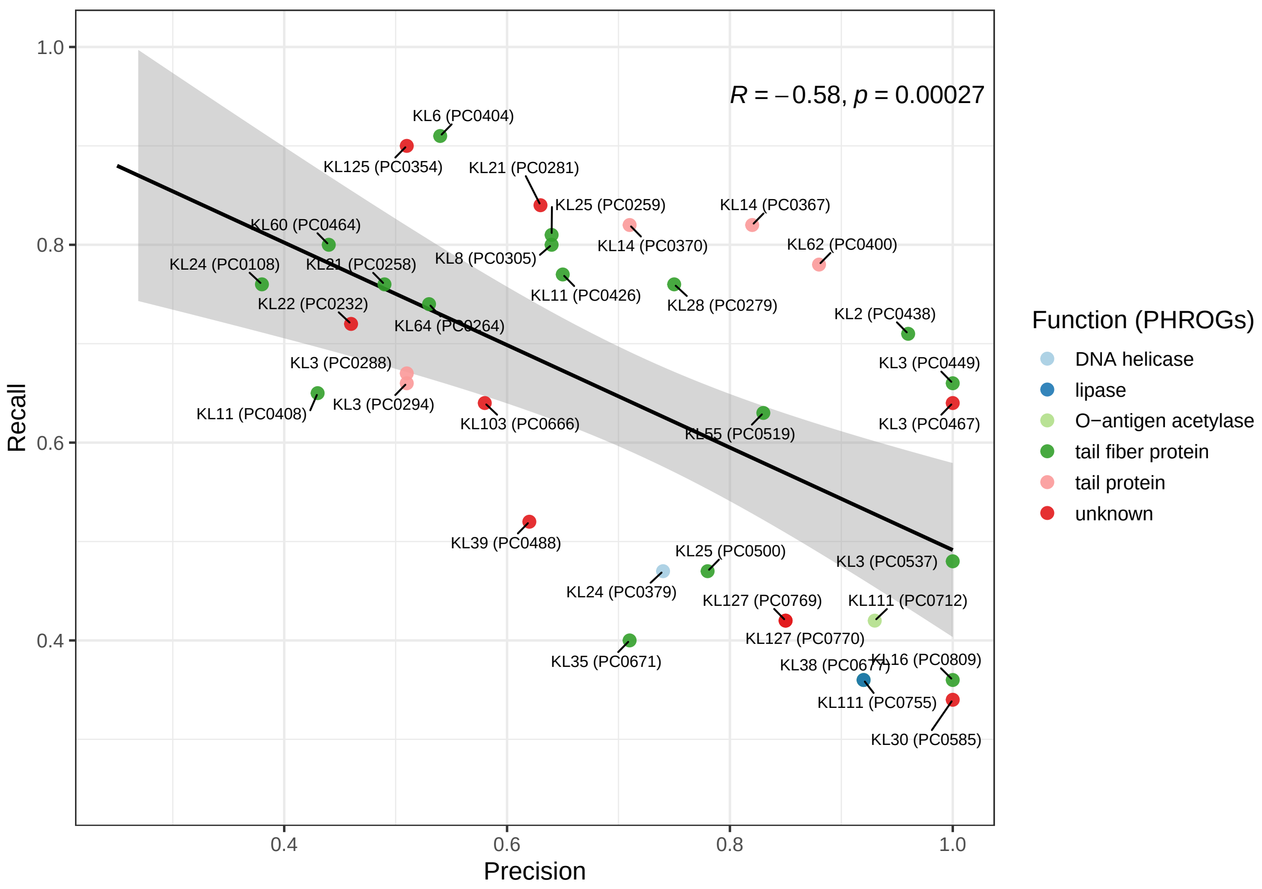


**Figure S3.** Relationship between Precision and Recall for the 35 genetic variants from Fig. 2A, i.e., those with F1 ⩾ 0.5 and MCC ⩾ 0.5. The black line shows the best-fit linear model (Pearson correlation) to these data. Colours indicate the corresponding functional predictions obtained using PHROGs. The Pearson correlation coefficient and associated p-value are shown in the plot.

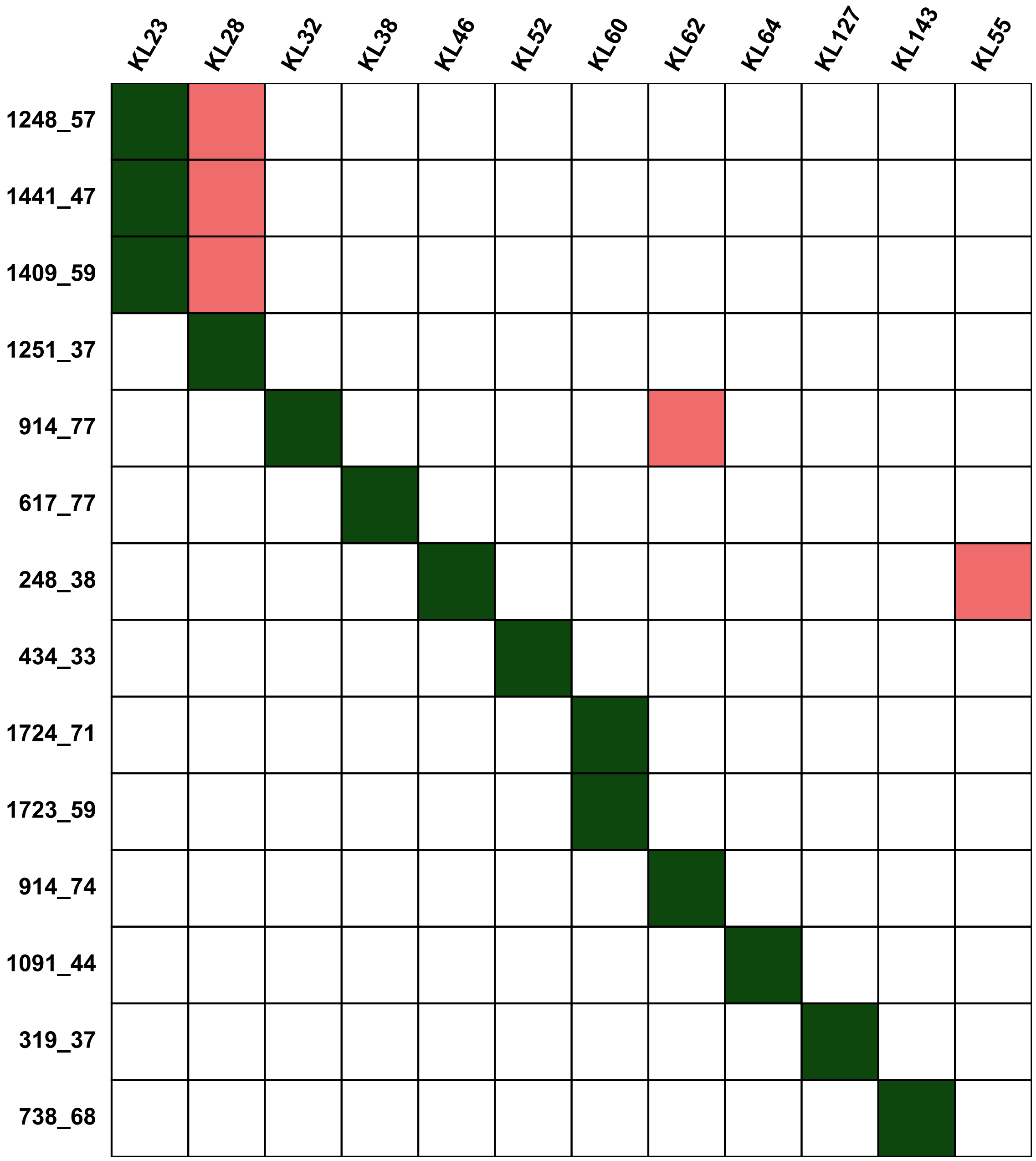


**Figure S4.** Each row represents one of the 14 recombinant proteins from Figure 4 that showed enzymatic activity against a panel of 119 reference K-types. Columns indicate K-loci corresponding to the bacterial hosts of the prophages from which the proteins originated. Green cells denote K-loci on which the enzyme was active; pink cells indicate the K-locus of the host from which the enzyme's prophage was derived. Rows with only green cells indicate activity against the host's own K-locus, while rows with both pink and green highlight activity on a different K-locus than the prophage's host.

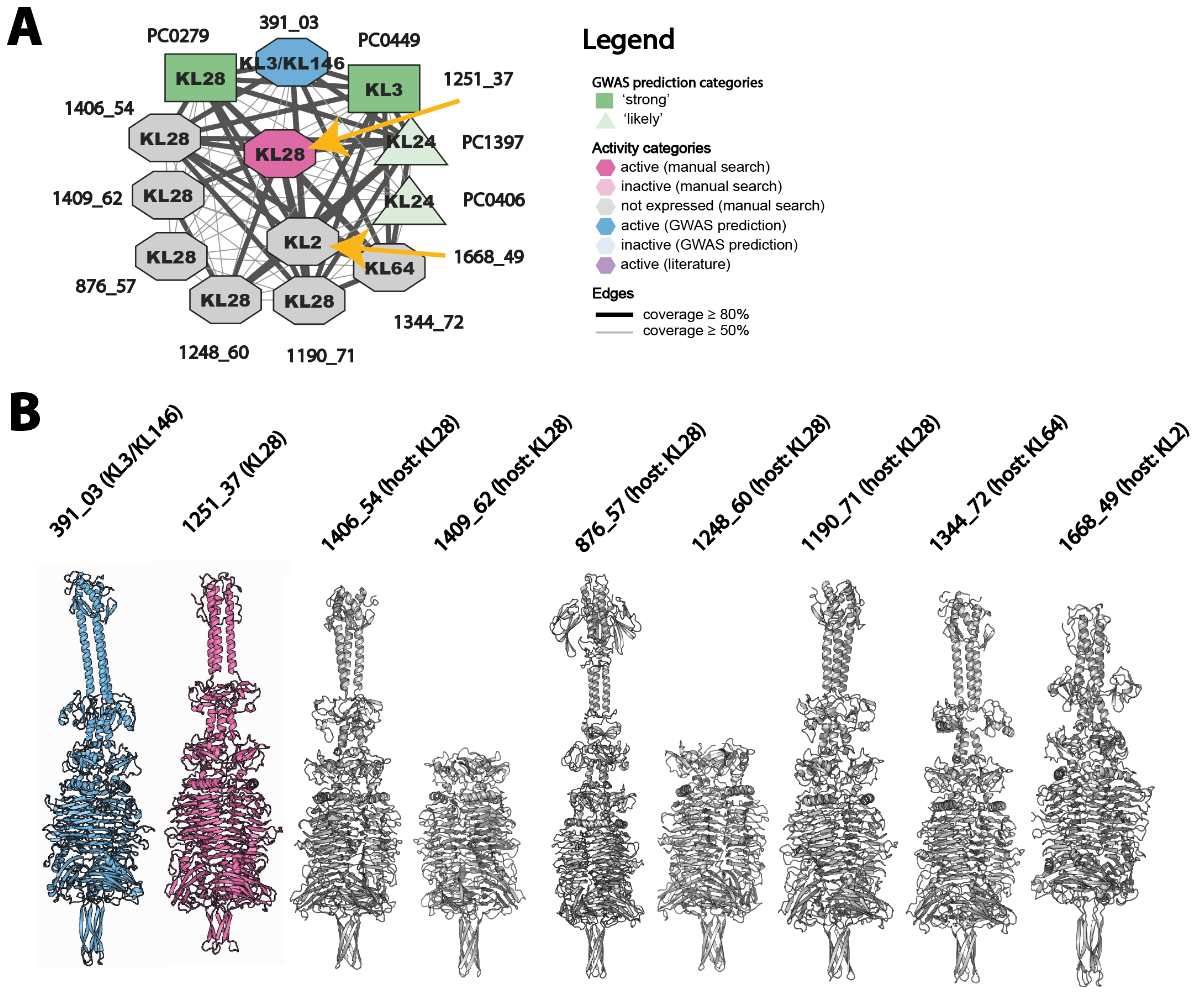


**C**

| proteinID | prophageID | prophage_completeness | prophage_confidence | ST | K_locus | K_locus_confidence |
|---|---|---|---|---|---|---|
| 391_03 | KPN_B3_PHAGE391_M | 98.17 | medium | ST896 | KL3 | Very high |
| 1251_37 | KPN_B19_PHAGE043_M | 100.0 | high | ST20 | KL28 | Very high |
| 1406_54 | KVV_B2_PHAGE063_M | 100.0 | high | ST925 | KL28 | Good |
| 1409_62 | KVV_B2_PHAGE067_M | 100.0 | high | ST925 | KL28 | Good |
| 876_57 | KPN_B20_PHAGE149_M | 100.0 | high | ST20 | KL28 | Very high |
| 1248_60 | KPN_B19_PHAGE044_M | 100.0 | high | ST20 | KL28 | Very high |
| 1190_71 | KPN_B3_PHAGE143_M | 100.0 | high | ST661 | KL28 | Very high |
| 1344_72 | KPN_B2_PHAGE046_M | 100.0 | high | ST20 | KL64 | Very high |
| 1668_49 | KPN_B3_PHAGE386_M | 100.0 | medium | ST27 | KL2 | High |

**Figure S5.** Only a fraction of predicted depolymerases can be overexpressed. Analysis of recombinant depolymerases from connected component 1 in the sequence similarity network (manuscript Figure 4) that did not overexpress. (A) Connected component 1 from the sequence similarity network of recombinant and predicted depolymerases (B) AlphaFold3 homotrimer models of proteins from A which were active or did not overexpress. (C) CheckV and Kaptive metadata for prophages and bacterial K-loci from which the genes encoding putative depolymerase were cloned.

.

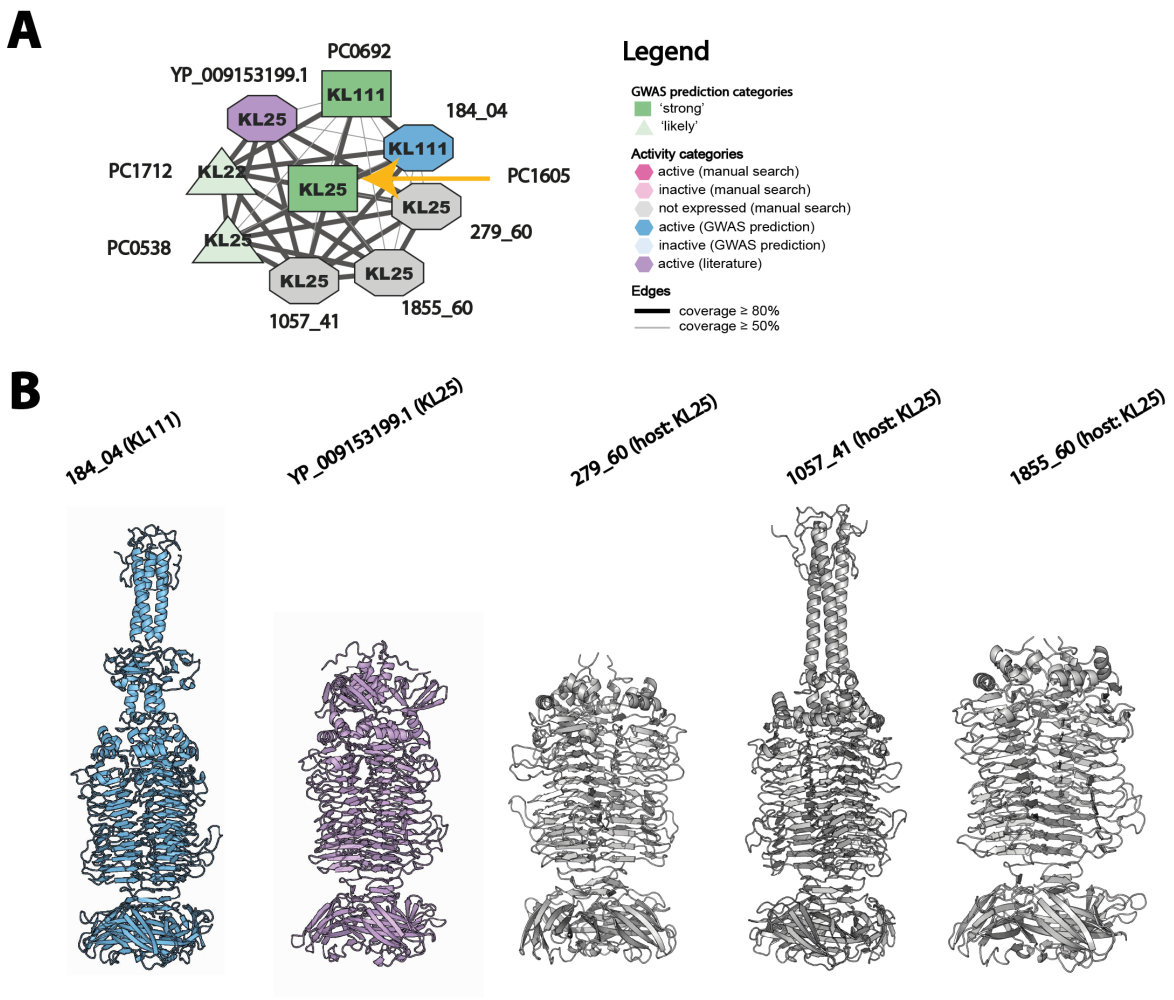


C

| proteinID | prophageID | prophage_completeness | prophage_confidence | ST | K_locus | K_locus_confidence |
|---|---|---|---|---|---|---|
| 184_04 | KPN_B9_PHAGE184_M | 100.0 | high | ST17 | KL111 | Very high |
| YP_009153199.1 | n/a | n/a | n/a | n/a | n/a | n/a |
| 279_60 | KPN_B20_PHAGE101_M | 100.0 | high | ST17 | KL25 | High |
| 1057_41 | KPN_B9_PHAGE130_M | 100.0 | high | ST461 | KL25 | High |
| 1855_60 | KPN_B20_PHAGE101_M | 100.0 | high | ST17 | KL25 | High |

**Figure S6.** Only a fraction of predicted depolymerases can be overexpressed. Analysis of recombinant depolymerases from connected component 4 in the sequence similarity network (manuscript Figure 4) that did not overexpress. (A) Connected component 4 from the sequence similarity network of recombinant and predicted depolymerases. (B) AlphaFold3 homotrimer models of proteins from A which were active or did not overexpress. (C) CheckV and Kaptive metadata for prophages and bacterial K-loci from which the genes encoding putative depolymerase were cloned.

.

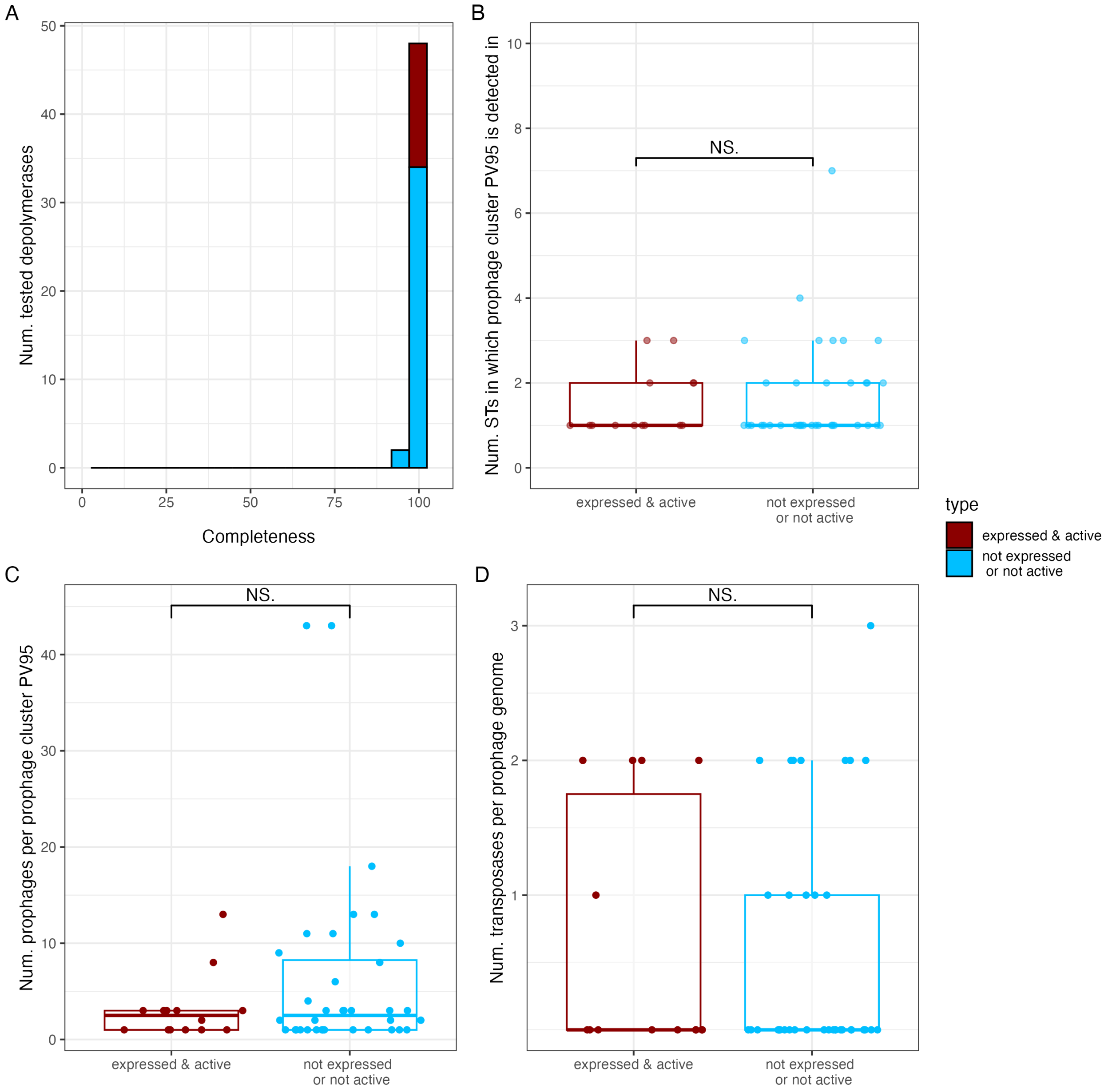


**Figure S7.** Comparison of prophage quality and genomic features between active and inactive depolymerases. We tested whether the lack of detectable activity or expression could be explained by the genomic context of the corresponding prophages. Each depolymerase was classified as expressed & active (dark red) or not expressed or not active (light blue), and the prophage of origin was analysed for several genomic attributes. (A) Prophage completeness estimated by CheckV. (B) Number of *K. pneumoniae* sequence types (STs) containing closely related prophages, used as a proxy for prophage age and vertical inheritance. (C) Size of the prophage cluster (number of similar prophages in the dataset), reflecting the frequency of the element across hosts. (D) Number of annotated transposases per prophage genome, a hallmark of prophage degradation. All tested prophages were complete or near-complete, and no significant differences were observed between active and inactive groups for any feature (Fisher's exact and Wilcoxon tests, p ¿ 0.1). These analyses indicate that inactivity or low expression cannot be explained by prophage degradation or domestication.

.

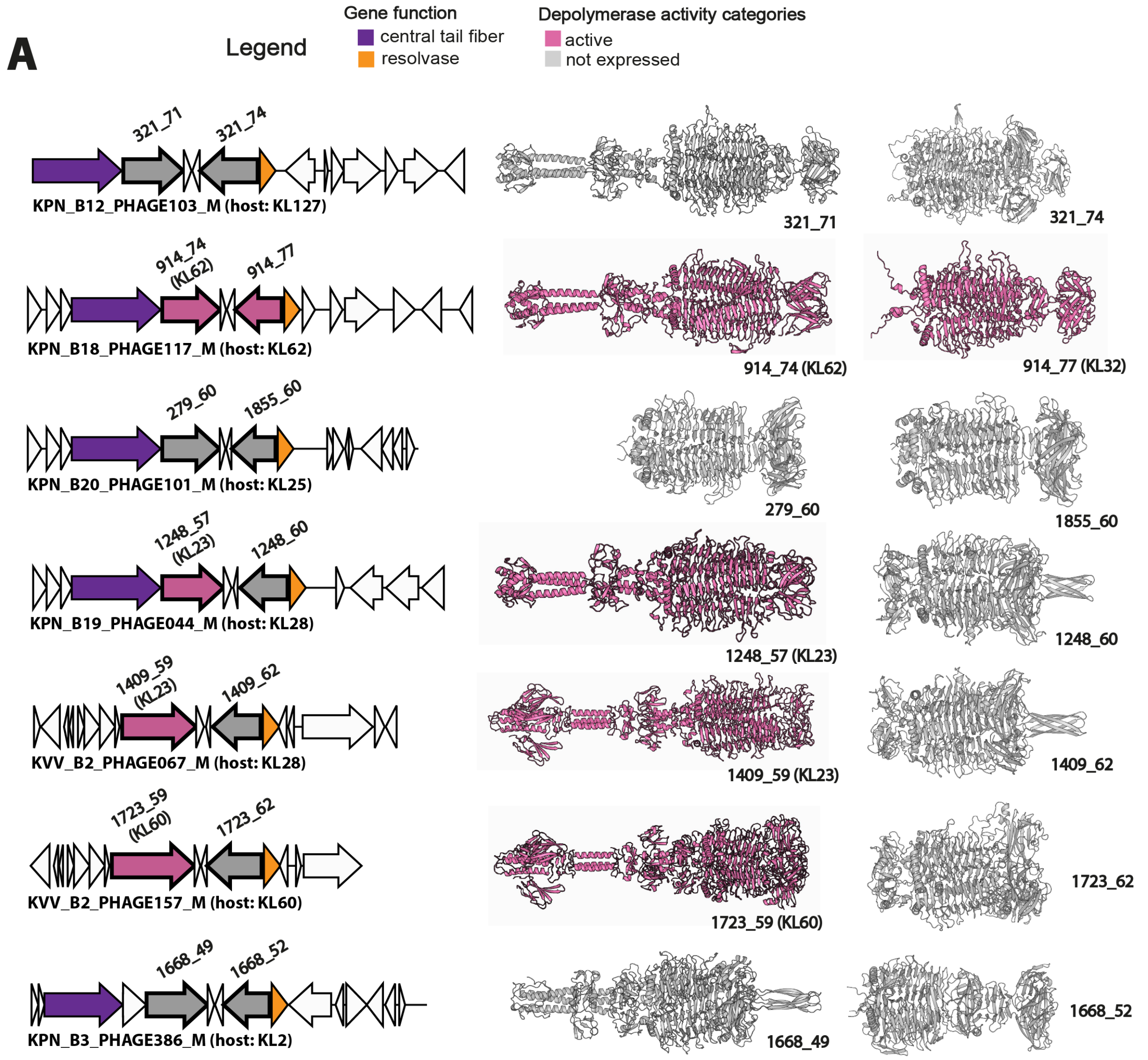


**B**

| proteinID | prophageID | prophage_completeness | prophage_confidence | ST | K_locus | K_locus_confidence |
|---|---|---|---|---|---|---|
| 321_71 | KPN_B12_PHAGE103_M | 100.0 | high | ST230 | KL127 | Very high |
| 321_74 | KPN_B12_PHAGE103_M | 100.0 | high | ST230 | KL127 | Very high |
| 914_74 | KPN_B18_PHAGE117_M | 100.0 | high | ST2513 | KL62 | Very high |
| 914_77 | KPN_B18_PHAGE117_M | 100.0 | high | ST2513 | KL62 | Very high |
| 279_60 | KPN_B20_PHAGE101_M | 100.0 | high | ST17 | KL25 | High |
| 1855_60 | KPN_B20_PHAGE101_M | 100.0 | high | ST17 | KL25 | High |
| 1248_57 | KPN_B19_PHAGE044_M | 100.0 | high | ST20 | KL28 | Very high |
| 1248_60 | KPN_B19_PHAGE044_M | 100.0 | high | ST20 | KL28 | Very high |
| 1409_59 | KVV_B2_PHAGE067_M | 100.0 | high | ST925 | KL28 | Good |
| 1409_62 | KVV_B2_PHAGE067_M | 100.0 | high | ST925 | KL28 | Good |
| 1723_59 | KVV_B2_PHAGE157_M | 100.0 | high | ST641 | KL60 | Very high |
| 1723_62 | KVV_B2_PHAGE157_M | 100.0 | high | ST641 | KL60 | Very high |
| 1668_49 | KPN_B3_PHAGE386_M | 100.0 | medium | ST27 | KL2 | High |
| 1668_52 | KPN_B3_PHAGE386_M | 100.0 | medium | ST27 | KL2 | High |

**Figure S8.** Prophages carrying multiple depolymerases from the manual search. (A) Prophage tail gene modules from which $\geqslant 2$ putative depolymerase genes were cloned and tested along with their AlphaFold3 homotrimer models. Colours correspond to active depolymerases (pink), putative depolymerases which did not overexpress (grey), central tail fiber (violet), lysozyme (red), resolvase (orange). (B) CheckV and Kaptive metadata for prophages and bacterial K-loci loci from which the genes encoding putative depolymerase were cloned.

.

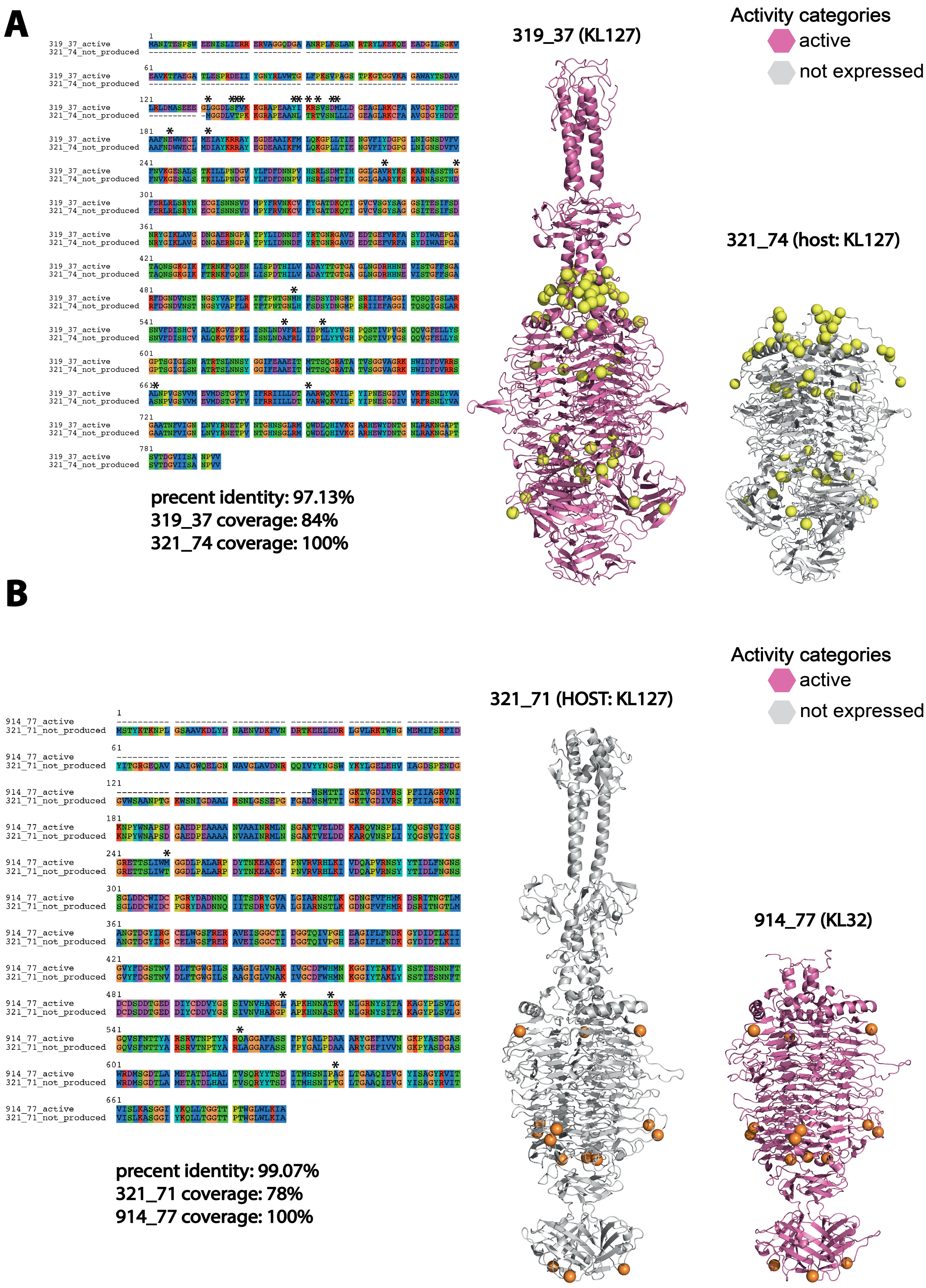


**Figure S9.** Similarity of active and not produced depolymerases. Two examples of protein pairs selected for recombinant overexpression and enzymatic activity testing (A, B). For each pair, the receptor-binding domains share >97% percentage identity in amino-acid sequence, and the corresponding AlphaFold3 homotrimer models are shown. Amino-acid substitutions between proteins in a pair are mapped onto the models and displayed as yellow/orange balls (on each monomer), showing their distribution across multiple parts of the structures.

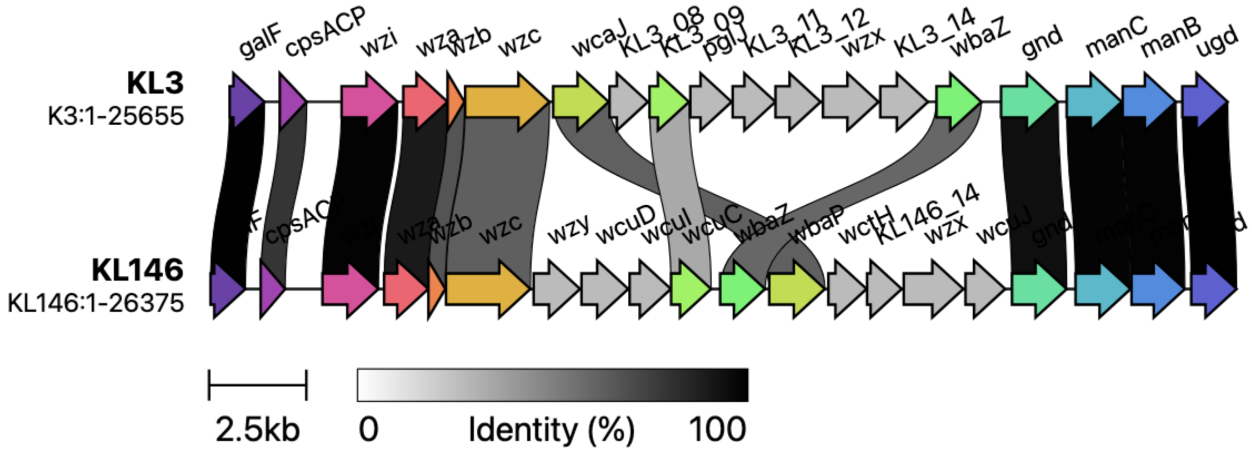


**Figure S10.** Comparison of the K-loci KL3 and KL146 using Clinker. Each arrow denotes a coding gene, shaded areas show similarity between protein sequences of genes in both loci, and opacity denotes identity weighted by gene overlap. K-locus sequences were obtained from Kaptive.

.

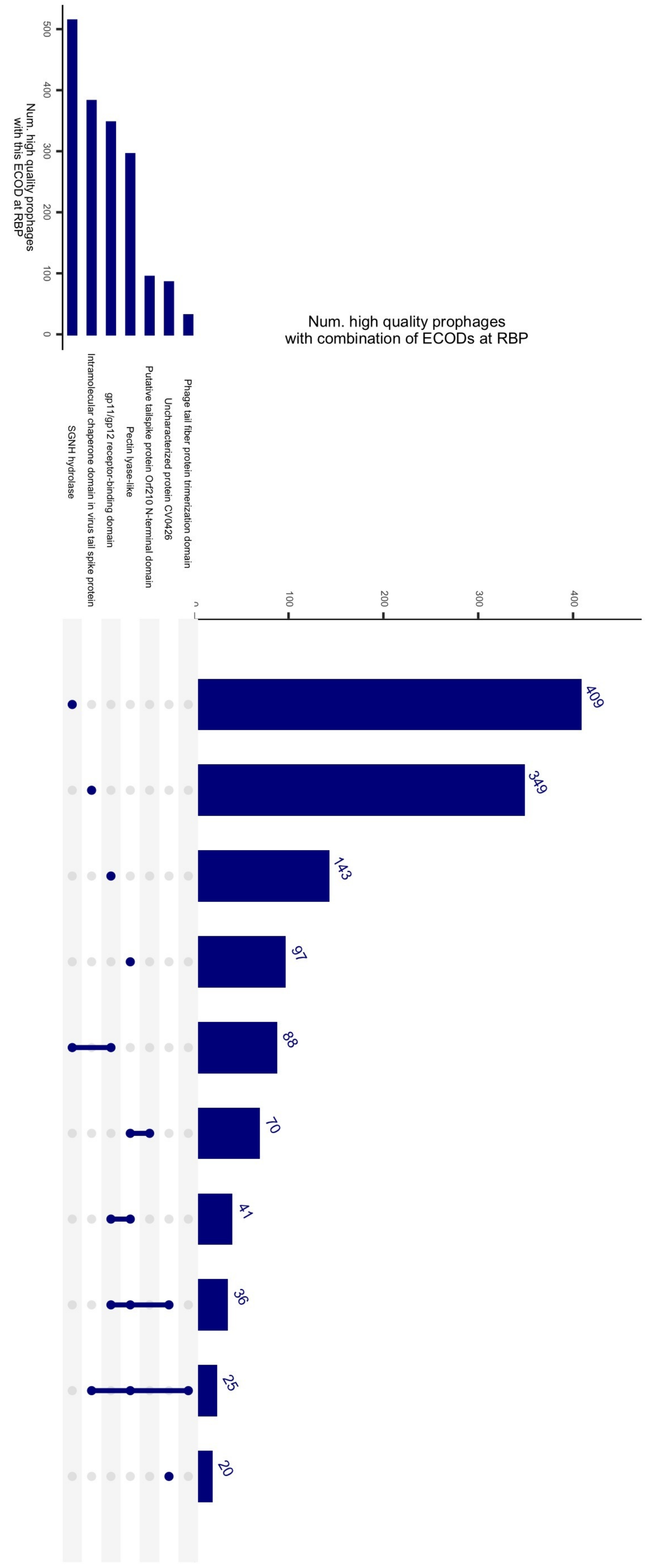


**Figure S11.** Upset plot showing the number of high-quality (at least 99% complete) prophages encoding each ECOD domain or combination of ECOD domains shown in Fig 5A within PHROG-defined RBPs, stratified by K-locus. The most frequent multi-domain combinations are gp11/gp12 + SGNH hydrolase and pectin-lyase-like + Orf210 N-terminal or gp11/gp12 domains.

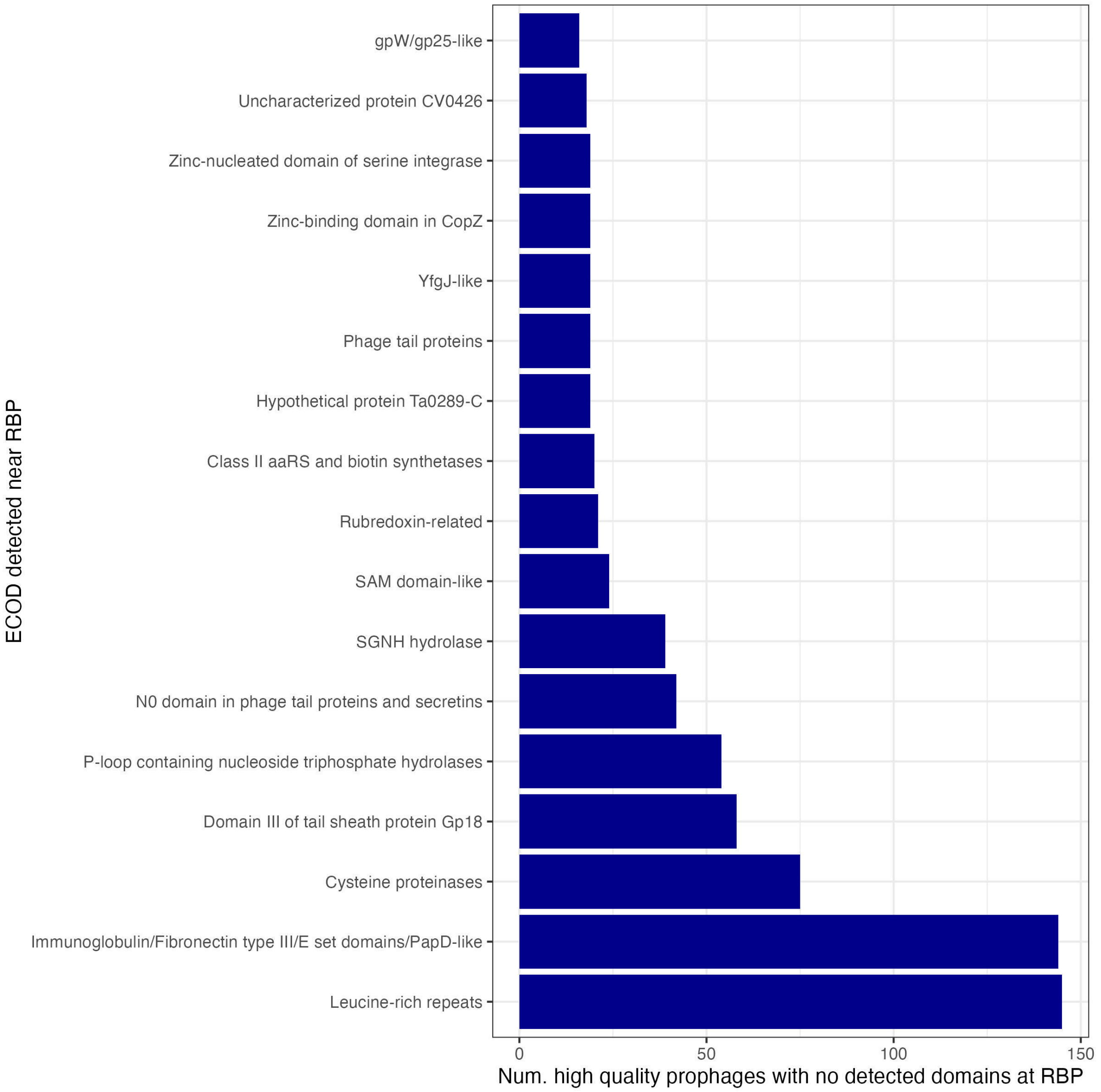


**Figure S12.** Number of high-quality (at least 99% complete) prophages whose RBPs lack any of the domains shown in Fig 5A but encode each ECOD domain up to 2 protein-coding genes upstream or downstream from the protein annotated as RBP.

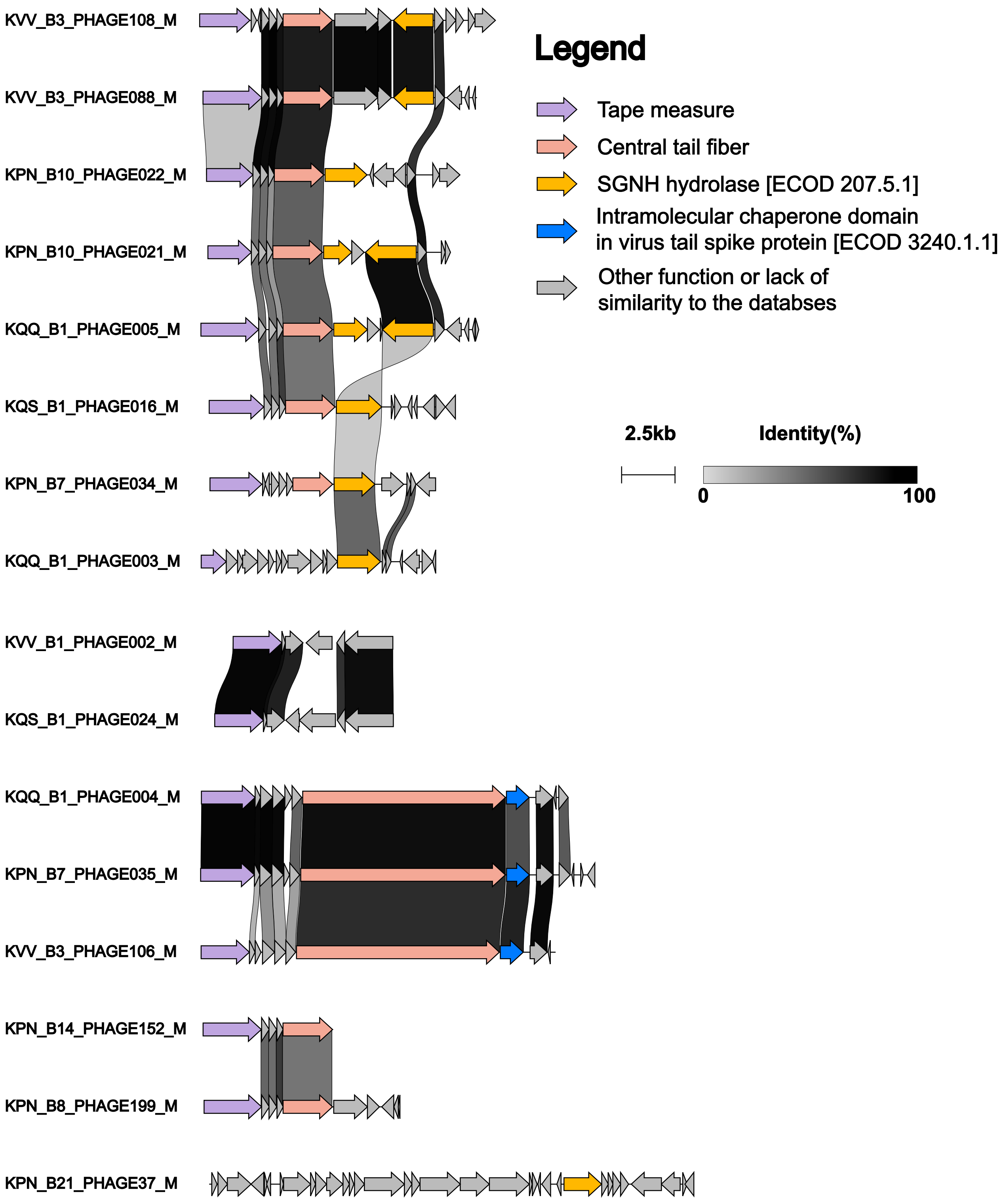


**Figure S13.** Tail regions of representative 16 prophages found in *K. pneumoniae* isolates with the KL1 locus. These prophages were chosen from all high-quality (100% completeness and 'high' confidence) prophages in KL1 isolates by accounting for bacterial lineage (SC) and phage variant (wGRR = 0.5 threshold). For clarity, prophage genomes were manually curated to highlight tail gene clusters by removing genes located downstream of the tail length tape measure protein. For two prophages (KVV_B1_PHAGE002_M and KQS_B1_PHAGE024_M) genes upstream of the tail tube protein were also removed. Proteins were annotated by the ECOD with a e-value of $10^{-3}$. Tail regions were compared using Clinker, and the protein identity is indicated as in Supplementary Figure S5.

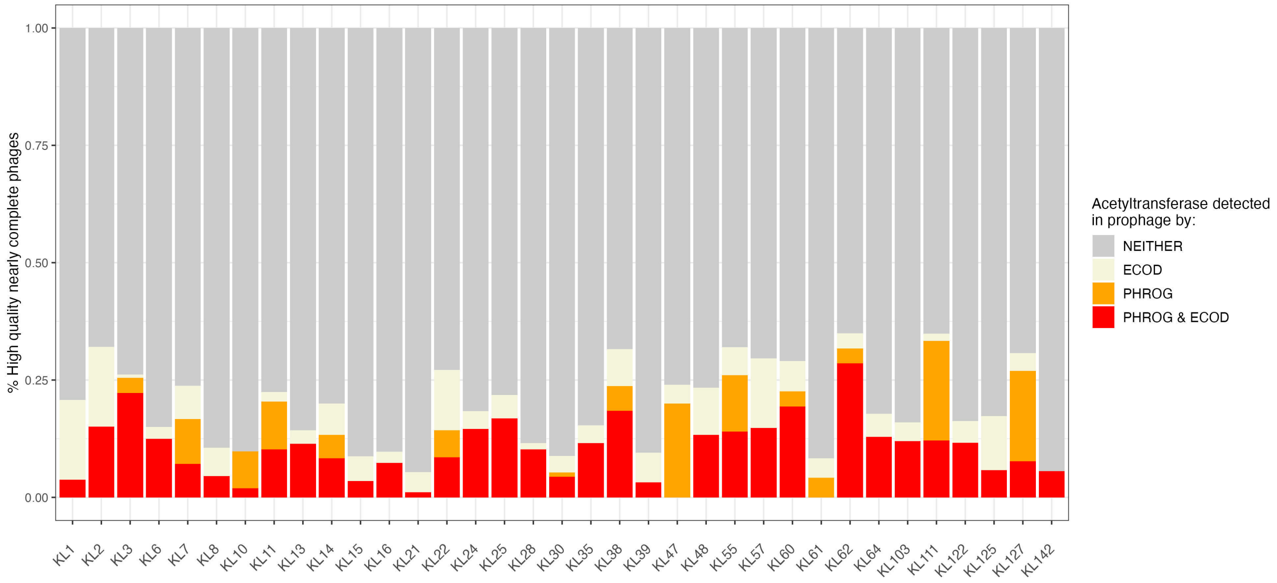


**Figure S14.** Number of high-quality (at least 99% complete) prophages that encode acetyltransferases as detected by PHROG (hit to ‘acetyltransferase’ or ‘O-acetyltransferase’) with prob $\geqslant 0.9$ and qcov/scov $\geqslant 0.5$, or by ECOD (hit to any ECOD containing a string `acetyltransf` with prob $\geqslant 0.9$ and scov $\geqslant 0.1$. Yellow shows hits to ECOD-only, orange shows hits to PHROG only, red shows hits to both and grey shows this to neither database.

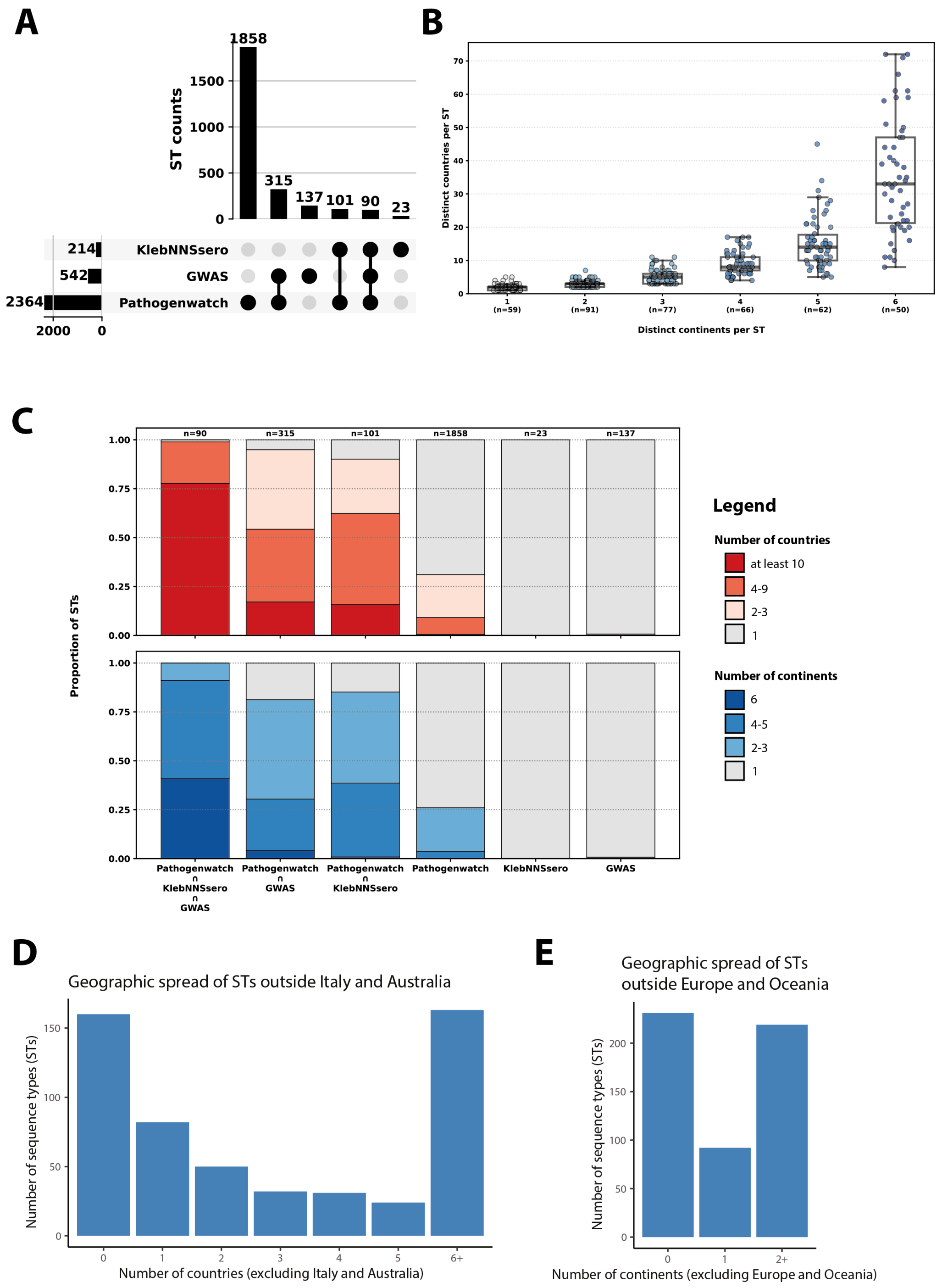


**Figure S15. Global distribution of *K. pneumoniae* sequence types (STs) across the GWAS, KlebNNSsero and Pathogenwatch datasets.** (A) Overlap in *K. pneumoniae* STs between the three datasets, showing shared and dataset-unique ST subsets. (B) Jitter plot with boxplots showing, for each of the 405 shared STs, the number of countries in which the ST has been detected (Y-axis) plotted against the number of continents where it occurs (X-axis). (C) Stacked bar plots illustrating the country and continent spread of STs within each subset, normalised by the total number of STs in that subset. (D) Bar plot showing the number of sequence types (STs; $n = 542$) in the GWAS dataset (y-axis) as a function of the number of countries in which they have been reported (x-axis), excluding the two GWAS source countries (Italy and Australia). (E) Analogous bar plot showing the number of STs as a function of the number of continents in which they have been reported, excluding the two GWAS source continents (Europe and Oceania). Across the 542 STs in the GWAS dataset, 390 STs (72%) have been reported in at least two countries and 314 STs (58%) in at least three countries, while 347 STs (64%) have been reported on at least two continents. Notably, 382 STs (70%) were detected in at least one additional country beyond Italy and Australia, and 311 STs (57%) were detected on at least one additional continent beyond Europe and Oceania. Together, these results indicate that the majority of lineages represented in the GWAS dataset are globally distributed.

.

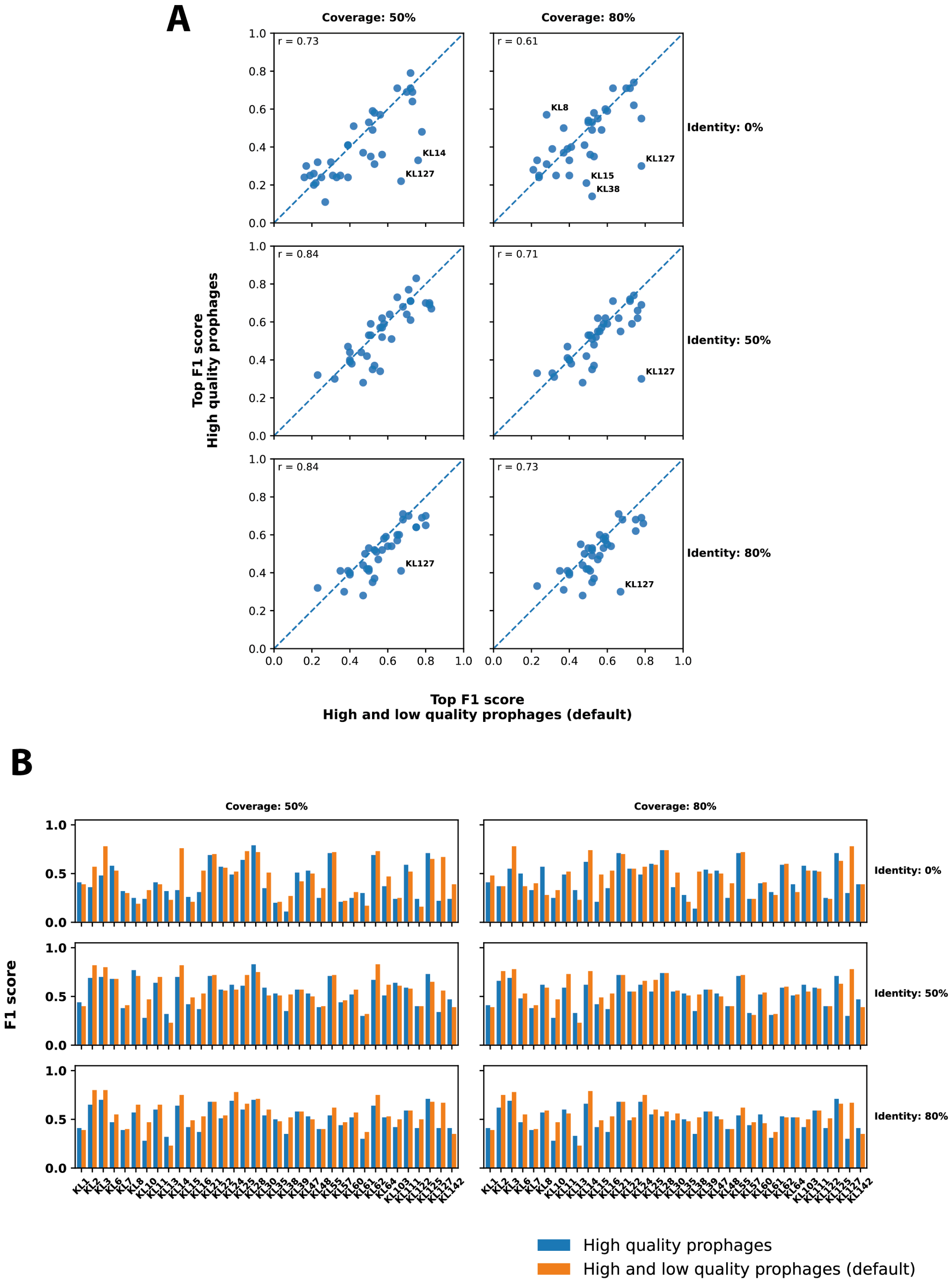


**Figure S16.** Effect of low-quality prophages on GWAS predictions. Low-quality prophages were defined as CheckV completeness <99% or medium/low CheckV confidence; high-quality prophages had completeness ⩾99% and high confidence. (A) Scatter plots compare, for 35 K-loci, the top F1 scores obtained from GWAS runs on 2 prophage datasets, proteins from high- and low-quality prophages (X-axis) versus proteins from high-quality prophage only (Y-axis), and six protein clustering settings. Each point represents a K-locus; row and column labels correspond to the parameters used for prophage protein clustering – i.e. bidirectional protein sequence coverage (column label 'coverage') and protein amino-acid sequence identity (row label 'identity'); outliers with $\Delta$F1 outside the interquartile range are labelled; dashed lines indicate the diagonal; Pearson correlation coefficients (r) are shown in the top-left corner of each panel. (B) Bar plots show, for each K-locus (X-axis), the top F1 score (Y-axis) across the same six clustering settings, comparing top F1-scores obtained from GWAS run on proteins from high- and low- quality prophages (orange) versus high-quality prophages only (blue). Top GWAS predictors from two prophage datasets show overall positive correlation across all clustering. The exclusion of low-quality prophages for most outliers leads to decrease in F1-score value (eg, KL127).

.
